# The complex inhibitory mechanism of glycomimetics with human heparanase

**DOI:** 10.1101/2023.01.05.522817

**Authors:** Cassidy Whitefield, Yen Vo, Brett D Schwartz, Caryn Hepburn, F. Hafna Ahmed, Hideki Onagi, Martin G. Banwell, Keats Nelms, Lara R. Malins, Colin J Jackson

## Abstract

Heparanase (HPSE) is the only mammalian endo-β-glucuronidase known to catalyse the degradation of heparan sulfate. Dysfunction of HPSE activity has been linked to several disease states, resulting in HPSE becoming the target of numerous therapeutic programs, yet no drug has passed clinical trials to date. Pentosan polysulfate sodium (PPS) is a heterogeneous FDA-approved drug for the treatment of interstitial cystitis and a known HPSE inhibitor. However, due to its heterogeneity, characterisation of its mechanism of HPSE inhibition is challenging. Here we show that inhibition of HPSE by PPS is complex, involving multiple overlapping binding events, each influenced by factors such as oligosaccharide length and inhibitor-induced changes in protein secondary structure. The present work advances our molecular understanding of the inhibition of HPSE, which will aid the development of therapeutics for the treatment of a broad range of pathologies associated with enzyme dysfunction including cancer, inflammatory disease and viral infections.

## Introduction

Heparan sulfate proteoglycans (HSPGs) are ubiquitous macromolecules associated with the cell surface and extracellular matrix (ECM) of animal tissue, where they mediate critical interactions between cells and their environment.^1^ HSPGs consist of pericellular or extracellular core proteins to which several heparan sulfate (HS) chains are covalently bound *via* oxygen-based linkages.^2^ The enormous structural variation available within HS chains allow HSPGs to interact with a wide range of proteins, thereby influencing many biological processes. These processes include ECM homeostasis, signalling, developmental patterning, cell adhesion, barrier formation and endocytosis.^3–8^ HS also regulates the activity of bioactive molecules such as growth factors, cytokines, chemokines and coagulation factors by providing low-affinity storage within the ECM.^9–12^ Cleavage of HS side chains therefore not only alters the integrity of the ECM, but also leads to the release of such HS-bound bioactive molecules.

Human heparanase (HPSE) is the only known endo-β-glucuronidase that cleaves HS side chains, producing shorter oligosaccharides.^13–16^ HPSE is a heterodimeric protein with an 8 kDa subunit comprising residues Gln36-Glu109 and a 50 kDa subunit comprising Lys159-Ile543. HPSE consists of two domains: HPSE consists of two domains: the (β/α)_8_ domain, which contains the active site, flanked by a smaller β-sandwich domain (**Figure 1A**). Baseline HPSE activity is highly regulated and is only seen at low levels in platelets, immune cells and the placenta. Increased HPSE expression is often observed in disease states, perhaps most notably in cancer and viral infections.^17,18^ The increased activity can significantly alter cell motility by weakening the structural HSPG networks within the ECM and basal membranes^19^ facilitating angiogenesis,^20^ inflammation^21^ and invasion of the surrounding tissue.^22,23^ In addition to modifying the structure of the ECM, the breakdown of HS chains by HPSE can release latent pools of growth factors.^24^ This can have the cumulative effect of altering cell proliferation, motility and activation of important intracellular signalling pathways, including those involving pro-inflammatory cytokines.^25,26^

**Figure 1.**
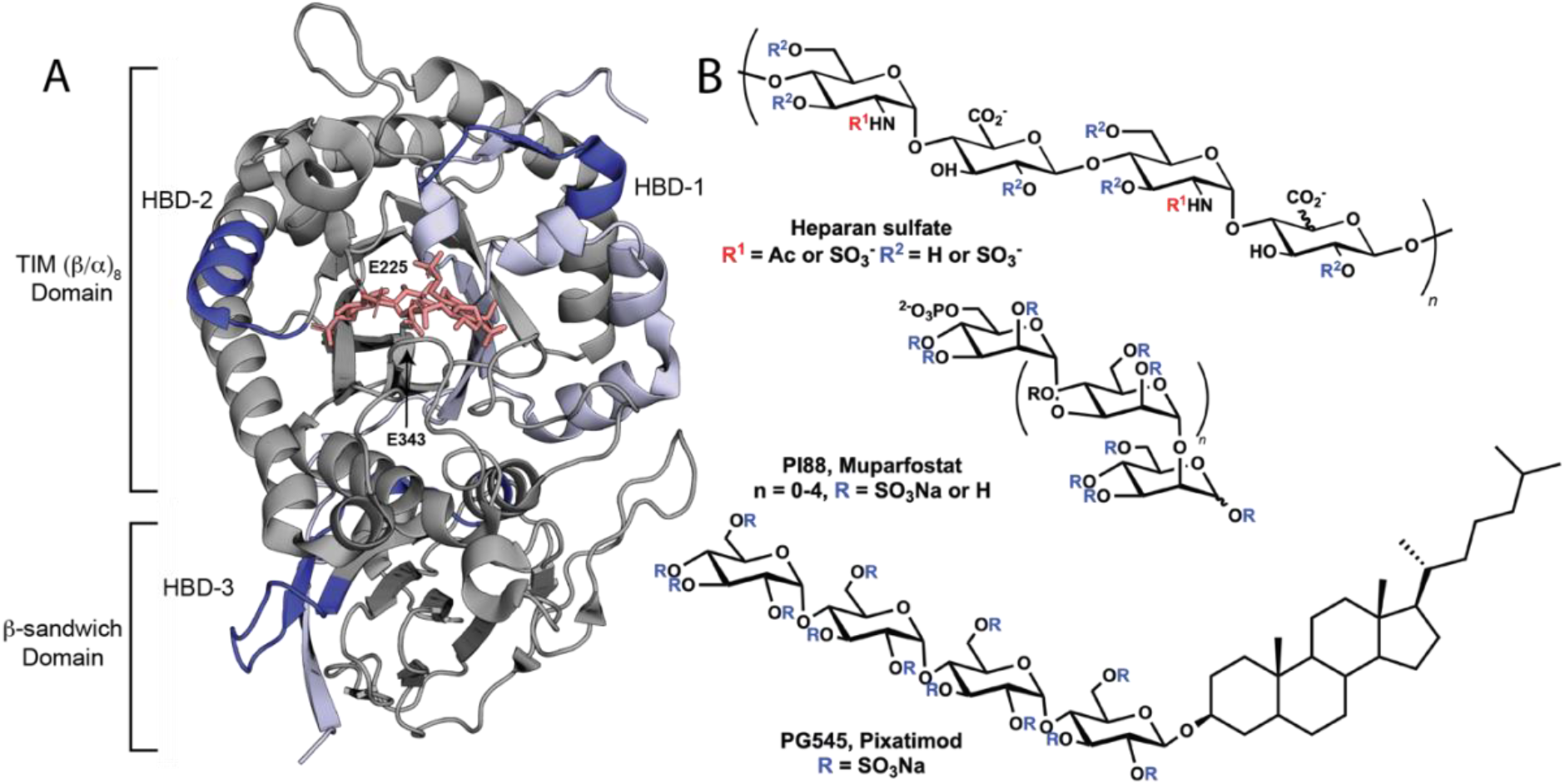
Crystal structure of HPSE and known inhibitors. **A)** Crystal structure of HPSE (PDB ID: 5E9C). Chain-A is shown in grey, and chain-B in light blue. Heparin binding domains (HBDs) 1-3 are depicted in dark blue and the catalytic residues Glu225 (acid/base) and Glu343 (nucleophile) in the enzyme binding cleft are shown in stick representation in grey. The substrate analogue is shown in pink (PDB: 5E9C). **B)** The structure of heparan sulfate (top) compared to studied HS mimetic inhibitors PI88 (middle) and PG 545 (bottom).

Due to the role HPSE plays in promoting tumour growth and metastasis, it has been a target of many therapeutic development programs. HPSE inhibitors in development include HS-glycomimetics,^27–30^ synthetically-produced small molecule compounds,^31–33^ nucleic acid-based inhibitors,^34^ covalent inhibitors,^35^ monoclonal antibodies,^36^ proteins^37^ and natural products.^38^ HS-glycomimetics and certain derivatives are the only such compounds to advance to clinical trials. These include muparfostat (PI88) (**Figure 1B**),^27^ pixatimod (PG545),^28^ roneparstat (SST0001),^29^ and necuparanib (M402),^30^ which have been shown to exert anti-cancerous and anti-metastatic effects in animal models and in early clinical trials.^39,40^ Although initially promising, clinical trials of all but PG545, which is currently in phase II trials,^41^ have either been paused or terminated.

Pentosan polysulfate (PPS; Elmiron) is an FDA-approved, semi-synthetic mixture of polysulfated xylans now deployed as an orally-administered treatment for bladder pain or discomfort associated with interstitial cystitis.^42,43^ Currently, PPS is undergoing clinical trials for the treatment of non-infectious arthritis^44^ while recent research has shown that it might also be effective against alphavirus-induced arthritis. PPS is a known cytokine-binding molecule that possesses biological neutralisation capacities and exerts broad anti-inflammatory effects.^45^ However, despite the interest in PPS and related sulfated oligosaccharides as HPSE inhibitors,^46^ the molecular basis for these inhibitory effects is not comprehensively understood.

In this work, we describe the mechanisms by which PPS analogues of different lengths inhibit HPSE. HPSE inhibition was examined in combination with X-ray protein crystallography and hydrogen/deuterium exchange mass spectrometry, leading to the identification of three oligosaccharide binding sites including a lesser-studied remote site. Enzyme kinetic and inhibition studies, alongside circular dichroism, were used to reveal the complex nature of HS-mimetic-induced inhibition of HPSE. This revealed that larger oligosaccharide molecules, including PPS, caused aggregation and loss of HPSE secondary structure, a phenomenon that emphasises the challenges confronting the clinical deployment of such compounds. This data advances our understanding of HS-glycomimetic inhibitors and will aid future drug development directed toward inhibition of the enzyme.

## Results

### Pentosan length affects HPSE inhibition

PPS is composed of a heterogeneous mixture of β-1-4 linked and sulfated xylooligosaccharides that include occasional sulfated 4-*O*-methyl-α-D-glucuronic acid residues attached to the C-2 position of the xylose backbone **(Figure 2A).** Deconvoluting the heterogeneity of PPS is therefore an essential step towards establishing those features of its constituents are the most important for activity. To investigate the molecular composition of PPS, size-exclusion chromatography was conducted, revealing twelve discrete peaks with each successive peak representing a homologue incorporating an additional sulfated xylan unit **(Figure 2B**). Each component was subjected to an HPSE inhibition assay using mammalian HPSE which revealed that the larger species were more potent **(Figure 2C)**. Oligosaccharides with nine or more sugar units proved to be better inhibitors than bulk PPS while those shorter than this displayed weaker activity. This shows that an increased oligosaccharide length leads to greater HPSE inhibition, even beyond the discrete active site binding capacity of HPSE of four oligosaccharide residues.^47^ It also reveals that most of the inhibition from heterogeneous (bulk) PPS is derived from the very large oligosaccharide species (9+ oligosaccharide units) present PPS.

**Figure 2.**
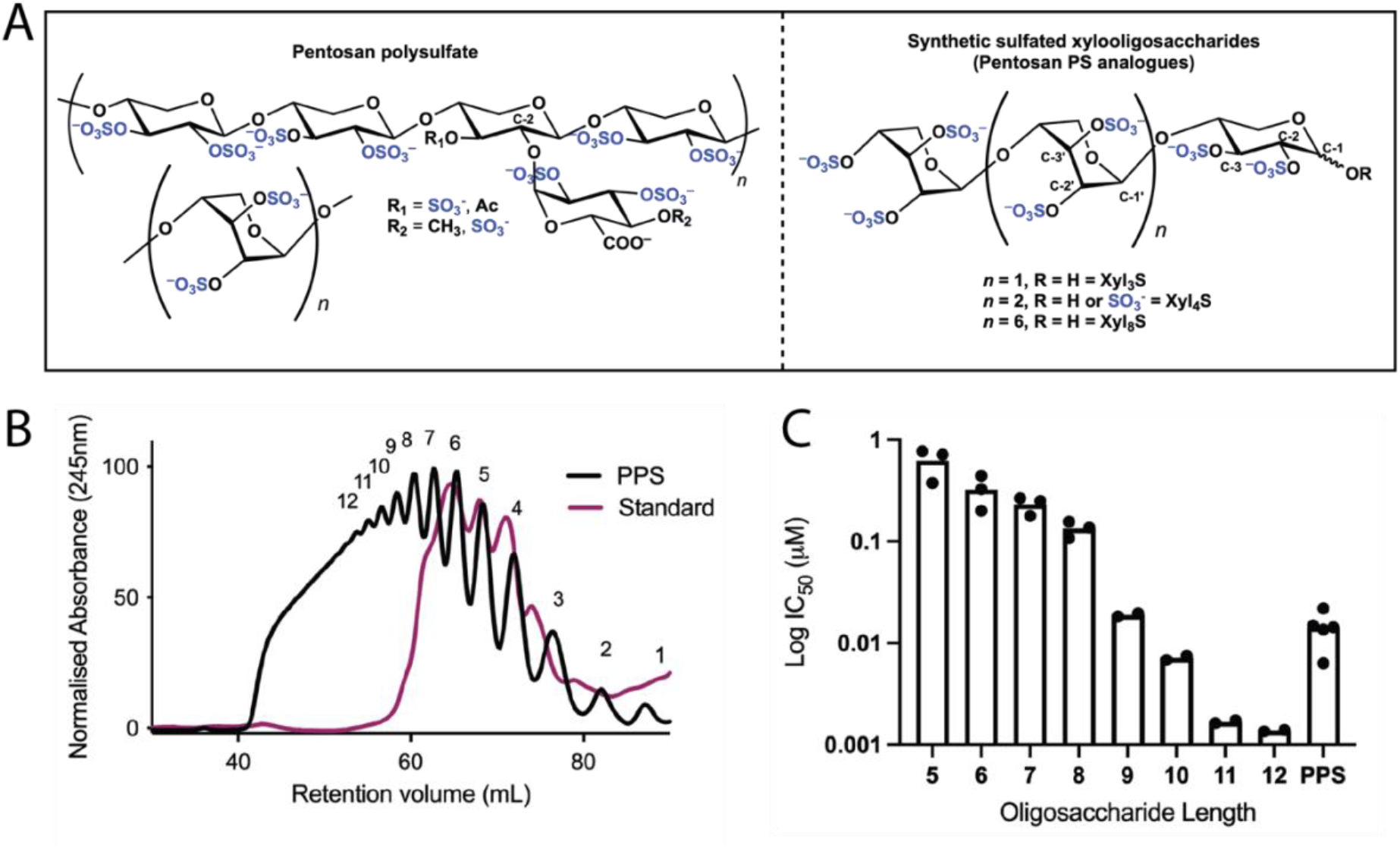
Structure, separation and activity of the individual components of PPS. **A)** Major components comprising the complex mixture of pentosan polysulfate sodium (PPS) along with synthetic analogues Xyl_3_S, Xyl_4_S and Xyl_8_S (sodium counter-ions omitted for clarity). **B)** Size-exclusion chromatography of PPS (in black) compared to an oligosaccharide standard mix (in purple).^48^ Each oligosaccharide length is highlighted. **C)** Dose-response profile (using a fondaparinux-based assay) to test the IC_50_ of each separated component vs that of PPS. All replicates shown, data presented on a log scale.

Due to the importance of PPS length in HPSE inhibition, three PPS analogues of defined structure and lacking glucuronic acid branches (side-chains) were synthesised to better understand the binding interactions.^48^ The three compounds prepared for this purpose were Xyl_3_S, Xyl_4_S and Xyl_8_S, comprising three, four and eight xylose-backbone residues, respectively. HPSE-P6 was used for these assays and subsequent studies as it has been shown to have essential identical overall structure, dynamics and activity as mammalian WT HPSE, while yielding high levels of soluble expression in *E. coli* and producing protein crystals diffract to high resolution (<2Å) within ~24h.^49^ Importantly, HPSE-P6 shows an identical inhibitory response with PPS, i.e. the conservative mutations (remote from any binding site) in HPSE-P6 do not appear to effect its function or interaction with substrate or inhibitors.^49^ Inhibition measurements revealed that PPS and the three short-chain analogues all inhibit HPSE activity in a concentration-dependant manner (**Table 1; SI Figure 1**). PPS was found to be the most active followed by Xyl_8_S, Xyl_4_S and Xyl_3_S (**Table 1**). The increased binding interactions and inhibition therefore correlates with increased oligosaccharide length. PPS, Xyl_4_S and Xyl_8_S each showed complete inhibition of HPSE at saturating levels while Xyl_3_S only inhibited up to a maximum of 66.6% of the enzyme activity at the highest concentration tested (125μM; **Table 1**) Since competitive inhibition of HPSE by HS-mimetics is typically assumed, the incomplete inhibition by Xyl_3_S is unexpected although this has been reported previously with other triose oligosaccharide inhibitors of HPSE.^50^ We assume that the shorter length of Xyl_3_S relative to the full length of the binding cleft may result in partial rather than complete blockage of substrate binding, given that the substrate analogue, fondaparinux, comprises five monosaccharide units.

**Table 1.** HPSE binding data for the three synthetic PPS analogues, including maximum inhibition and IC_50_ values (derived from inhibition curves). *K*_i_ values obtained from Lineweaver-Burk enzyme kinetics experiments. *K*_D_ values and stoichiometry of binding were obtained from ITC analysis. Errors are one standard deviation from the mean.

|  | Xyl<sub>3</sub>S | Xyl<sub>4</sub>S | Xyl<sub>8</sub>S | PPS |
| --- | --- | --- | --- | --- |
| **Max inhibition (%)** | 66.6 | 95.7 | 97.0 | 100 |
| **IC<sub>50</sub> (μM)** | 34.6 ± 4.2 | 4.01 ± 1.0 | 0.128 ± 0.006 | 0.01 ± 0.002 |
| **K<sub>i</sub> (μM)** | ND | 1.26 | 0.021 | 0.0022 |
| **K<sub>D</sub> combined (μM)** | ND | 5.44 ± 1.97 | 2.17 ± 0.552 | ND |
| **K<sub>D</sub> Site 1 (μM)** | ND | 0.430 ± 0.486 | 1.58 ± 0.21 | ND |
| **K<sub>D</sub> Site 2 (μM)** | ND | 5.87 ± 0.857 | 10.3 ± 1.21 | ND |
| **Unit of binding (N)** | ND | 2.7 | 1.8 | ND |

### Full length PPS and shorter oligosaccharides inhibit HPSE *via* different mechanisms

Given the significant differences between the inhibition of HPSE by bulk PPS and the short-chain oligosaccharides, the mechanistic basis for HPSE inhibition by the synthetically-derived compounds was investigated. Since Xyl_3_S did not result in complete inhibition and given its close similarity to Xyl_4_S, it was omitted from these experiments. Double reciprocal (Lineweaver-Burk) enzyme kinetic analyses at varying substrate and inhibitor concentrations indicated that Xyl_4_S, Xyl_8_S and PPS are all competitive inhibitors of HPSE (**Figure 3A, C, E**). However, when the gradients of the double-reciprocal graphs are plotted against inhibitor concentration, it was clear that while the responses of Xyl_4_S and Xyl_8_S were linear, long-chain PPS fits to a Hill-like model (**Figure 3F**).^51,52^ These results suggest that unlike the linear competitive inhibition seen with Xyl_4_S and Xyl_8_S, PPS is a parabolic competitive inhibitor, which has also been observed for the HPSE inhibitors PG545^50^ and SST0001.^51^ The Hill-type kinetic behaviour (**Eq.2, Methods**) displayed in the PPS inhibition of HPSE revealed a *K*_i_ of 2.2 nM and suggests that more than one inhibitor molecule is binding to the enzyme, or that PPS causes aggregation of HPSE (or both).^53,54^ In contrast, a linear competitive inhibition model (**Eq.3, Methods**) was used for Xyl_4_S and Xyl_8_S. From this analysis, *K*_i_ values for Xyl_4_S and Xyl_8_S were determined to be 1.26 μM and 21 nM, respectively. The large difference in *K_i_* values between Xyl_4_S and Xyl_8_S, relative to the number of monosaccharide residues in the two inhibitors, combined with the parabolic binding profile of PPS, suggests that the binding of these inhibitors to HPSE is a complex process.

**Figure 3.**
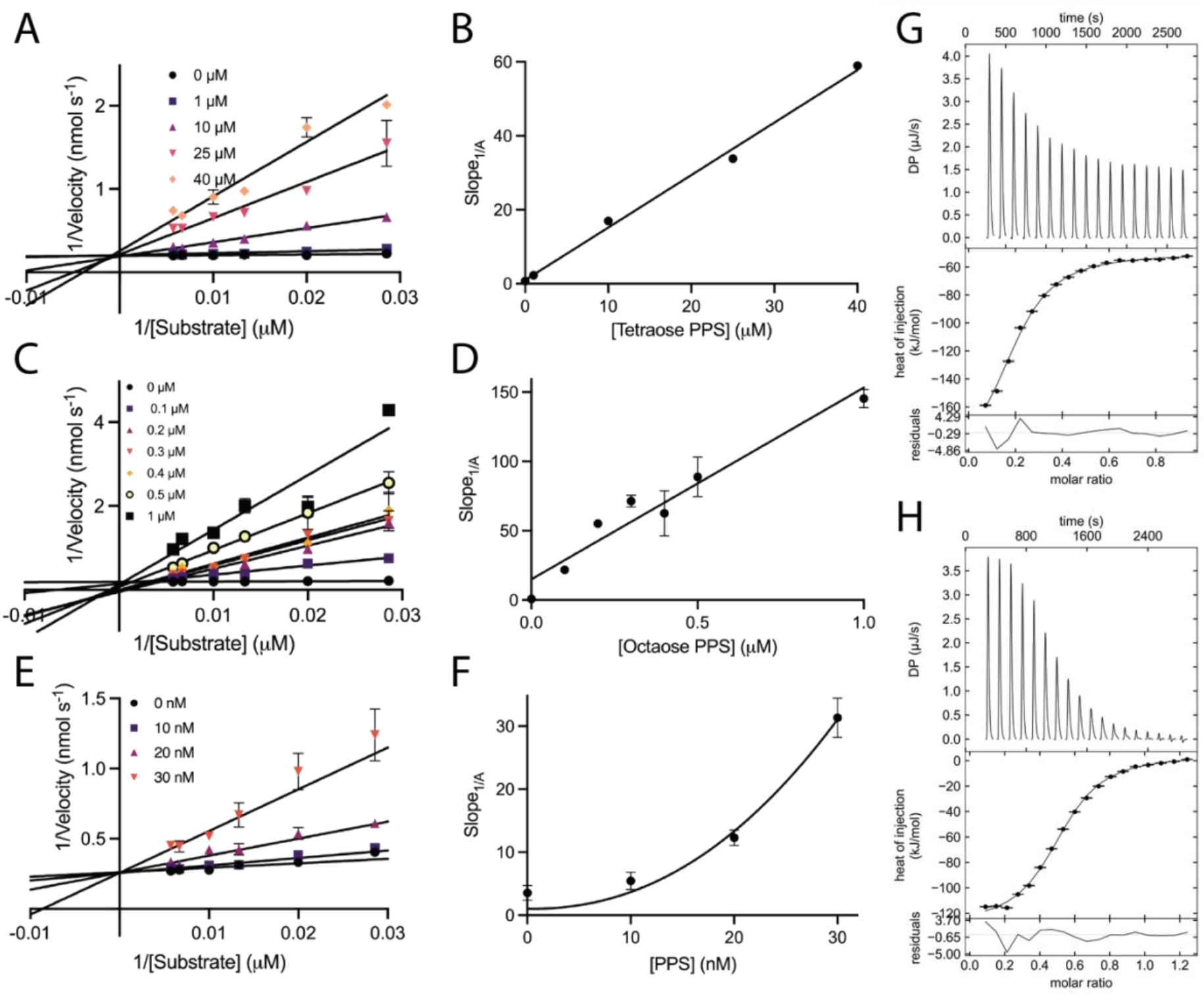
Double-reciprocal analysis of HPSE inhibition by Xyl_4_S, XyL_8_S and PPS **(A**, **C** and **E)**. Data are means of two measurements while errors are shown as standard deviation (SD). The slopes from panels A, C and E were replotted as a function of inhibitor concentration in **B**, **D** and **F**, respectively. The plots in panels **B** and **D** represent the global fit of the competitive inhibitor equation (eq. 3). The plot in panel **F** represents the fit to the Hill-type slope (eq. 2) (*R^2^* = 0.959). (**G** and **H)** ITC reverse titration data for Xyl_4_S and Xyl_8_S, respectively using the A + B ↔ AB binding model. Stoichiometry is shown as molar ratio of proteins binding to each ligand, reversed to provide ligand:protein stoichiometry.

### Thermodynamics of inhibitor binding

The binding interactions between HPSE and Xyl_4_S and Xyl_8_S were probed further using isothermal titration calorimetry (ITC). ITC allows for greater understanding of the thermodynamics of ligand binding, as well as the stoichiometries involved. Both forward and reverse titrations (ligand into protein and protein into ligand) were used to gain insights into the binding stoichiometry. Forward titrations, wherein the ligand is titrated into protein, should reveal the affinity of a ligand for an individual binding site (**SI Figure 2**), while reverse titrations, wherein protein is titrated into the ligand solution, should enable a global analysis of the affinity of the ligands for HPSE and the stoichiometry involved (**Figure 3G, H)**. Combined affinity calculations derived from the reverse ITC titrations showed that *K_D_* decreased (affinity increased) with increasing oligosaccharide length (**Table 1**) with Xyl_4_S and Xyl_8_S showing combined K_D_ values of 5.44 ± 1.97 μM and 2.17 ± 0.55 μM, respectively. These same titrations also established that Xyl_4_S and Xyl_8_S bind to HPSE with ~3:1 and ~2:1 stoichiometries, respectively (**Figure 3G, H**). Forward titration measurements with both compounds reveal that there are likely multiple binding events/sites with different affinities (**SI Figure 2**). The correlation between length and binding affinity and the magnitude of change of the *K*_D_ values for the individual sites are more consistent with the length of the oligosaccharides than the enzyme kinetic inhibition results (**Table 1**). The comparison between the *K*_i_ and *K*_D_ values for Xyl_4_S support competitive binding (*K*_i_ is of the same magnitude as *K*_D_). However, this is not the case for Xyl_8_S, for which the *K*_i_ is approximately 100-fold lower than the *K*_D_ (*K*_i_ << *K*_D_). These results suggest that the inhibition of HPSE by Xyl_8_S (and longer-chain components in PPS) is a more complex process than one involving a simple competitive mechanism. Given that ITC measurements are recorded instantaneously, while the kinetic assays take place over a longer timescale, the difference could result from the longer PPS species affecting the HPSE structure.

### Structural interactions of PPS showing multiple binding sites to HPSE

To further probe the interactions of PPS oligosaccharides with HPSE at the molecular level, we solved crystal structures of HPSE-P6 in complex with Xyl_3_S and Xyl_4_S (**SI Table 3**). Crystals of HPSE were soaked with ligand before flash-cooling and data collection. Electron density corresponding to Xyl_3_S and Xyl_4_S ligands were identified in the crystal structures. In contrast, it was not possible to obtain a complex of HPSE with Xyl_8_S, due to structural disruption of the protein (crystal deterioration and loss of diffraction). Two molecules of Xyl_3_S and Xyl_4_S were seen around the active site, at the previously described heparan binding domains (HBDs) 1 (Lys158-Lys162) and 2 (Pro271-Met278) (**Figure 4, SI Figure 3**). The electron density for some regions of the oligosaccharides were weak, indicating the binding is relatively dynamic or the ligands were bound at less than full occupancy (**Figure 4, SI Figure 3**). However, the electron density was unambiguous in certain regions and so confirming the analogues were bound at these sites. At HBD-1 there are interactions between the Xyl_3_S sulfate moieties and Asn64, Lys98, Lys159, Lys161 and Tyr391. For Xyl_4_S, further interactions are also seen with Tyr97, Lys232, and Gln383. Although the two PPS analogues bind at similar locations, Xyl_3_S faces out of the binding site towards the solvent, possibly allowing for substrate still interact with HPSE resulting in the partial inhibition observed in the assays (**Table 1**). In contrast, Xyl_4_S has more extensive interactions with HPSE and so allowing it to sterically block more of the active site, providing a structural explanation for the partial inhibition observed for Xyl_3_S (**Table 1**). In both Xyl_x_S:HPSE complexes, the HBD-1 residues of Lys159, Phe160 and Lys161 undergo a large conformational change compared to the apo and ligand bound forms of HPSE that have been observed in other crystallographic studies (PDB: 7RG8 and 5E9C). These residues rotate out towards the solvent, allowing for Xyl_x_S to bind, whereas they usually face towards the binding site, over the top of the substrate (**SI Figure 4**). These residues have been shown to play an important role in HPSE inhibition previously.^55^ At HBD-2, the binding interactions of Xyl_3_S and Xyl_4_S are very similar (**Figure 4C and SI Figure 3C**) with HBD-2 residues Asn238, Ser240, Lys274, Lys277, Met278 and Ser281 involved in binding.

**Figure 4.**
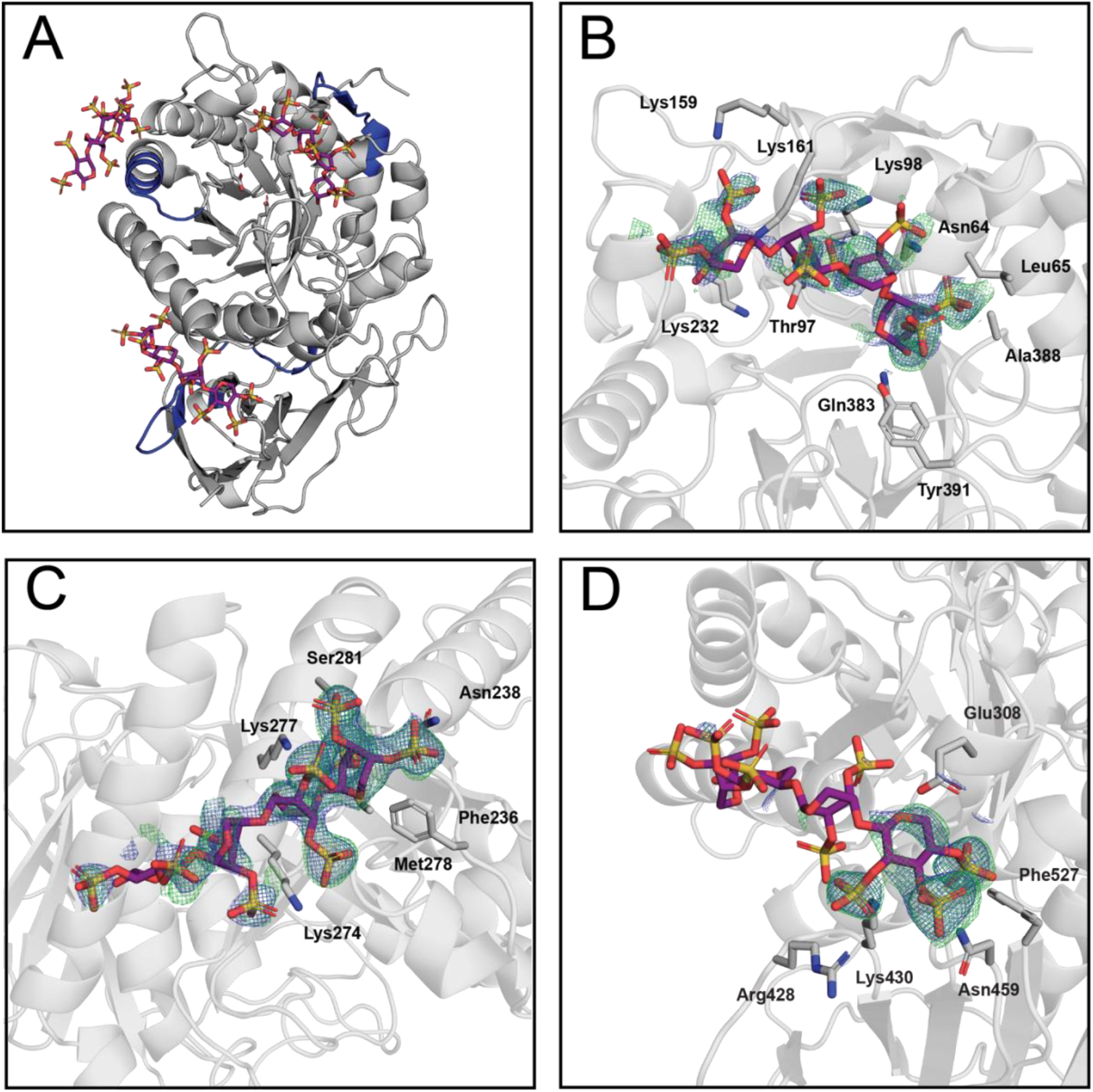
Crystal structure of the Xyl_4_S-bound HPSE (8E08). **A)** Crystal structure of Xyl_4_S bound to HPSE, with HBDs highlighted in blue. **B)** shows a zoomed in view of the Xyl_4_S ligand with density at HBD-1, **C)** HBD-2, **D)** and HBD-3. mFo-DFc maps represented in green (2.5σ) and 2mFo-DFc map in blue (1σ).

Consistent with the predicted stoichiometry from the ITC measurements (**Figure 3G, H**), a third binding site can also be discerned in the crystal structure involving Xyl_4_S (**Figure 4D**). HBD-3 was first proposed by Levy-Adam *et al*. but because of an inability to purify the properly folded protein with this domain deleted, it was suggested that this region was important for heterodimer formation and most likely did not bind to HS.^55^ Here, we observe electron density for the non-reducing end of Xyl_4_S, indicating that it is bound at the HBD-3 site (involving residues Lys411-Lys417, Lys427-Arg432)^55^ while Asn459, Phe527 and Glu308 are also show to interact. The remaining units of this oligosaccharide are not well resolved in the crystal structure, most likely due to the high conformational flexibility of the oligosaccharide in this region. Thus, a third oligosaccharide binding domain likely exists in HPSE, clarifying ambiguity in the literature surrounding this site.

The observed conformations of the sulfated xylooligosaccharides in the crystal structures are consistent with those of the isolated Xyl_2-8_S systems recently reported by Vo *et al*.^48^ Specifically, the sulfated, non-reducing xylose residues preferentially adopt a ^1^C_4_ conformation. However, when bound to HSPE, the internal sulfated xylose reside of Xyl_3_S adopts a ^4^C_1_ conformation, suggesting that hydrogen bonding to Lys98 and Lys161 stabilises this over the ^1^C_4_ conformation observed in solution.^48^ Likewise, the reducing end of Xyl_4_S adopts a ^1^C_4_ conformation and, as a result, hydrogen bonds between Tyr391 and the β-C-1 hydroxyl are observed while Asn64 and Lys98 interact with the sulfate residues at C-2 and C-3, respectively.

### Hydrogen/deuterium exchange mass spectrometry measurements and mutagenesis support multiple binding sites

Hydrogen/deuterium exchange mass spectrometry (HDX-MS) was employed to confirm that the binding of Xyl_3_S occurs in the regions suggested by X-ray protein crystallography in solution. Xyl_3_S was the only analogue studied with this method due to the high level of aggregation seen with the other analogues. In-line pepsin digestion and filtering yielded 206 peptides with 98.7% coverage of the primary sequence. In the presence of Xyl_3_S, several regions of HPSE were shielded from solvent exchange, with the most noticeable regions being residues Lys159 – Val170 (including HBD-1), Gly265 – Leu283 (including HBD-2) and to a lesser extent, Ser422 – Leu435 (including HBD-3) (**Figure 5A, SI Figure 6**). These regions of reduced deuterium incorporation are consistent with the HPSE-Xyl_4_S co-crystal structure showing binding at HBD-1 and HBD-2, and HBD-3. It is likely that Xyl_3_S was not observed at HBD-3 in the crystal structure owing to lower affinity compared with Xyl_4_S at this site. Other regions of HPSE also exhibited minor shielding from deuterium exchange, such as the region spanning residues Asn224 – Ile237 at the active site of HPSE, encapsulating one of the catalytic residues Glu225 as well as the neighbouring HBD-2 loop. Surface-exposed random coils of the β-sandwich domain also experience variations in deuterium uptake, namely, residues Tyr468 – Gln477 and Val503 – Ser521. These variations may be explained by the considerable structural dynamics of HPSE, suggesting that ligand binding could affect conformational sampling.

**Figure 5.**
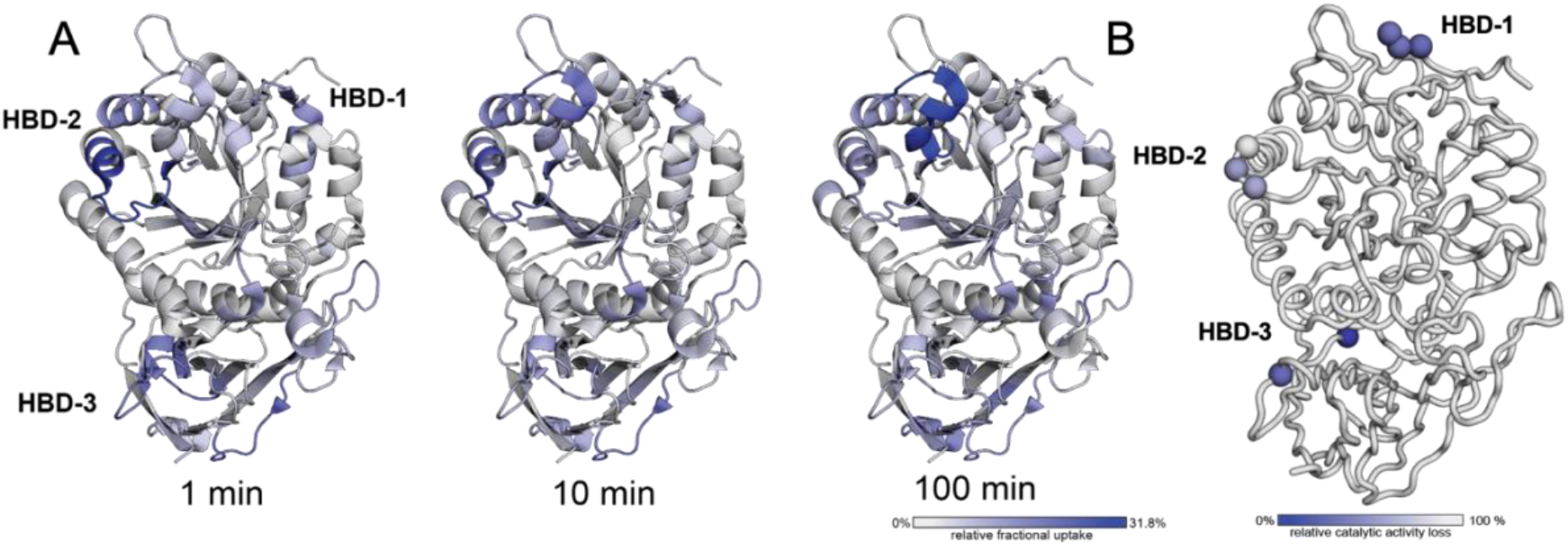
Hydrogen/deuterium-exchange mass spectrometry difference data and effect of HBD mutations. **A)** Representation of the HDX-MS difference data for apo- and Xyl_3_S-bound HPSE at 5 °C at the 1, 10 and 100 min timepoints overlaid on HPSE P6 crystal structure (7RG8). **C)** Relative catalytic activity of point mutations made on HBD1-3 compared to HPSE P6 activity.

Site-directed mutagenesis of residues at the three heparin binding domains led to a reduction in activity, confirming the importance of these regions by reducing the relative activity (**Figure 5B, SI Table 1**). All mutations localised at HBD-1 (Lys159Ala, Phe160Ala, Lys161Ala) caused a 35-40% reduction in activity and showed the importance of this domain. The HBD-2 mutation Arg272Ala caused a decrease in activity to 57%, although Lys274Ala had no effect. Lastly, HBD-3 mutations Lys417Ala and Arg428Ala resulted in a loss of activity to 17% and 35%, respectively. These results are notable in that they suggest that the remote HBD-3 site can affect catalytic activity allosterically.

### Larger oligosaccharides induce loss of secondary structure and aggregation in HPSE

The structural effects of larger oligosaccharides binding to HPSE could not be assessed using X-ray crystallography or HDX-MS techniques due to protein aggregation and structural disruption. To further investigate and to probe the link between this and the anomalous enzyme inhibition effects exerted by long-chain oligosaccharides, circular dichroism (CD), enzyme kinetic assays and non-specific aggregation assays were conducted to establish whether PPS and its analogues affect HPSE function other than by direct competitive inhibition.

CD analysis showed that 10 minutes after the addition of a 10:1 molar ratio of Xyl_3_S or Xyl_4_S with HPSE, there was no significant change in the protein secondary structure, as compared to the apo-protein. In contrast, addition of Xyl_8_S and PPS in the same molar ratio resulted in a large shift in the secondary structure distribution with loss of the α-helix troughs at 210 nm and 222 nm, as well as a shift to a minimum at 228 nm, a change which may be due to aggregation^56^ (**Figure 6A**). Notably, upon removal of Xyl_8_S and PPS through dialysis, the secondary structure remained altered (**Figure 6B**). These results are suggestive of a loss of typical secondary structure for an α/β protein (210 nm and 222 nm) and gain in CD signatures corresponding to an aberrant, and likely aggregated state. We observed the same effect on protein structure when we tested another long-chain, high-affinity oligosaccharide HPSE inhibitor, the drug candidate PI-88 (**SI Figure 5**).

**Figure 6.**
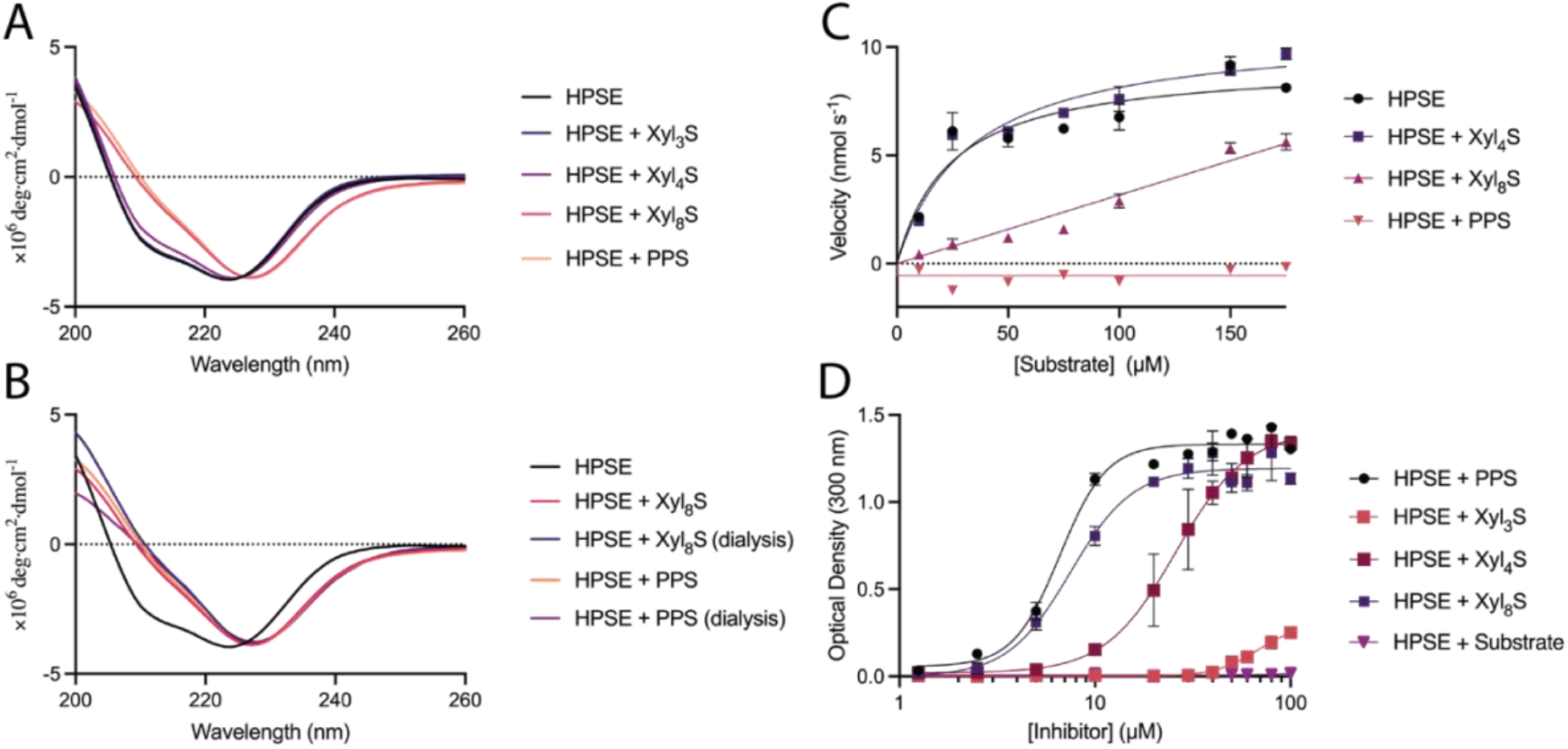
PPS-induced aggregation promotes secondary structure changes and reversibility. **A)** Circular dichroism of HPSE upon the presence of PPS and the synthetic analogues shows secondary structure is altered with longer oligosaccharide lengths. **B)** Circular dichroism of HPSE before and after dialysis showing the secondary structure change is not reversible upon attempted removal of inhibitor. **C)** Michaelis-Menten kinetics of HPSE after removal of PPS and synthetic analogues (by dialysis), showing an increased trend between length and irreversible loss of activity. **D)** Optical density assay showing macroscopic aggregation of HPSE with PPS and its synthetic analogues, as well as substrate (fondaparinux) showing that with increased PPS length, there is increased aggregation. A 1:1 molar ratio occurs at 10 μM Inhibitor concentrations.

Activity assays were conducted with the dialysed HPSE samples to understand how changes in secondary structure affected the catalytic activity. HPSE incubated with Xyl_4_S was able to regain full activity after dialysis (6 hours) to remove the bound ligand (**Figure 6C**). In contrast, Xyl_8_S-incubated HPSE only partially regained catalytic activity (15% of WT) and HPSE incubated with PPS did not regain any catalytic activity after dialysis. The observation that the Xyl_8_S-treated HPSE regains some activity while PPS effects irreversible inactivation suggests that the length and branching variations within the components of PPS could make it impossible to completely remove it from the system due to the increased complexity of the aggregation process(es) or the high affinity of the interaction.

Finally, aggregation assays were conducted to confirm aggregation of HPSE, a feature that may be contributing to long-timescale inhibitory effects. Optical density measurements were used to identify the level of inhibitor-induced aggregation of HPSE at a concentration of 10 μM (**Figure 6**). At a 1:1 stoichiometry, (10 μM concentration) PPS and Xyl_8_S caused aggregation of 80% and 70% HPSE, respectively. In contrast, at 1:1 stoichiometry Xyl_4_S resulted in just over 10% aggregation, while Xyl_3_S resulted in a very low level of aggregation even at 10:1 stoichiometry. The high levels of aggregation detailed above are consistent with the observed changes in secondary structure induced by Xyl_8_S and PPS (**Figure 6A**), where even low concentrations of Xyl_8_S and PPS affect protein structure. The results from the Xyl_4_S-based studies suggest that although high concentrations of this ligand induce aggregation, the binding interaction is weaker, not causing a change in secondary structure, and allowing aggregation to be reversible. Importantly, fondaparinux, a synthetic substrate of HPSE, does not induce aggregation (**Figure 6D**), suggesting that the HS mimetics under study here interact differently than HS or that the ability to cleave the substrate can prevent aggregation. The observation that PI-88 also results in macro-aggregation of HPSE at low stoichiometries suggests that this effect could be a general property of long-chain glycomimetic inhibitors (**SI Figure 5**).

These results explain the anomalous kinetic and thermodynamic data presented in **Figures 3** and **4**. Specifically, the parabolic inhibition of HPSE by PPS is likely due to aggregation and simultaneous binding of multiple HPSE enzymes.^53,54^ Furthermore, the 100-fold difference between the *K*_D_ and *K*_i_ for Xyl_8_S derives from a time-dependent effect: the affinity for Xyl_8_S is likely only on the order of 2 μM, but over the duration of the assay, the protein is inactivated through aggregation and structure related events, resulting in an apparent *K*_i_ of 0.02 μM.

## Discussion

HPSE has proven to be a challenging therapeutic target, with few inhibitors reaching clinical trials and no approved HPSE-specific drug despite decades of effort. The most promising candidates have been HS glycomimetics, but these often present undesired side-effects and/or limited benefits that have complicated their clinical deployment, despite potent inhibition *in vitro*. This gap between *in vitro* potency and poor clinical outcomes has been difficult to rationalize based on our current understanding of HPSE. Indeed, undesirable effects of HS-glycomimetics have been noted, such as the observation that chronic use of PPS is associated with a novel pigmentary maculopathy, that is attributed to primary retinal pigment epithelium (RPE) injury and toxicity, an effect potentially connected to the drug’s interaction with HPSE.^57,58^ Our understanding of PPS:HPSE interactions (as well as other HS-glycomimetics) has been limited by a lack of data on the binding interactions with HPSE. This study was undertaken with chemically and structurally well-defined oligosaccharide structures to further understand the binding interactions of such compounds with HPSE and help overcome the issues presented by the heterogeneity of many HS-glycomimetics.

Our analysis of the PPS binding mechanism reveals that there are various modes of HPSE inhibition, and these increase in complexity with increasing oligosaccharide length (**Figure 7**). Three potential binding sites on HPSE have been identified for PPS, overlapping with previously identified (and putative) HS binding domains. While Xyl_3_S, which cannot completely span the active site, showed incomplete competitive inhibition, Xyl_4_S behaved as a relatively straightforward competitive inhibitor (*K*_D_ ≈ *K*_i_) As the length and interactions increase, as seen with Xyl_8_S (*K*_D_ >> *K*_i_), a significant increase in protein aggregation is observed. This is attended by a loss of native secondary structure and so leading to irreversible inhibition. This effect increases with full-length PPS, which exhibits parabolic competitive inhibition, consistent with binding of multiple proteins by a single PPS molecule.

**Figure 7.**
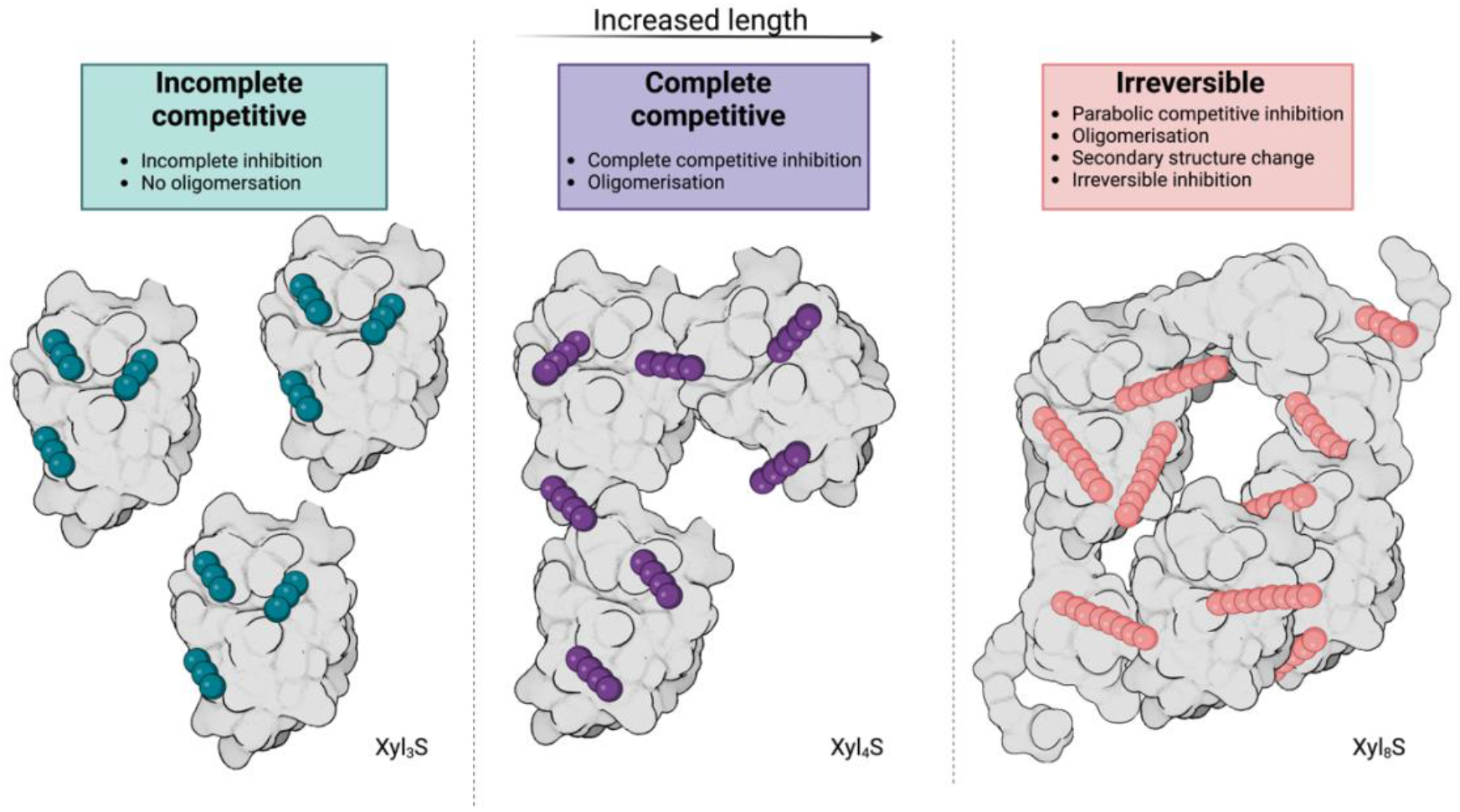
Schematic representation of the HPSE-ligand binding modes with increasing oligosaccharide length. HPSE is shown in light grey with pentosan glycol units (coloured spheres) represented in teal for Xyl_3_S, purple for Xyl_4_S and pink for Xyl_8_S.

This multiple protein-binding mode appears to be somewhat generalizable as other sulfated oligosaccharides, such as PI-88, show similar mechanisms of protein aggregation and secondary structure disruption upon incubation with HPSE. Indeed, the parabolic mode of inhibition observed here for PPS has also been observed in other HPSE inhibitors which have been deployed in clinical trials.^50,51^ For example, it has been suggested that Roneparstat (SST0001) binding could be consistent with oligomerisation of HPSE, with molecular dynamics suggesting that Roneparstat binds at the heparan binding domains surrounding the active site, orientating towards the solvent.^51^ NMR spectroscopic and MD studies suggest that Pixatimod (PG545) also undergoes multi-inhibitor binding, with two such molecules occupying the area around the HBD-1 and −2.^59^

Overall, this study demonstrates that the length of the oligosaccharides backbones in PPS plays an important role in the inhibition of HPSE and that this is not due to an increased affinity at the binding site but, rather, results from the ability to interact with numerous HPSE molecules and so indirectly affecting inhibition by deactivating HPSE through aggregation. This complex inhibitory mechanism is likely shared by other glycomimetic compounds that have entered clinical trials based on their high potency observed during *in vitro* studies. The irreversible aggregation of HPSE by these compounds could account for many of the undesirable clinical effects of these drug candidates and highlights the need for potent and specific small molecule inhibitors. This study will hopefully help other drug development programs in the future to identify the mode of inhibition seen with these types of inhibitors, which will allow for better designed drugs for this complex drug target.

## Methods

HPSE P6 was obtained following the methods by Whitefield *et al.^49^* while the chemical synthesis of Xyl_4_S is reported in detail in the supplementary material. The preparation of Xyl_3_S and Xyl_8_S have been described previously.^48^ Samples of PI-88 were donated by Professor Chris Parish of the John Curtin School of Medical Research at the Australian National University.

### Pentosan separation

Size-exclusion chromatography (Superdex S30 pg GE Healthcare) was used to separate PPS into its individual components. Thus, PPS (500 μL of 50 mg/mL) was injected onto a column equilibrated with 300 mM aqueous ammonium bicarbonate and the eluting compounds detected through their absorbance at 254 nm. Molecular weights were determined by plotting retention volumes against a standard curve for sulfated sugars of defined composition, the line of best fit being defined by;

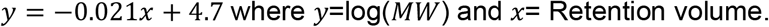

Samples were collected and freeze-dried before being reconstituted in MQ at defined concentrations.

### HPSE P6 mutagenesis

The cloning of HPSE P6 is shown in Whitefield *et al*.^49^ Genes for most HPSE P6 mutants (F159A+K160A, K161A, R272A, K274A, K411Q, G415E,) were synthesised by Twist bioscience. The original 50kDa subunit was removed and linearized using NdeI and XhoI restriction enzymes (Fast Digest, Thermo) and mutant genes were inserted into the multiple cloning site 2 by Gibson assembly.^60^ K417A and R428A mutations were introduced though PCR, amplified with mid_duet and mutagenesis primers (Table S2) and ligated with Gibson assembly. All ligated DNA was transformed in *E. coli* TOP10 cells, the plasmid DNA was extracted and sent to the Garvan Institute (Sydney) for Sanger sequencing conformation

### Protein expression and purification

HPSE P6 expression was conducted following previously described methods.^49^ HPSE P6 was transformed into *E. coli* SHuffle T7 Express cells (NEB), together with GroEL/ES + Trigger factor chaperones in a pACYC vector and spread on an Agar plate with ampicillin and chloramphenicol. 1% overnight seed culture from a single colony was inoculated into 1 L of LB medium supplemented with ampicillin (100 mg L^-1^) and chloramphenicol (34 mg L^-1^), then incubated at 37 °C for 5 hours. Overexpression was induced by adding IPTG to a final concentration of 0.05 mM and the culture was further incubated for 3 hours at 37 °C. The derived cell pellet was resuspended in buffer A (20 mM HEPES pH 8, 300 mM NaCl, 5 mM β - mercaptoethanol, 10% (v/v) glycerol, 20 mM Imidazole) with Turbonuclease (Sigma) then lysed by sonication (Omni Sonic Ruptor 400 Ultrasonic homogenizer). The lysate was filtered (0.45 μm) and loaded onto a Ni-NTA column (GE healthcare) and eluted with 100% buffer B (buffer A + 500 mM Imidazole). The peak eluent was diluted 5 times with buffer C (20 mM HEPES pH 7.4, 200 mM NaCl, 5mM β-mercaptoethanol, 10% (v/v) glycerol) and loaded onto a heparin affinity column (GE healthcare) and eluted with 100% buffer D (buffer C + 1.5 M NaCl). The peak eluent was loaded onto a size-exclusion column (HiLoad 26/600 Superdex 200 pg, GE healthcare) and eluted into buffer E (20 mM Sodium Acetate pH 5, 200 mM NaCl, 10% (v/v) glycerol, 1 mM TCEP). The final concentration of the monomeric HPSE from the gel filtration was estimated by absorbance at 280 nm using NanoDrop One (Thermo).

### HPSE activity assays

Assays were conducted using the colorimetric assay designed by Hammond *et al*.^61^ Bovine serum albumin-coated 96 well microplates were used for all assays and were prepared by incubation of the plates with 1% BSA dissolved in phosphate-buffered saline (PBS) with 0.05% Tween-20 (PBST) at 37 °C for 75 minutes. The plates were then washed three times with PBST, dried and stored at 4 °C. Assay mixtures contained 40 mM sodium acetate buffer (pH 5.0), 0.8 nM HPSE in 0.01% Tween 20 sodium acetate buffer and 100 μM fondaparinux (GlaxoSmithKline) with or without increasing concentrations of inhibitor. Plates were incubated at 37 °C for 2–20 hours before the reaction was terminated using 100 μL of 1.69 mM 4-[3-(4-iodophenyl)-2-(4-nitrophenyl)-2*H*-5-tetrazolio]-1,3-benzene disulfonate (WST-1) in 0.1 M aqueous NaOH. The plates were resealed and developed at 60 °C for 1 hour, and the absorbance was measured at 584 nm (Synergy 2). Kinetics were carried out with a standard curve constructed with *D*-galactose as the reducing sugar standard, prepared in the same buffer and volume over the range of 0–2 μM.

### Kinetic data analysis

The Hill-type model reported by Cao *et al*.^52^ was used to plot parabolic inhibition as it proved a better fit compared to the standard parabolic competitive equation^62^ and has been demonstrated for other HPSE inhibitors.

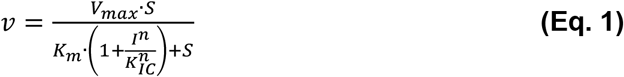

This equation can be rearranged for analysis of the slope data. (Error! Reference source not found.**.F)**

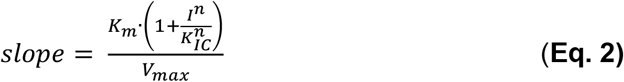

The competitive inhibition equation (**Eq. 3**) was used for Xyl_4_S and Xyl_8_S (as they exhibited the same mode of binding) and fitted to the velocity data using global nonlinear regression.

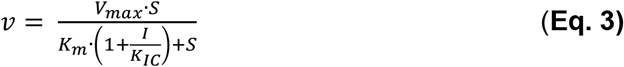

All curve fitting to calculate IC_50_ values and Michaelis–Menten constants were carried out using GraphPad Prism software (v 9.4).

### Aggregation assays

Assay mixtures contained 10 μM HPSE in buffer E and inhibitor added to a concentration of between 1.25 μM and 100 μM to a volume of 200 μL. Samples were agitated every 5 seconds between reads, for 60 minutes, and absorbance was measured at 300 nM. Negative controls were measured by testing HPSE with no inhibitor present. Curves were fit using GraphPad Prism software (v 9.4)

### Isothermal Titration Calorimetry (ITC)

ITC experiments were performed using a Nano ITC low-volume calorimeter (TA Instruments) and carried out at 25 °C with stirring at 350 rpm. Samples were prepared in SEC buffer (20 mM sodium acetate pH 5.0, 200 mM NaCl, 10% (v/v) glycerol, 1 mM TCEP). Inhibitors dissolved in MQ to 2.5 mM were diluted into SEC buffer with the same MQ dilution matched precisely to the protein buffer. Both forward and reverse titrations were undertaken, with forward titrations involving 50 μL of various inhibitor concentrations injected continuously into 500 μL of 20-25 μM protein and reverse titrations involving 50 μL of various protein concentrations injected continuously into 500 μL of 30-80 μM inhibitor over with 150 second injection intervals. NITPIC (version 1.2) was used to integrate the thermograms. The data were serially integrated and placed into a single SEDPHAT configuration file for global analysis.

The integrated ITC data were analysed using SEDPHAT (version 12.1b). The “A + B + B ↔ B + AB ↔ BA + B ↔ ABB with two non-symmetric sites, microscope K” model was used (A defined as HPSE and B as one of synthetic variants) for the forward reactions, and reverse reactions used the A + B ↔ AB model.

Since NITPIC provides error estimates for all integrated data points, the SEDPHAT option to use these as weights in the fitting sessions was activated. Switching between the Simplex and Marquardt-Levenberg optimization routines was necessary to achieve convergence of the parameter set. Standard deviations were calculated for K_D_ for both the forward and reverse reactions using GraphPad Prism software (version 9.4).

### Circular Dichroism

Circular dichroism (CD) spectra for HPSE were obtained using a 1 mm quartz cuvette and an Applied Photophysics Chirascan Spectrometer. Purified enzyme in SEC purification buffer (20 mM Sodium Acetate pH 5, 200 mM NaCl, 10% (v/v) glycerol, 1 mM TCEP) were measured at a protein concentration of 0.2 mg/mL along with the addition of inhibitors at a 1:10 molar ratio. Samples were scanned at 20 °C between 200 – 260 nm using a band width of 1 nm, and a scan rate of 1 second (with adaptive sampling enabled). Spectra were recorded in triplicate, and a buffer blank was subtracted from the results. Molar ellipticity (deg*cm^2^/dmol) was used for all subsequent analyses.

### Protein Crystallography

Well-diffracting single crystals were obtained by the hanging-drop vapor-diffusion method at 18 °C by combining the protein (6–8 mg mL-1) and the well solution [1.9 M (NH4)2SO4] in a ratio of 1.5:1.5 μL. Crystals appeared within a week and continued to grow for 1-2 months. Small amounts of ligand compound were added to the crystal drop and left to soak between 4 hrs to 2 days. Crystals were then frozen directly in liquid nitrogen without cryoprotectant. Crystallographic data were collected at 100 K at the Australian Synchrotron (MX2,^63^ 0.9537 Å). The derived diffraction data were indexed and integrated with DIALS.^64^ Resolution estimation and data truncation were performed using the AIMLESS as in implemented in CCP4^65,66^ All structures were solved by molecular replacement using the MOLREP program in CCP4^65^ using the structure deposited under PDB accession code 7RG8 as a starting model. The models were refined using phenix.refine,^67^ and the model was subsequently optimized by iterative model building with the program COOT v0.9.^68^ Alternative conformations were modelled based on mFo–DFc density and the occupancies and B-factors were determined using phenix.refine.^67^ Ligands were optimised *via* elBOW and fitted into the structure *via* LigandFit. The structures were then evaluated using MolProbity^69^ in Phenix. Details of the refinement statistics were produced by Phenix (version 1.19)^70^ and are summarized in SI Table 2. The structures were visualized and analysed using PyMol (version 2.5).^71^

### HDX-Mass Spectrometry Deuterium labelling and quench conditions

A Trajan LEAP HDX Automation manager was used to automate labelling, quenching and injection of samples. Inhibitor-bound proteins were prepared at a concentration of 12 μM with a 10x molar ratio of inhibitor. 3 μL of protein sample was incubated in 57μL sodium acetate buffer pH 5.0 reconstituted in D_2_O (99.90%, Sigma). Deuterium labelling was performed for 0.5 min, 1 min, 10 min and 100 min, followed by quenching of 50μL the deuterium exchange reaction mixture in 50μL pre-chilled 50mM sodium acetate quench solution containing 2 M guanidine hydrochloride (Sigma) and 200 mM TCEP, to lower the pH to 2.5 and lower temperature to 0.1 °C. A post-quench reaction time of 30 sec was used.

### Mass Spectrometry and peptide identification

Quenched samples (80 μL) were injected onto chilled Trajan HDX Manager. Samples were subjected to online digestion using an immobilized Waters Enzymate BEH pepsin column (2.1 × 30 mm) in 0.1% formic acid in water at 100 μL/min. The proteolyzed peptides were trapped in a 2.1 × 5 mm C18 trap (ACQUITY BEH C18 VanGuard Pre-column, 1.7 μm, Waters, Milford, MA). The proteolyzed peptides were eluted using acetonitrile and 0.1 % formic acid gradient (5 to 35 % 6 min, 35% to 40% 1min, 40% to 95% 1 min, 95% 2 min) at a flow rate of 40 μL/min using an ACQUITY UPLC BEH C18 Column (1.0 × 100 mm, 1.7 μm, Waters, Milford, MA) pumped by UPLC I-Class Binary Solvent Manager (Waters, Milford, MA). A positive electrospray ionization source fitted with a low flow probe was used to ionize peptides sprayed into a SYNAPT G2-Si mass spectrometer (Waters, Milford, MA). Data was acquired in Masslynx 4.2 in MS^E^ acquisition mode using 200pg/μL Leucine enkephalin and 100 fmol/μL [Glu1]-fibrinopeptideB ([Glu1]-Fib). Lockspray was introduced by infusion at a flow rate of 5 μL/min into the mass spectrometer. Protein Lynx Global Server (PLGS) (version 3.0) was used to identify peptides in non-deuterated protein samples. The identified peptides were further filtered in DynamX (version 3.0) using a minimum intensity cut-off of 10,000 for product and precursor ions, minimum products per amino acids of 0.3 and a precursor ion mass tolerance of 5 ppm using DynamX (version 3.0) (Waters, Milford, MA). Deuterium exchange plots, relative deuterium exchange and difference plots were generated. All deuterium exchange experiments were performed in triplicate and reported values are not corrected for deuterium back exchange.

## Acknowledgements

We acknowledge the ARC Centre of Excellence for Innovations in Peptide and Protein Science (CE200100012), the ARC Centre of Excellence in Synthetic Biology (CE200100029) and an ARC Linkage Grant (LP160101552) for funding. We thank the staff of the MX2 beamline at the Australian Synchrotron (part of ANSTO) who made use of the Australian Cancer Research Foundation (ACRF) detector. We also acknowledge the University of Melbourne, Bio21 Mass Spectrometry Proteomics Facility for use of equipment. The table of contents entry was created with BioRender.com

## Contributions

C.W., C.J., M.B, B.S., K.N, conceived the study. B.D.S. and Y.V. developed synthetic pathways and chemically synthesised the sulfated xylooligosaccharide analogues and prepared the corresponding supplementary material. C.W. and H.A. separated and tested PPS fractions. C.W. purified and crystallised protein. C.W. collected, processed and refined X-ray crystallography data with assistance from C.J. C.W. performed assays, isothermal titration calorimetry, and circular dichroism studies. C.H. collected and processed HDX-MS data. H.O. provided invaluable assistance in separation science. C.W. and C.J.J. analysed results. C.W. prepared the manuscript with input from other authors. C.J.J., M.G.B., L.M. supervised the project.

## References

(1) Sarrazin, S.; Lamanna, W. C.; Esko, J. D. Heparan Sulfate Proteoglycans. Cold Spring Harb Perspect Biol 2011, 3 (7). https://doi.org/10.1101/cshperspect.a004952.

(2) Kjellén, L.; Lindahl, U. Proteoglycans: Structures and Interactions. Annu. Rev. Biochem. 1991, 60 (1), 443–475. https://doi.org/10.1146/annurev.bi.60.070191.002303.

(3) Perrimon, N.; Bernfield, M. Specificities of Heparan Sulphate Proteoglycans in Developmental Processes. Nature 2000, 404 (6779), 725–728. https://doi.org/10.1038/35008000.

(4) Bode, L.; Murch, S.; Freeze, H. H. Heparan Sulfate Plays a Central Role in a Dynamic in Vitro Model of Protein-Losing Enteropathy. J. Biol. Chem. 2006, 281 (12), 7809–7815. https://doi.org/10.1074/jbc.M510722200.

(5) Soares, M. A.; Teixeira, F. C. O. B.; Fontes, M.; Arêas, A. L.; Leal, M. G.; Pavão, M. S. G.; Stelling, M. P. Heparan Sulfate Proteoglycans May Promote or Inhibit Cancer Progression by Interacting with Integrins and Affecting Cell Migration. BioMed Res. Int. 2015, 2015, e453801. https://doi.org/10.1155/2015/453801.

(6) Christianson, H. C.; Belting, M. Heparan Sulfate Proteoglycan as a Cell-Surface Endocytosis Receptor. Matrix Biol. 2014, 35, 51–55. https://doi.org/10.1016/j.matbio.2013.10.004.

(7) Knelson, E. H.; Nee, J. C.; Blobe, G. C. Heparan Sulfate Signaling in Cancer. Trends Biochem. Sci. 2014, 39 (6), 277–288. https://doi.org/10.1016/j.tibs.2014.03.001.

(8) Poulain, F. E.; Yost, H. J. Heparan Sulfate Proteoglycans: A Sugar Code for Vertebrate Development? Dev. Camb. Engl. 2015, 142 (20), 3456–3467. https://doi.org/10.1242/dev.098178.

(9) Yayon, A.; Klagsbrun, M.; Esko, J. D.; Leder, P.; Ornitz, D. M. Cell Surface, Heparin-like Molecules Are Required for Binding of Basic Fibroblast Growth Factor to Its High Affinity Receptor. Cell 1991, 64 (4), 841–848. https://doi.org/10.1016/0092-8674(91)90512-W.

(10) Bitan, M.; Weiss, L.; Reibstein, I.; Zeira, M.; Fellig, Y.; Slavin, S.; Zcharia, E.; Nagler, A.; Vlodavsky, I. Heparanase Upregulates Th2 Cytokines, Ameliorating Experimental Autoimmune Encephalitis. Mol. Immunol. 2010, 47 (10), 1890–1898. https://doi.org/10.1016/j.molimm.2010.03.014.

(11) Amara, A.; Lorthioir, O.; Valenzuela, A.; Magerus, A.; Thelen, M.; Montes, M.; Virelizier, J. L.; Delepierre, M.; Baleux, F.; Lortat-Jacob, H.; Arenzana-Seisdedos, F. Stromal Cell-Derived Factor-1α Associates with Heparan Sulfates through the First β-Strand of the Chemokine. J. Biol. Chem. 1999, 274 (34), 23916–23925. https://doi.org/10.1074/JBC.274.34.23916.

(12) Liu, J.; Pedersen, L. C. Anticoagulant Heparan Sulfate: Structural Specificity and Biosynthesis. Appl. Microbiol. Biotechnol. 2007, 74 (2), 263–272. https://doi.org/10.1007/s00253-006-0722-x.

(13) Vlodavsky, I.; Friedmann, Y.; Elkin, M.; Aingorn, H.; Atzmon, R.; Ishai-Michaeli, R.; Bitan, M.; Pappo, O.; Peretz, T.; Michal, I.; Spector, L.; Pecker, I. Mammalian Heparanase: Gene Cloning, Expression and Function in Tumor Progression and Metastasis. Nat. Med. 1999, 5 (7), 793–802. https://doi.org/10.1038/10518.

(14) Hulett, M. D.; Freeman, C.; Hamdorf, B. J.; Baker, R. T.; Harris, M. J.; Parish, C. R. Cloning of Mammalian Heparanase, an Important Enzyme in Tumor Invasion and Metastasis. Nat. Med. 1999, 5 (7), 803–809. https://doi.org/10.1038/10525.

(15) Kussie, P. H.; Hulmes, J. D.; Ludwig, D. L.; Patel, S.; Navarro, E. C.; Seddon, A. P.; Giorgio, N. A.; Bohlen, P. Cloning and Functional Expression of a Human Heparanase Gene. Biochem. Biophys. Res. Commun. 1999, 261 (1), 183–187. https://doi.org/10.1006/BBRC.1999.0962.

(16) Toyoshima, M.; Nakajima, M. Human Heparanase: Purification, Characterization, Cloning, and Expression. J. Biol. Chem. 1999, 274 (34), 24153–24160. https://doi.org/10.1074/JBC.274.34.24153.

(17) Arvatz, G.; Weissmann, M.; Ilan, N.; Vlodavsky, I. Heparanase and Cancer Progression: New Directions, New Promises. Hum. Vaccines Immunother. 2016, 12 (9), 2253–2256. https://doi.org/10.1080/21645515.2016.1171442.

(18) B, B.; C, Y.; A, de N.; I, G.; ML, M.-H.; I, J.; NAF, J.; N, R.; M, de G.; P, P.; M, K.; LAB, J.; T, N.; MG, N.; L, H.; FL, van de V.; R, D.; Q, de M.; J, van der V. Increased Plasma Heparanase Activity in COVID-19 Patients. Front. Immunol. 2020, 11. https://doi.org/10.3389/FIMMU.2020.575047.

(19) Bar-Ner, M.; Mayer, M.; Schirrmacher, V.; Vlodavsky, I. Involvement of Both Heparanase and Plasminogen Activator in Lymphoma Cell-Mediated Degradation of Heparan Sulfate in the Subendothelial Extracellular Matrix. J. Cell. Physiol. 1986, 128 (2), 299–306. https://doi.org/10.1002/jcp.1041280223.

(20) Elkin, M.; Ilan, N.; Ishai-Michaeli, R.; Friedmann, Y.; Papo, O.; Pecker, I.; Vlodavsky, I. Heparanase as Mediator of Angiogenesis: Mode of Action. FASEB J. Off. Publ. Fed. Am. Soc. Exp. Biol. 2001, 15 (9), 1661–1663.

(21) Digre, A.; Singh, K.; Åbrink, M.; Reijmers, R. M.; Sandler, S.; Vlodavsky, I.; Li, J.-P. Overexpression of Heparanase Enhances T Lymphocyte Activities and Intensifies the Inflammatory Response in a Model of Murine Rheumatoid Arthritis. Sci. Rep. 2017, 7 (1), 46229. https://doi.org/10.1038/srep46229.

(22) Sasaki, N.; Higashi, N.; Taka, T.; Nakajima, M.; Irimura, T. Cell Surface Localization of Heparanase on Macrophages Regulates Degradation of Extracellular Matrix Heparan Sulfate. J Immunol 2004, 172 (6), 3830–3835.

(23) Gallagher, J. T. Heparan Sulfate: Growth Control with a Restricted Sequence Menu. J Clin Invest 2001, 108 (3), 357–361. https://doi.org/10.1172/JCI13713.

(24) Ishai-Michaeli, R.; Eldor, A.; Vlodavsky, I. Heparanase Activity Expressed by Platelets, Neutrophils, and Lymphoma Cells Releases Active Fibroblast Growth Factor from Extracellular Matrix. Cell Regul. 1990, 1 (11), 833–842.

(25) Gutter-Kapon, L.; Alishekevitz, D.; Shaked, Y.; Li, J. P.; Aronheim, A.; Ilan, N.; Vlodavsky, I. Heparanase Is Required for Activation and Function of Macrophages. Proc Natl Acad Sci U A 2016, 113 (48), E7808–E7817. https://doi.org/10.1073/pnas.1611380113.

(26) Bitan, M.; Weiss, L.; Reibstein, I.; Zeira, M.; Fellig, Y.; Slavin, S.; Zcharia, E.; Nagler, A.; Vlodavsky, I. Heparanase Upregulates Th2 Cytokines, Ameliorating Experimental Autoimmune Encephalitis. Mol. Immunol. 2010, 47 (10), 1890–1898. https://doi.org/10.1016/j.molimm.2010.03.014.

(27) Parish, C. R.; Freeman, C.; Brown, K. J.; Francis, D. J.; Cowden, W. B.; Hampson, I. N. Identification of Sulfated Oligosaccharide-Based Inhibitors of Tumor Growth and Metastasis Using Novel in Vitro Assays for Angiogenesis and Heparanase Activity. Cancer Res. 1999, 59 (14), 3433–3441. https://doi.org/10.1158/0008-5472.can-03-2718.

(28) Ferro, V.; Liu, L.; Johnstone, K. D.; Wimmer, N.; Karoli, T.; Handley, P.; Rowley, J.; Dredge, K.; Li, C. P.; Hammond, E.; Davis, K.; Sarimaa, L.; Harenberg, J.; Bytheway, I. Discovery of PG545: A Highly Potent and Simultaneous Inhibitor of Angiogenesis, Tumor Growth, and Metastasis. J. Med. Chem. 2012, 55 (8), 3804–3813. https://doi.org/10.1021/jm201708h.

(29) Naggi, A.; Casu, B.; Perez, M.; Torri, G.; Cassinelli, G.; Penco, S.; Pisano, C.; Giannini, G.; Ishai-Michaeli, R.; Vlodavsky, I. Modulation of the Heparanase-Inhibiting Activity of Heparin through Selective Desulfation, Graded N-Acetylation, and Glycol Splitting. J. Biol. Chem. 2005, 280 (13), 12103–12113. https://doi.org/10.1074/jbc.M414217200.

(30) Zhou, H.; Roy, S.; Cochran, E.; Zouaoui, R.; Chu, C. L.; Duffner, J.; Zhao, G.; Smith, S.; Galcheva-Gargova, Z.; Karlgren, J.; Dussault, N.; Kwan, R. Y. Q.; Moy, E.; Barnes, M.; Long, A.; Honan, C.; Qi, Y. W.; Shriver, Z.; Ganguly, T.; Schultes, B.; Venkataraman, G.; Kishimoto, T. K. M402, a Novel Heparan Sulfate Mimetic, Targets Multiple Pathways Implicated in Tumor Progression and Metastasis. PLoS ONE 2011, 6 (6), e21106. https://doi.org/10.1371/journal.pone.0021106.

(31) Xu, Y.-J.; Miao, H.-Q.; Pan, W.; Navarro, E. C.; Tonra, J. R.; Mitelman, S.; Camara, M. M.; Deevi, D. S.; Kiselyov, A. S.; Kussie, P.; Wong, W. C.; Liu, H. N-(4-{[4-(1H-Benzoimidazol-2-Yl)-Arylamino]-Methyl}-Phenyl)-Benzamide Derivatives as Small Molecule Heparanase Inhibitors. Bioorg. Med. Chem. Lett. 2006, 16 (2), 404–408. https://doi.org/10.1016/J.BMCL.2005.09.070.

(32) Courtney, S. M.; Hay, P. A.; Buck, R. T.; Colville, C. S.; Phillips, D. J.; Scopes, D. I. C.; Pollard, F. C.; Page, M. J.; Bennett, J. M.; Hircock, M. L.; McKenzie, E. A.; Bhaman, M.; Felix, R.; Stubberfield, C. R.; Turner, P. R. Furanyl-1,3-Thiazol-2-Yl and Benzoxazol-5-Yl Acetic Acid Derivatives: Novel Classes of Heparanase Inhibitor. Bioorg. Med. Chem. Lett. 2005, 15 (9), 2295–2299. https://doi.org/10.1016/J.BMCL.2005.03.014.

(33) Madia, V. N.; Messore, A.; Pescatori, L.; Saccoliti, F.; Tudino, V.; De Leo, A.; Bortolami, M.; Scipione, L.; Costi, R.; Rivara, S.; Scalvini, L.; Mor, M.; Ferrara, F. F.; Pavoni, E.; Roscilli, G.; Cassinelli, G.; Milazzo, F. M.; Battistuzzi, G.; Di Santo, R.; Giannini, G. Novel Benzazole Derivatives Endowed with Potent Antiheparanase Activity. J. Med. Chem. 2018, 61 (15), 6918–6936. https://doi.org/10.1021/acs.jmedchem.8b00908.

(34) Simmons, S. C.; McKenzie, E. A.; Harris, L. K.; Aplin, J. D.; Brenchley, P. E.; Velasco-Garcia, M. N.; Missailidis, S. Development of Novel Single-Stranded Nucleic Acid Aptamers against the Pro-Angiogenic and Metastatic Enzyme Heparanase (HPSE1). PLoS ONE 2012, 7 (6), e37938. https://doi.org/10.1371/journal.pone.0037938.

(35) de Boer, C.; Armstrong, Z.; Lit, V. A. J.; Barash, U.; Ruijgrok, G.; Boyango, I.; Weitzenberg, M. M.; Schröder, S. P.; Sarris, A. J. C.; Meeuwenoord, N. J.; Bule, P.; Kayal, Y.; Ilan, N.; Codée, J. D. C.; Vlodavsky, I.; Overkleeft, H. S.; Davies, G. J.; Wu, L. Mechanism-Based Heparanase Inhibitors Reduce Cancer Metastasis in Vivo. Proc. Natl. Acad. Sci. 2022, 119 (31), e2203167119. https://doi.org/10.1073/pnas.2203167119.

(36) He, X.; Brenchley, P. E. C.; Jayson, G. C.; Hampson, L.; Davies, J.; Hampson, I. N. Hypoxia Increases Heparanase-Dependent Tumor Cell Invasion, Which Can Be Inhibited by Antiheparanase Antibodies. Cancer Res. 2004, 64 (11), 3928–3933. https://doi.org/10.1158/0008-5472.CAN-03-2718.

(37) Temkin, V.; Aingorn, H.; Puxeddu, I.; Goldshmidt, O.; Zcharia, E.; Gleich, G. J.; Vlodavsky, I.; Levi-Schaffer, F. Eosinophil Major Basic Protein: First Identified Natural Heparanase-Inhibiting Protein. J. Allergy Clin. Immunol. 2004, 113 (4), 703–709. https://doi.org/10.1016/j.jaci.2003.11.038.

(38) Shiozawa, H.; Takahashi, M.; Takatsu, T.; Kinoshita, T.; Tanzawa, K.; Hosoya, T.; Furuya, K.; Takahashi, S.; Furihata, K.; Seto, H. Trachyspic Acid, a New Metabolite Produced by Talaromyces Trachyspermus, That Inhibits Tumor Cell Heparanase: Taxonomy of the Producing Strain, Fermentation, Isolation, Structural Elucidation, and Biological Activity. J. Antibiot. (Tokyo) 1995, 48 (5), 357–362. https://doi.org/10.7164/antibiotics.48.357.

(39) Jia, L.; Ma, S. Recent Advances in the Discovery of Heparanase Inhibitors as Anti-Cancer Agents. Eur. J. Med. Chem. 2016, 121, 209–220. https://doi.org/10.1016/J.EJMECH.2016.05.052.

(40) Heyman, B.; Yang, Y. Mechanisms of Heparanase Inhibitors in Cancer Therapy. Exp. Hematol. 2016, 44 (11), 1002–1012. https://doi.org/10.1016/j.exphem.2016.08.006.

(41) Davar, D. Phase IIA Basket Study of Pixatimod (PG545) in Combination With Nivolumab in PD-1 Relapsed/Refractory Metastatic Melanoma and NSCLC and Pixatimod (PG545) in Combination With Nivolumab and Low-Dose Cyclophosphamide in MSS Metastatic Colorectal Carcinoma (MCRC); Clinical trial registration NCT05061017; clinicaltrials.gov, 2022. https://clinicaltrials.gov/ct2/show/NCT05061017 (accessed 2022-11-28).

(42) Schuchman, E. H.; Ge, Y.; Lai, A.; Borisov, Y.; Faillace, M.; Eliyahu, E.; He, X.; Iatridis, J.; Vlassara, H.; Striker, G.; Simonaro, C. M. Pentosan Polysulfate: A Novel Therapy for the Mucopolysaccharidoses. PLoS One 2013, 8 (1), e54459. https://doi.org/10.1371/journal.pone.0054459.

(43) Anger, J. T.; Zabihi, N.; Clemens, J. Q.; Payne, C. K.; Saigal, C. S.; Rodriguez, L. V. Treatment Choice, Duration, and Cost in Patients with Interstitial Cystitis and Painful Bladder Syndrome. Int Urogynecol J 2011, 22 (4), 395–400. https://doi.org/10.1007/s00192-010-1252-8.

(44) Kongtawelert, P.; Brooks, P. M.; Ghosh, P. Pentosan Polysulfate (Cartrophen) Prevents the Hydrocortisone Induced Loss of Hyaluronic Acid and Proteoglycans from Cartilage of Rabbit Joints as Well as Normalizes the Keratan Sulfate Levels in Their Serum. J. Rheumatol. 1989, 16 (11), 1454–1459.

(45) Sanden, C.; Mori, M.; Jogdand, P.; Jönsson, J.; Krishnan, R.; Wang, X.; Erjefält, J. S. Broad Th2 Neutralization and Anti-Inflammatory Action of Pentosan Polysulfate Sodium in Experimental Allergic Rhinitis. Immun. Inflamm. Dis. 2017. https://doi.org/10.1002/iid3.164.

(46) Parish, C. R.; Coombe, D. R.; Jakobsen, K. B.; Bennett, F. A.; Underwood, P. A. Evidence That Sulphated Polysaccharides Inhibit Tumour Metastasis by Blocking Tumour-Cell-Derived Heparanases. Int. J. Cancer 1987, 40 (4), 511–518. https://doi.org/10.1002/ijc.2910400414.

(47) Wu, L.; Viola, C. M.; Brzozowski, A. M.; Davies, G. J. Structural Characterization of Human Heparanase Reveals Insights into Substrate Recognition. Nat. Struct. Mol. Biol. 2015, 22 (12), 1016–1022. https://doi.org/10.1038/nsmb.3136.

(48) Vo, Y.; Schwartz, B. D.; Onagi, H.; Ward, J. S.; Gardiner, M. G.; Banwell, M. G.; Nelms, K.; Malins, L. R. A Rapid and Mild Sulfation Strategy Reveals Conformational Preferences in Therapeutically Relevant Sulfated Xylooligosaccharides. Chem. – Eur. J. 2021, 27 (38), 9830–9838. https://doi.org/10.1002/chem.202100527.

(49) Whitefield, C.; Hong, N.; Mitchell, J. A.; Jackson, C. J. Computational Design and Experimental Characterisation of a Stable Human Heparanase Variant. RSC Chem. Biol. 2022, 3 (3), 341–349. https://doi.org/10.1039/D1CB00239B.

(50) Hammond, E.; Handley, P.; Dredge, K.; Bytheway, I. Mechanisms of Heparanase Inhibition by the Heparan Sulfate Mimetic PG545 and Three Structural Analogues. FEBS Open Bio 2013, 3 (1), 346–351. https://doi.org/10.1016/j.fob.2013.07.007.

(51) Pala, D.; Rivara, S.; Mor, M.; Milazzo, F. M.; Roscilli, G.; Pavoni, E.; Giannini, G. Kinetic Analysis and Molecular Modeling of the Inhibition Mechanism of Roneparstat (SST0001) on Human Heparanase. Glycobiology 2016, 26 (6), 640–654. https://doi.org/10.1093/glycob/cww003.

(52) Cao, R.; Zeidan, A. A.; Rådström, P.; van Niel, E. W. J. Inhibition Kinetics of Catabolic Dehydrogenases by Elevated Moieties of ATP and ADP – Implication for a New Regulation Mechanism in Lactococcus Lactis. FEBS J. 2010, 277 (8), 1843–1852. https://doi.org/10.1111/j.1742-4658.2010.07601.x.

(53) Acker, M. G.; Auld, D. S. Considerations for the Design and Reporting of Enzyme Assays in High-Throughput Screening Applications. Perspect. Sci. 2014, 1 (1), 56–73. https://doi.org/10.1016/j.pisc.2013.12.001.

(54) Prinz, H.; Schönichen, A. Transient Binding Patches: A Plausible Concept for Drug Binding. J. Chem. Biol. 2008, 1 (1–4), 95–104. https://doi.org/10.1007/s12154-008-0011-5.

(55) Levy-Adam, F.; Abboud-Jarrous, G.; Guerrini, M.; Beccati, D.; Vlodavsky, I.; Ilan, N. Identification and Characterization of Heparin/Heparan Sulfate Binding Domains of the Endoglycosidase Heparanase. J. Biol. Chem. 2005, 280 (21), 20457–20466. https://doi.org/10.1074/jbc.M414546200.

(56) Mu, X.; Eckes, K. M.; Nguyen, M. M.; Suggs, L. J.; Ren, P. Experimental and Computational Studies Reveal an Alternative Supramolecular Structure for Fmoc-Dipeptide Self-Assembly. Biomacromolecules 2012, 13 (11), 3562–3571. https://doi.org/10.1021/bm301007r.

(57) Van Bergen, T.; Etienne, I.; Jia, J.; Li, J.-P.; Vlodavsky, I.; Stitt, A.; Vermassen, E.; Feyen, J. H. M. Heparanase Deficiency Is Associated with Disruption, Detachment, and Folding of the Retinal Pigment Epithelium. Curr. Eye Res. 2021, 46 (8), 1166–1170. https://doi.org/10.1080/02713683.2020.1862239.

(58) Pearce, W. A.; Chen, R.; Jain, N. Pigmentary Maculopathy Associated with Chronic Exposure to Pentosan Polysulfate Sodium. Ophthalmology 2018, 125 (11), 1793–1802. https://doi.org/10.1016/j.ophtha.2018.04.026.

(59) Chhabra, M.; Wilson, J. C.; Wu, L.; Davies, G. J.; Gandhi, N. S.; Ferro, V. Structural Insights into Pixatimod (PG545) Inhibition of Heparanase, a Key Enzyme in Cancer and Viral Infections. Chem. – Eur. J. 2022, 28 (11), e202104222. https://doi.org/10.1002/chem.202104222.

(60) Gibson, D. G. Chapter Fifteen - Enzymatic Assembly of Overlapping DNA Fragments. In Synthetic Biology, Part B; Voigt, C. B. T.-M. in E., Ed.; Academic Press, 2011; Vol. 498, pp 349–361. https://doi.org/10.1016/B978-0-12-385120-8.00015-2.

(61) Hammond, E.; Li, C. P.; Ferro, V. Development of a Colorimetric Assay for Heparanase Activity Suitable for Kinetic Analysis and Inhibitor Screening. Anal. Biochem. 2010, 396 (1), 112–116. https://doi.org/10.1016/j.ab.2009.09.007.

(62) Hyperbolic and Parabolic Inhibition. In Comprehensive Enzyme Kinetics; Leskovac, V., Ed.; Springer US: Boston, MA, 2003; pp 95–110. https://doi.org/10.1007/0-306-48390-4_6.

(63) Aragão, D.; Aishima, J.; Cherukuvada, H.; Clarken, R.; Clift, M.; Cowieson, N. P.; Ericsson, D. J.; Gee, C. L.; Macedo, S.; Mudie, N.; Panjikar, S.; Price, J. R.; Riboldi-Tunnicliffe, A.; Rostan, R.; Williamson, R.; Caradoc-Davies, T. T. MX2: A High-Flux Undulator Microfocus Beamline Serving Both the Chemical and Macromolecular Crystallography Communities at the Australian Synchrotron. Urnissn1600-5775 2018, 25 (3), 885–891. https://doi.org/10.1107/S1600577518003120.

(64) Winter, G.; Waterman, D. G.; Parkhurst, J. M.; Brewster, A. S.; Gildea, R. J.; Gerstel, M.; Fuentes-Montero, L.; Vollmar, M.; Michels-Clark, T.; Young, I. D.; Sauter, N. K.; Evans, G. DIALS: Implementation and Evaluation of a New Integration Package. Acta Crystallogr. Sect. Struct. Biol. 2018, 74 (2), 85–97. https://doi.org/10.1107/S2059798317017235.

(65) Winn, M. D.; Ballard, C. C.; Cowtan, K. D.; Dodson, E. J.; Emsley, P.; Evans, P. R.; Keegan, R. M.; Krissinel, E. B.; Leslie, A. G. W.; McCoy, A.; McNicholas, S. J.; Murshudov, G. N.; Pannu, N. S.; Potterton, E. A.; Powell, H. R.; Read, R. J.; Vagin, A.; Wilson, K. S. Overview of the CCP4 Suite and Current Developments. Acta Crystallographica Section D: Biological Crystallography, 2011, 67, 235–242. https://doi.org/10.1107/S0907444910045749.

(66) Karplus, P. A.; Diederichs, K. Linking Crystallographic Model and Data Quality. Science 2012, 336 (6084), 1030–1033. https://doi.org/10.1126/science.1218231.

(67) Afonine, P. V.; Grosse-Kunstleve, R. W.; Echols, N.; Headd, J. J.; Moriarty, N. W.; Mustyakimov, M.; Terwilliger, T. C.; Urzhumtsev, A.; Zwart, P. H.; Adams, P. D. Towards Automated Crystallographic Structure Refinement with Phenix.Refine. Acta Crystallogr. D Biol. Crystallogr. 2012, 68 (4), 352–367. https://doi.org/10.1107/S0907444912001308.

(68) Emsley, P.; Cowtan, K. Coot: Model-Building Tools for Molecular Graphics. Acta Crystallogr. D Biol. Crystallogr. 2004, 60 (12 I), 2126–2132. https://doi.org/10.1107/S0907444904019158.

(69) Chen, V. B.; Arendall, W. B.; Headd, J. J.; Keedy, D. A.; Immormino, R. M.; Kapral, G. J.; Murray, L. W.; Richardson, J. S.; Richardson, D. C. MolProbity: All-Atom Structure Validation for Macromolecular Crystallography. Acta Crystallogr. D Biol. Crystallogr. 2010, 66 (1), 12–21. https://doi.org/10.1107/S0907444909042073.

(70) Afonine, P. V.; Grosse-Kunstleve, R. W.; Chen, V. B.; Headd, J. J.; Moriarty, N. W.; Richardson, J. S.; Richardson, D. C.; Urzhumtsev, A.; Zwart, P. H.; Adams, P. D. Phenix.Model-vs-Data: A High-Level Tool for the Calculation of Crystallographic Model and Data Statistics. J. Appl. Crystallogr. 2010, 43 (4), 669–676. https://doi.org/10.1107/S0021889810015608.

(71) DeLano, W. L. The PyMOL Molecular Graphics System. DeLano Sci. 2002.

