## Supplementary file for "The complex inhibitory mechanism of glycomimetics with human heparanase" (from another version of this preprint: version 1, 2023-01-05)

**Table of Contents**


| | |
|---|---|
| **Inhibition Dose Response Curves for Pentosan and Synthetic Analogues** | **S-2** |
| **Isothermal Titration Calorimetry for Pentosan Analogues** | **S-3** |
| **Crystal Structure of of $Xyl_3S$-bound HPSE** | **S-4** |
| **Shift of HBD-1 Residues due to Binding of PPS-type Inhibitors** | **S-5** |
| **Effects of Mutations** | **S-6** |
| **Mutagenesis Primers** | **S-7** |
| **Aggregation Results** | **S-8** |
| **Single Crystal X-ray Data and Refinement Statistics** | **S-9** |
| **Hydrogen/Deuterium Exchange Heatmap** | **S-10** |
| **Hydrogen/Deuterium Exchange Data** | **S-11** |
| **General Synthesis Methods** | **S-17** |
| **Specific Transformations** | **S-18** |
| **References** | **S-23** |
| **$^{1}H$, $^{13}C$ and/or 2D NMR Spectra of Compounds** | **S-24** |


*All sulfated compounds are sodiated but the sodium counter-ions have been omitted for clarity.*

### Inhibition Dose Response Curves for Pentosan and Synthetic Analogues

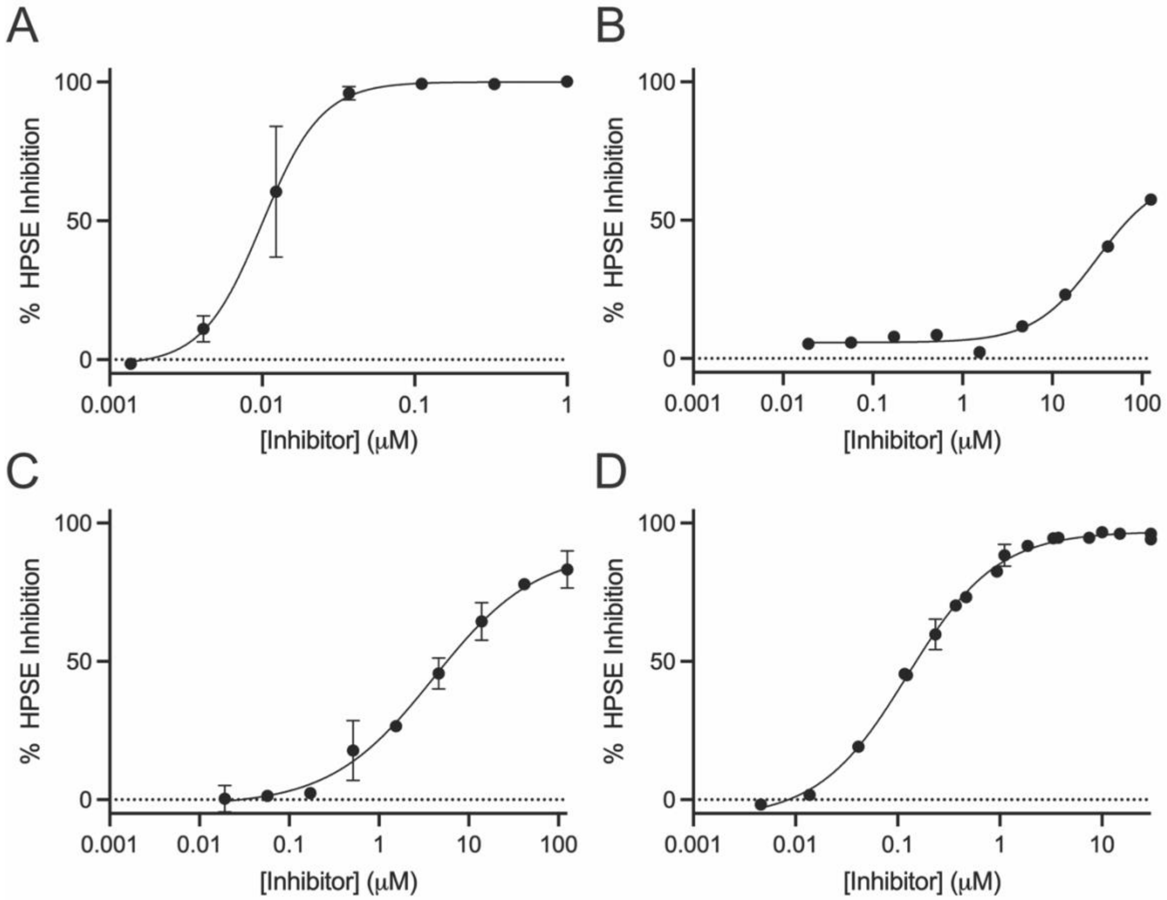


**Figure S1 Inhibition dose response curves for Pentosan and its synthetic analogues. A)** Pentosan dose response curve. **B)** Xyl$_3$S dose response curve. **C)** Xyl$_4$S dose response curve. **D)** Xyl$_8$S dose response curve. Error bars show standard error of 2-4 replicates

### Isothermal Titration Calorimetry for Pentosan Analogues

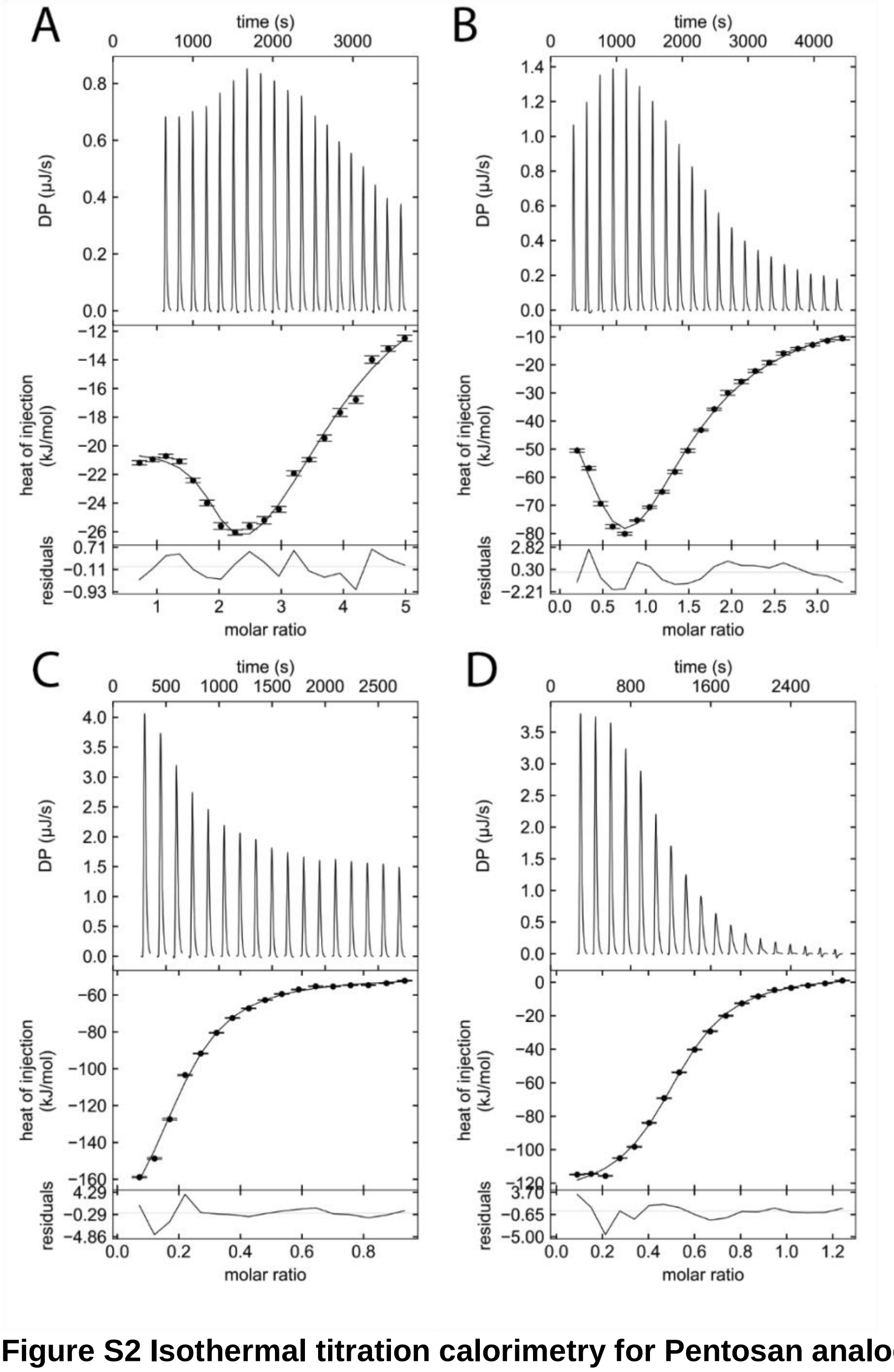


**Figure S2 Isothermal titration calorimetry for Pentosan analogues.** A-B showing a single forward titration of $Xyl_4S$ and $Xyl_8S$. C-D show a single reverse titration for $Xyl_4S$ and $Xyl_8S$, respectively.

**Crystal Structure of Xyl$_3$S-bound HPSE**

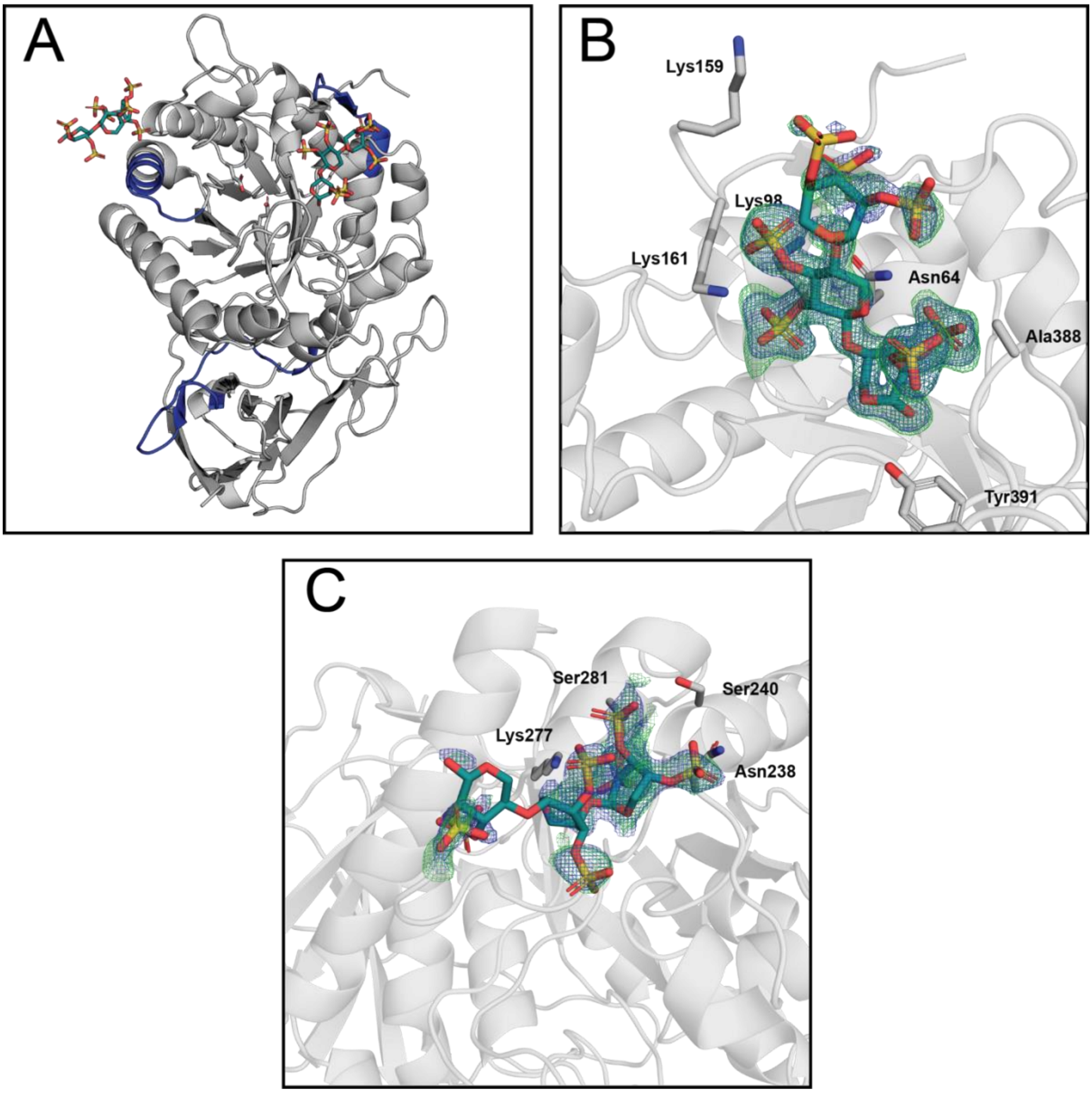


**Figure S3 Crystal structure of the Xyl$_4$S-bound HPSE (8E07). A)** Crystal structure of Xyl$_3$S bound to HPSE, with HBDs highlighted in blue **B)** shows a zoomed in view of the Xyl$_3$S ligand with density at HBD-1, **C)** HBD-2. Fo-Fc maps represented in green (2.5$\sigma$) and the 2Fo-Fc map in blue (1$\sigma$).

**Shift of HBD-1 Residues due to Binding of PPS-type Inhibitors**

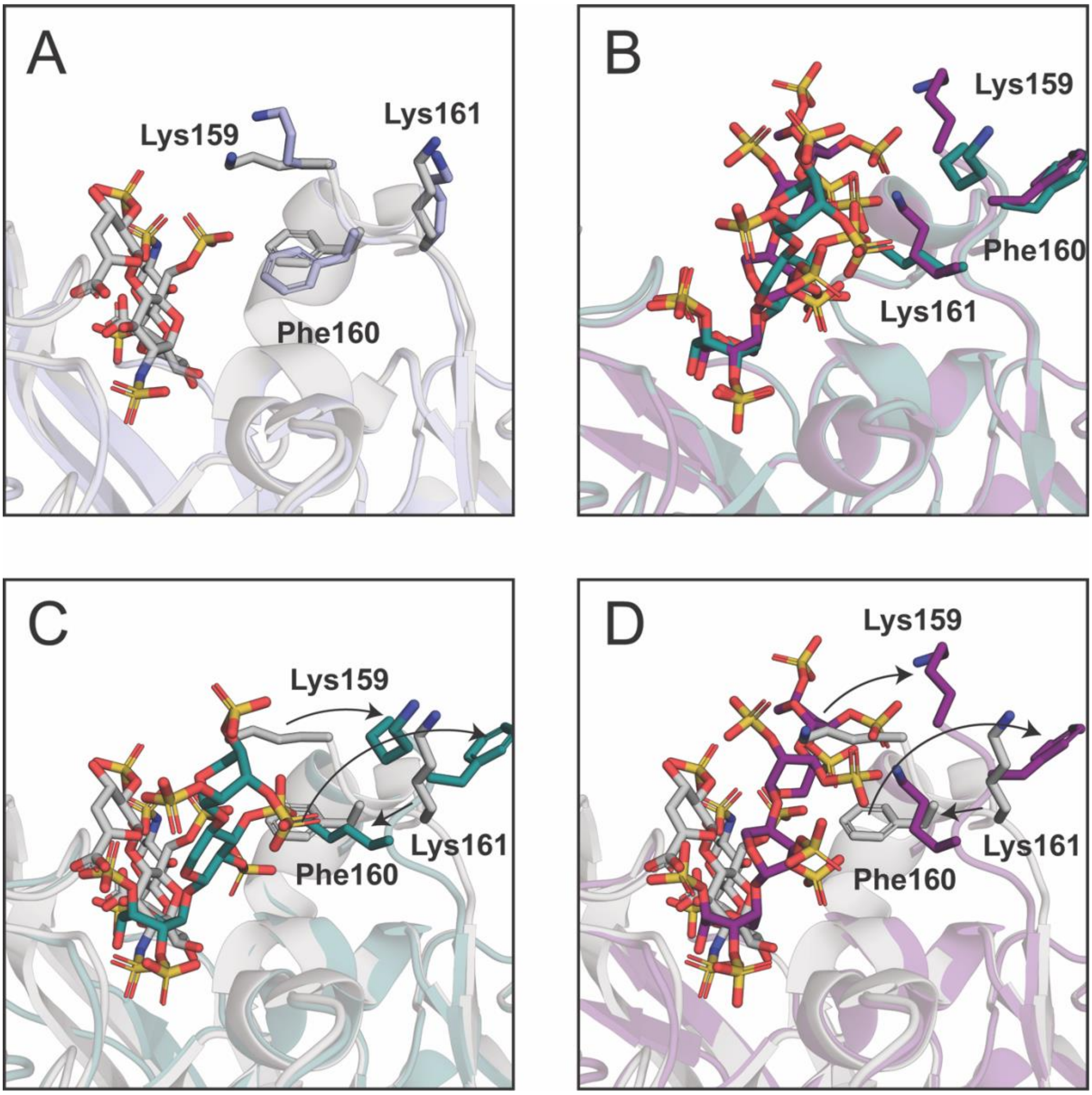


**Figure S4.** Shift of HBD-1 residues due to binding of PPS-type inhibitors. A) binding of the substrate analogue DP4 in grey (PDB: 5E9C) compared to apo form in light blue (PDB: 7RG8) with residues of HBD-1 highlighted. B) Binding of $Xyl_3SO_4$ in teal (PDB : 8E07) and $Xyl_4S$ in purple (PDB: 8E08) showing residues of HBD-1 highlighted. C) Binding of $Xyl_3S$ in teal and D) of $Xyl_4S$ in purple, compared to the substrate bound form of DP4, showing large shifts of HBD-1 residue conformation.

**Effects of Mutations**

**Table 1:** Effects of mutations upon residues within the heparin binding domains of HPSE. Errors provided are one standard deviation from the mean.

| Binding domain | Mutation | Max Activity (%) relative to HPSE P6 |
|---|---|---|
| HBD-1 | K159A, F160A | 39.36 ± 3.45 |
| | K161A | 35.66 ± 1.12 |
| HBD-2 | R272A | 56.64 ± 1.57 |
| | K274A | 100.69 ± 2.19 |
| HBD-3 | K417A | 17.05 ± 1.4 |
| | R428A | 35.19 ± 1.82 |

### Mutagenesis Primers

**Table 2:** Primers for HPSE P6 point mutations.

| Name | Sequence |
|---|---|
| K417A_F | CCTCCTTTTCAAAAAGTTAGTTGGTACAGCGGTGTTGATGGCAAGTG |
| K417A_R | CCCTGAACACTTGCCATCAACACCGCTGTACCAACTAACTTTTTG |
| R428A_F | GTGTTCAGGGTCAGGATGCGCGCAAACTTCGC |
| R428A_R | GATAAACGCGAAGTTTGCGCGCATCCTGACCCTGAAC |
| Mid_Duet_F | CCTTTTGCTGGCCTTTTGCTCACATGTTCTTTCCTGC |
| Mid_Duet_R | GCAGGAAAGAACATGTGAGCAAAAGGCCAGCAAAAGG |

### Aggregation Results

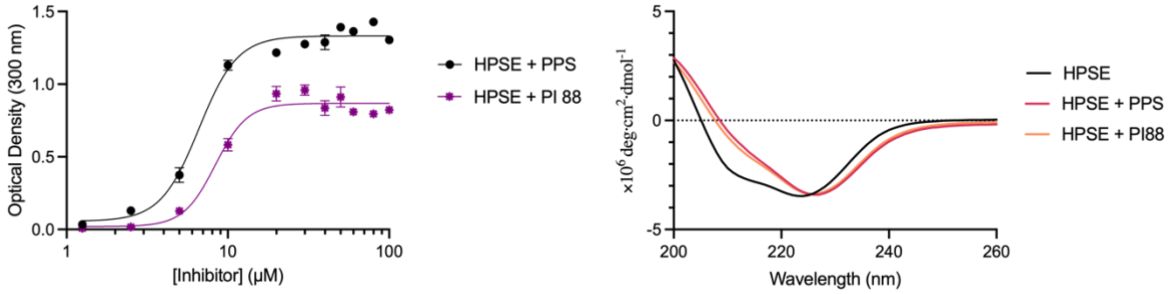


**Figure S5.** Aggregation results of other heparan sulfate mimetic inhibitors. **A)** aggregation response curves of PI-88 and PPS. **B)** Circular dichroism of HPSE in the presence of PPS and another HS-mimetic, PI-88, demonstrating that secondary structure is altered by such mimetics.

### Single crystal X-ray Data and Refinement Statistics

**Supplementary Table 3**. Data collection and refinement statistics

| PDB ID | 8E07 | 8E08 |
|---|---|---|
| **Data collection** | | |
| Space group | P $2_1$ $2_1$ $2_1$ | P $2_1$ $2_1$ $2_1$ |
| Cell dimensions | | |
| *a*, *b*, *c* (Å) | 59.54 76.19 124.97 | 59.27 75.63 125.47 |
| α, β, γ (°) | 90.00 90.00 90.00 | 90.00 90.00 90.00 |
| Resolution (Å) | 43.1 - 1.80 (1.864 - 1.80) | 46.65 - 1.93 (1.99 - 1.93) |
| $R_{merge}$ | 0.1591 (3.065) | 0.1298 (0.7661) |
| $I$ / σ$I$ | 12.05 (0.90) | 9.15 (0.80) |
| $CC_{1/2}$ | 0.998 (0.953) | 0.997 (0.919) |
| Completeness (%) | 99.41 (98.52) | 99.78 (99.91) |
| Redundancy | | 13.1 (13.7) |
| **Refinement** | | |
| Resolution (Å) | | |
| No. reflections | 53147 (5206) | 43064 (4214) |
| $R_{work}$ / $R_{free}$ | 0.172/0.203 | 0.173/0.234 |
| No. atoms | | |
| Protein | 3678 | 3636 |
| Ligand/ion | 145 | 318 |
| Water | 368 | 310 |
| *B*-factors | | |
| Protein | 33.11 | 33.86 |
| Ligand/ion | 66.51 | 82.76 |
| Water | 43.19 | 42.61 |
| R.m.s. deviations | | |
| Bond lengths (Å) | 0.006 | 0.007 |
| Bond angles (°) | 0.83 | 0.91 |
| Ramachandran plot | | |
| Preferred (%) | 98.02 | 98.24 |
| Allowed (%) | 1.98 | 1.54 |
| Outliers (%) | 0.00 | 0.22 |

*X-ray data were collected from single crystals. *Values in parentheses are for highest-resolution shell.

**Hydrogen/Deuterium Exchange Heatmap**

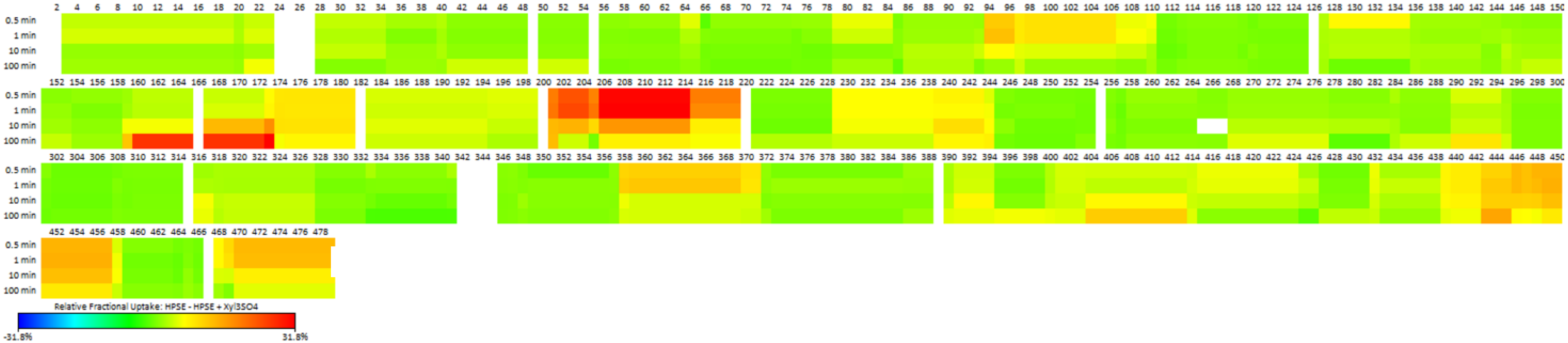


**Figure S6.** Hydrogen/deuterium-exchange mass spectrometry heat map representing the relative fractional uptake of individual peptides of HPSE relative to $Xyl_3S$ after 0.5-, 1-, 10-, and 100-minute incubation. Numbering of 1 starts from his-tagged 8 kDa subunit, with the 50 kDa subunit starting at 93.

### Hydrogen/Deuterium Exchange Data

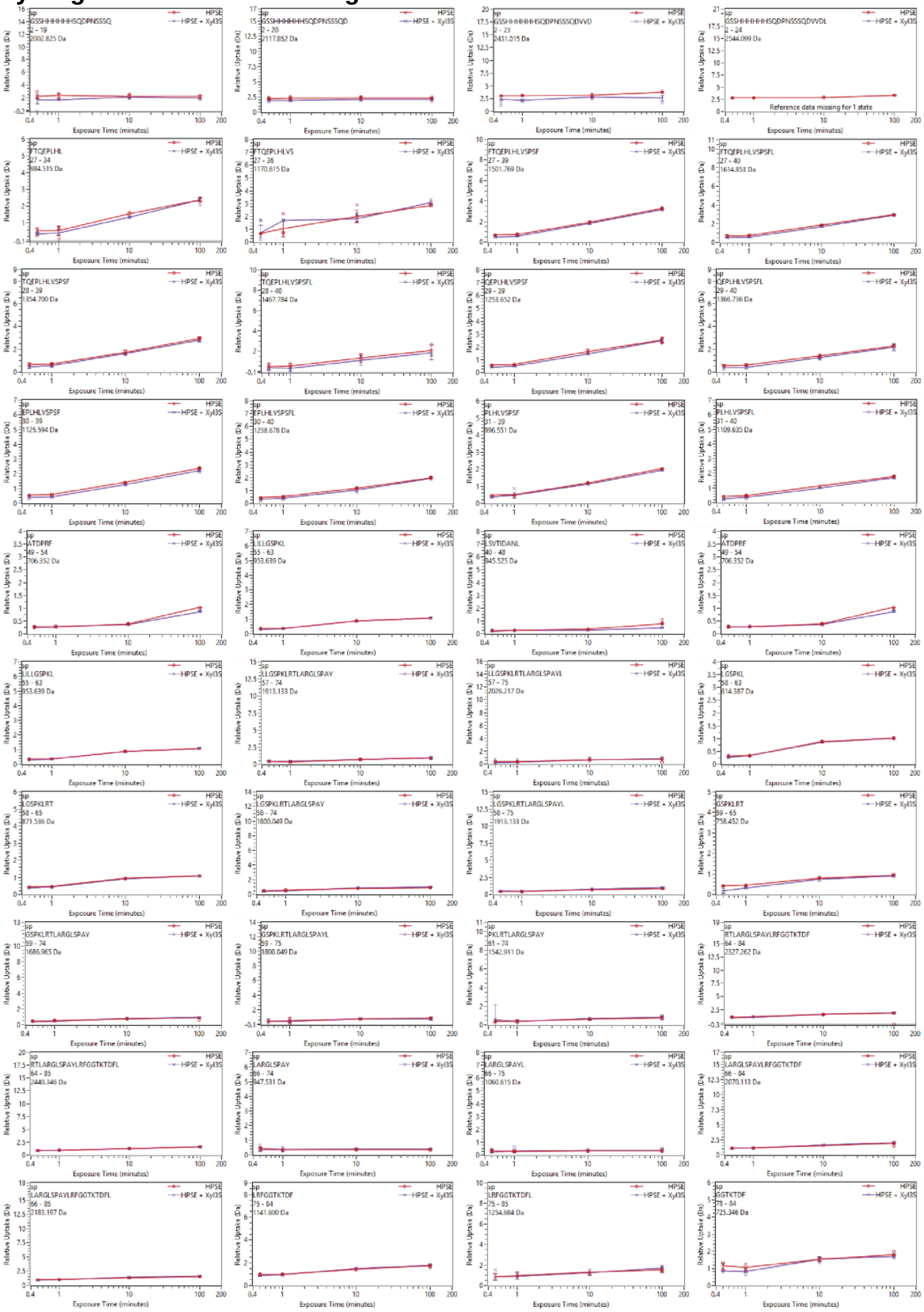

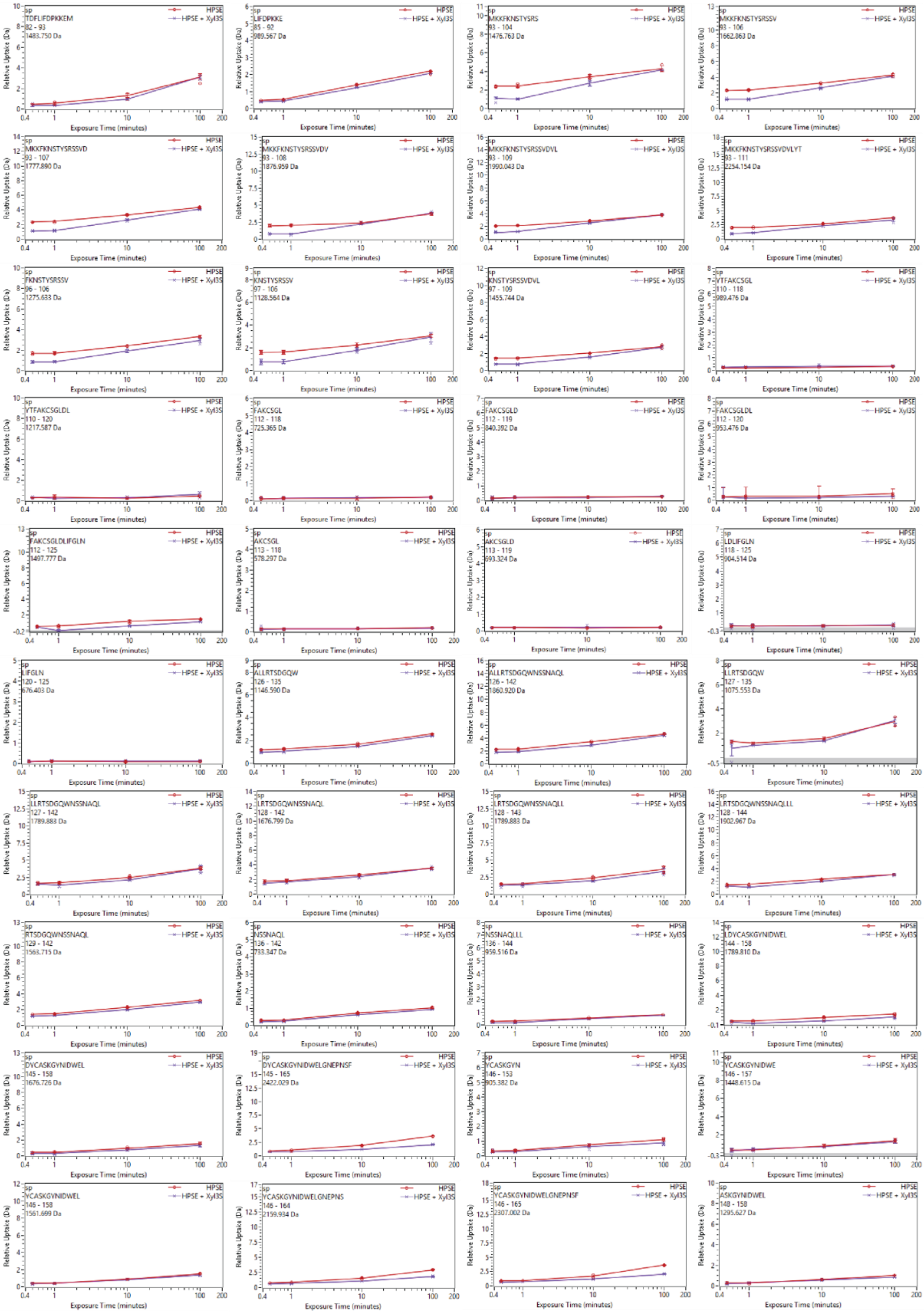
HPSE
HPSE + Xyl3S
Relative Uptake (Da)
Exposure Time (minutes)
sp TDFLIFDPKKEM 82 - 93 1483.750 Da
sp LIFDPKKE 85 - 92 989.567 Da
sp MKKFKNSTYSRS 93 - 104 1476.763 Da
sp MKKFKNSTYSRSSV 93 - 106 1662.863 Da
sp MKKFKNSTYSRSSVD 93 - 107 1777.890 Da
sp MKKFKNSTYSRSSVDV 93 - 108 1876.959 Da
sp MKKFKNSTYSRSSVDVL 93 - 109 1990.043 Da
sp MKKFKNSTYSRSSVDVLYT 93 - 111 2254.154 Da
sp FKNSTYSRSSV 96 - 106 1275.633 Da
sp KNSTYSRSSV 97 - 106 1128.564 Da
sp KNSTYSRSSVDVL 97 - 109 1455.744 Da
sp YTFAKCSGL 110 - 118 989.476 Da
sp YTFAKCSGLDL 110 - 120 1217.587 Da
sp FAKCSGL 112 - 118 725.365 Da
sp FAKCSGLD 112 - 119 840.392 Da
sp FAKCSGLDL 112 - 120 953.476 Da
sp FAKCSGLDLIFGLN 112 - 125 1497.777 Da
sp AKCSGL 113 - 118 578.297 Da
sp AKCSGLD 113 - 119 693.324 Da
sp LDLIFGLN 118 - 125 904.514 Da
sp LIFGLN 120 - 125 676.403 Da
sp ALLRTSDGQW 126 - 135 1146.590 Da
sp ALLRTSDGQWNSSNAQL 126 - 142 1860.920 Da
sp LLRTSDGQW 127 - 135 1075.553 Da
sp LLRTSDGQWNSSNAQL 127 - 142 1789.883 Da
sp LRTSDGQWNSSNAQL 128 - 142 1676.799 Da
sp LRTSDGQWNSSNAQLL 128 - 143 1789.883 Da
sp LRTSDGQWNSSNAQLLL 128 - 144 1902.967 Da
sp RTSDGQWNSSNAQL 129 - 142 1563.715 Da
sp NSSNAQL 136 - 142 733.347 Da
sp NSSNAQLLL 136 - 144 959.516 Da
sp LDYCASKGYNIDWEL 144 - 158 1789.810 Da
sp DYCASKGYNIDWEL 145 - 158 1676.726 Da
sp DYCASKGYNIDWELGNEPNSF 145 - 165 2422.029 Da
sp YCASKGYN 146 - 153 905.382 Da
sp YCASKGYNIDWE 146 - 157 1448.615 Da
sp YCASKGYNIDWEL 146 - 158 1561.699 Da
sp YCASKGYNIDWELGNEPNS 146 - 164 2159.934 Da
sp YCASKGYNIDWELGNEPNSF 146 - 165 2307.002 Da
sp ASKGYNIDWEL 148 - 158 1295.627 Da

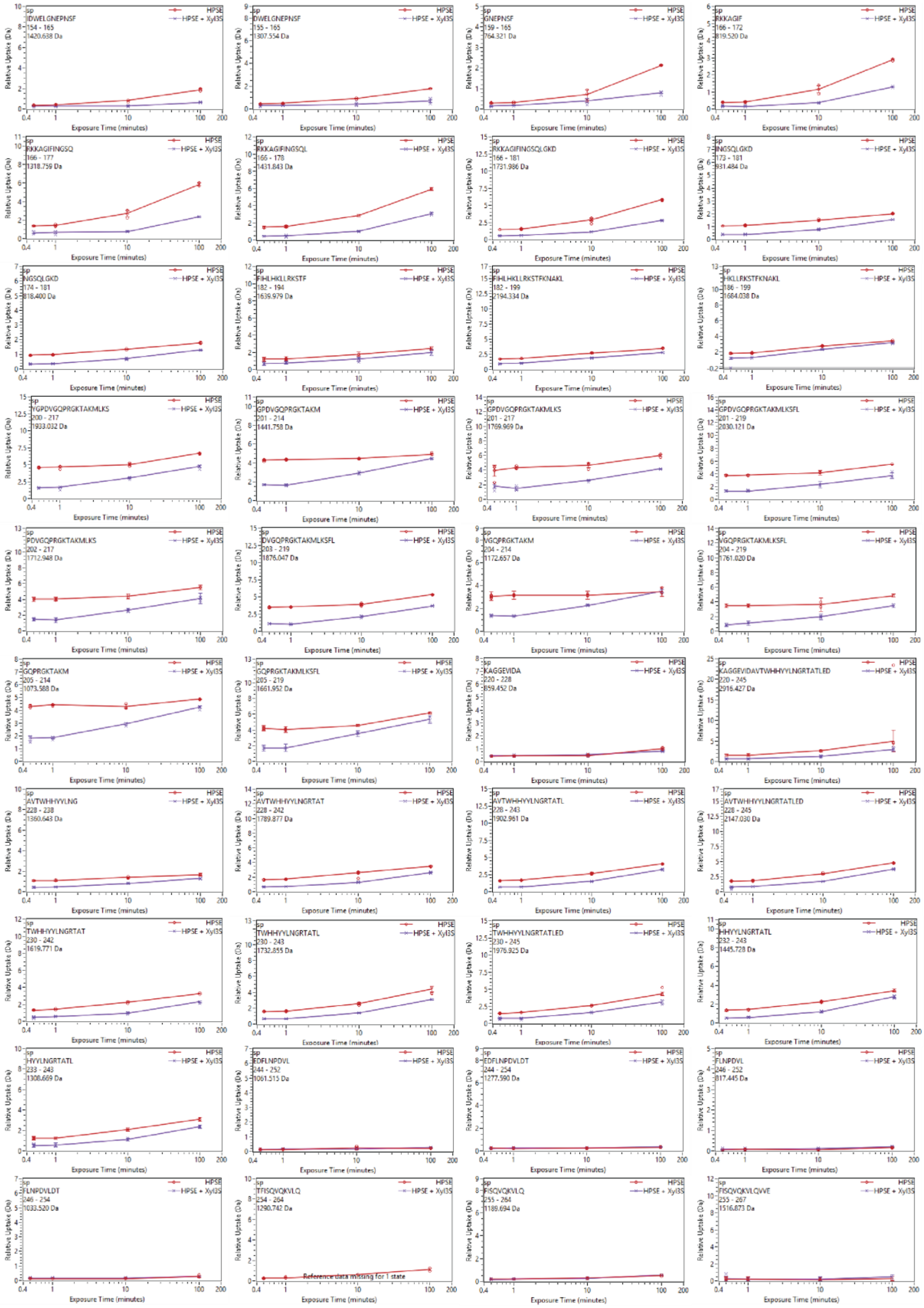
Relative Uptake (Da)
Exposure Time (minutes)
HPSE
HPSE + Xyl3S
sp IDWELGNEPNSF 154 - 165 1420.638 Da
sp DWELGNEPNSF 155 - 165 1307.554 Da
sp GNEPNSF 159 - 165 764.321 Da
sp RKKAGIF 166 - 172 819.520 Da
sp RKKAGIFNGSQ 166 - 177 1318.759 Da
sp RKKAGIFNGSQL 166 - 178 1431.843 Da
sp RKKAGIFNGSQLGKD 166 - 181 1731.986 Da
sp NGSQLGKD 173 - 181 931.484 Da
sp NGSQLGKD 174 - 181 818.400 Da
sp FIHLHKLLRKSTF 182 - 194 1639.979 Da
sp FIHLHKLLRKSTFKNAKL 182 - 199 2194.334 Da
sp HKLLRKSTFKNAKL 186 - 199 1684.038 Da
sp YGPDVGQPRGKTAKMLKS 200 - 217 1933.032 Da
sp GPDVGQPRGKTAKM 201 - 214 1441.758 Da
sp GPDVGQPRGKTAKMLKS 201 - 217 1769.969 Da
sp GPDVGQPRGKTAKMLKSFL 201 - 219 2030.121 Da
sp PDVGQPRGKTAKMLKS 202 - 217 1712.948 Da
sp DVGQPRGKTAKMLKSFL 203 - 219 1876.047 Da
sp VGQPRGKTAKM 204 - 214 1172.657 Da
sp VGQPRGKTAKMLKSFL 204 - 219 1761.020 Da
sp GQPRGKTAKM 205 - 214 1073.588 Da
sp GQPRGKTAKMLKSFL 205 - 219 1661.952 Da
sp KAGGEVIDA 220 - 228 859.452 Da
sp KAGGEVIDAVTWHHYYLNGRTATLED 220 - 245 2916.427 Da
sp AVTWHHYYLNG 228 - 238 1360.643 Da
sp AVTWHHYYLNGRTAT 228 - 242 1789.877 Da
sp AVTWHHYYLNGRTATL 228 - 243 1902.961 Da
sp AVTWHHYYLNGRTATLED 228 - 245 2147.030 Da
sp TWHHYYLNGRTAT 230 - 242 1619.771 Da
sp TWHHYYLNGRTATL 230 - 243 1732.855 Da
sp TWHHYYLNGRTATLED 230 - 245 1976.925 Da
sp HHYYLNGRTATL 232 - 243 1445.728 Da
sp HYYLNGRTATL 233 - 243 1308.669 Da
sp EDFLNPDVL 244 - 252 1061.515 Da
sp EDFLNPDVLDT 244 - 254 1277.590 Da
sp FLNPDVL 246 - 252 817.445 Da
sp FLNPDVLDT 246 - 254 1033.520 Da
sp TFISQVQKVLQ 254 - 264 1290.742 Da
Reference data missing for 1 state
sp FISQVQKVLQ 255 - 264 1189.694 Da
sp FISQVQKVLQVVE 255 - 267 1516.873 Da

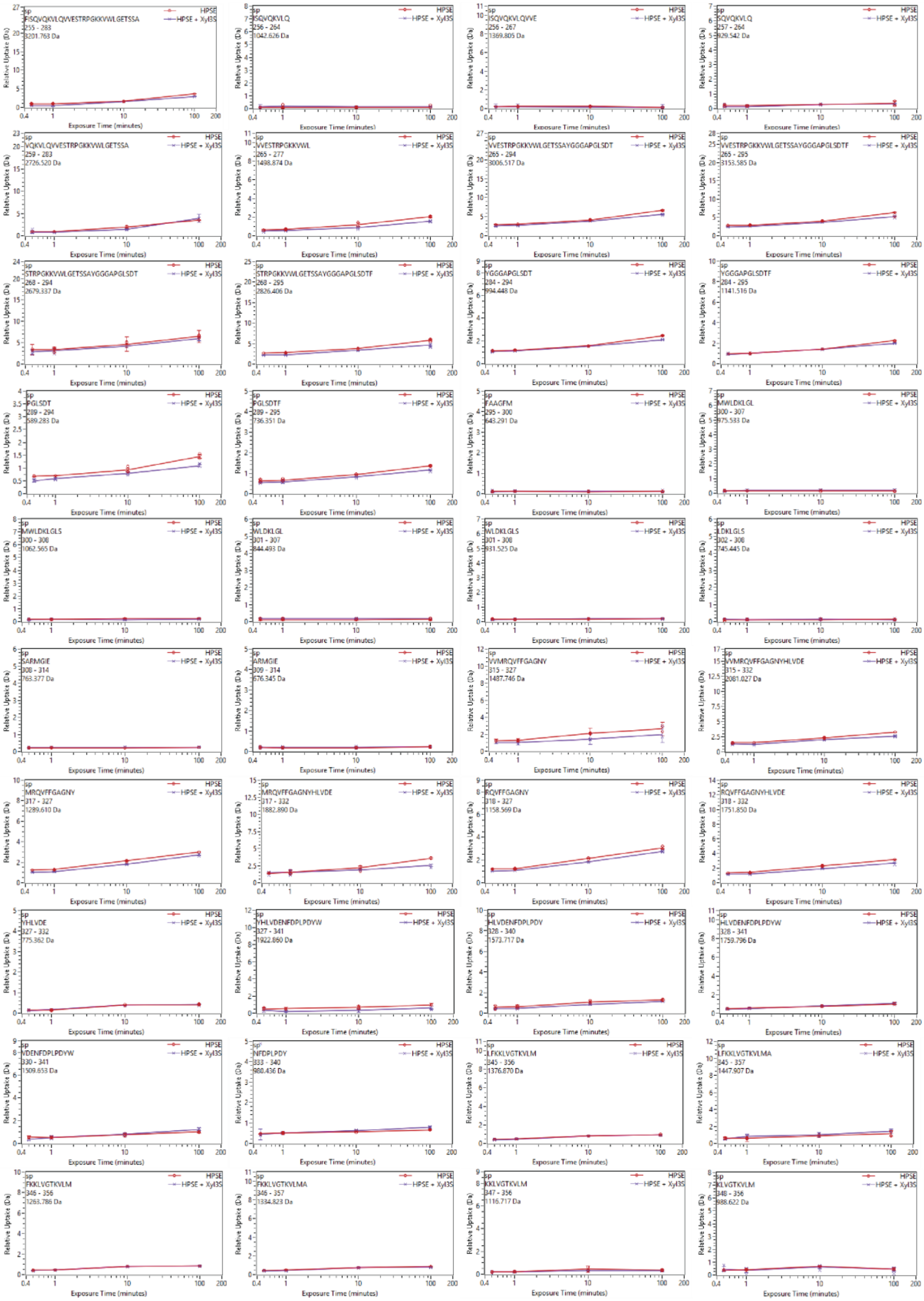
Relative Uptake (Da)
Exposure Time (minutes)
HPSE
HPSE + Xyl3S
sp
FISQVQKVLQVVESTRPGKKVWLGETSSA 255 - 283 3201.763 Da
ISQVQKVLQ 256 - 264 1042.626 Da
ISQVQKVLQVVE 256 - 267 1369.805 Da
SQVQKVLQ 257 - 264 929.542 Da
VQKVLQVVESTRPGKKVWLGETSSA 259 - 283 2726.520 Da
VVESTRPGKKVWL 265 - 277 1498.874 Da
VVESTRPGKKVWLGETSSAYGGGAPGLSDT 265 - 294 3006.517 Da
VVESTRPGKKVWLGETSSAYGGGAPGLSDTF 265 - 295 3153.585 Da
STRPGKKVWLGETSSAYGGGAPGLSDT 268 - 294 2679.337 Da
STRPGKKVWLGETSSAYGGGAPGLSDTF 268 - 295 2826.406 Da
YGGGAPGLSDT 284 - 294 994.448 Da
YGGGAPGLSDTF 284 - 295 1141.516 Da
PGLSDT 289 - 294 589.283 Da
PGLSDTF 289 - 295 736.351 Da
FAAGFM 295 - 300 643.291 Da
MWLDKLGL 300 - 307 975.533 Da
MWLDKLGLS 300 - 308 1062.565 Da
WLDKLGL 301 - 307 844.493 Da
WLDKLGLS 301 - 308 931.525 Da
LDKLGLS 302 - 308 745.445 Da
SARMGIE 308 - 314 763.377 Da
ARMGIE 309 - 314 676.345 Da
VVMRQVFFGAGNY 315 - 327 1487.746 Da
VVMRQVFFGAGNYHLVDE 315 - 332 2081.027 Da
MRQVFFGAGNY 317 - 327 1289.610 Da
MRQVFFGAGNYHLVDE 317 - 332 1882.890 Da
RQVFFGAGNY 318 - 327 1158.569 Da
RQVFFGAGNYHLVDE 318 - 332 1751.850 Da
YHLVDE 327 - 332 775.362 Da
YHLVDENFDPLPDYW 327 - 341 1922.860 Da
HLVDENFDPLPDY 328 - 340 1573.717 Da
HLVDENFDPLPDYW 328 - 341 1759.796 Da
VDENFDPLPDYW 330 - 341 1509.653 Da
NFDPLPDY 333 - 340 980.436 Da
LFKKLVGTKVLM 345 - 356 1376.870 Da
LFKKLVGTKVLMA 345 - 357 1447.907 Da
FKKLVGTKVLM 346 - 356 1263.786 Da
FKKLVGTKVLMA 346 - 357 1334.823 Da
KKLVGTKVLM 347 - 356 1116.717 Da
KLVGTKVLM 348 - 356 988.622 Da

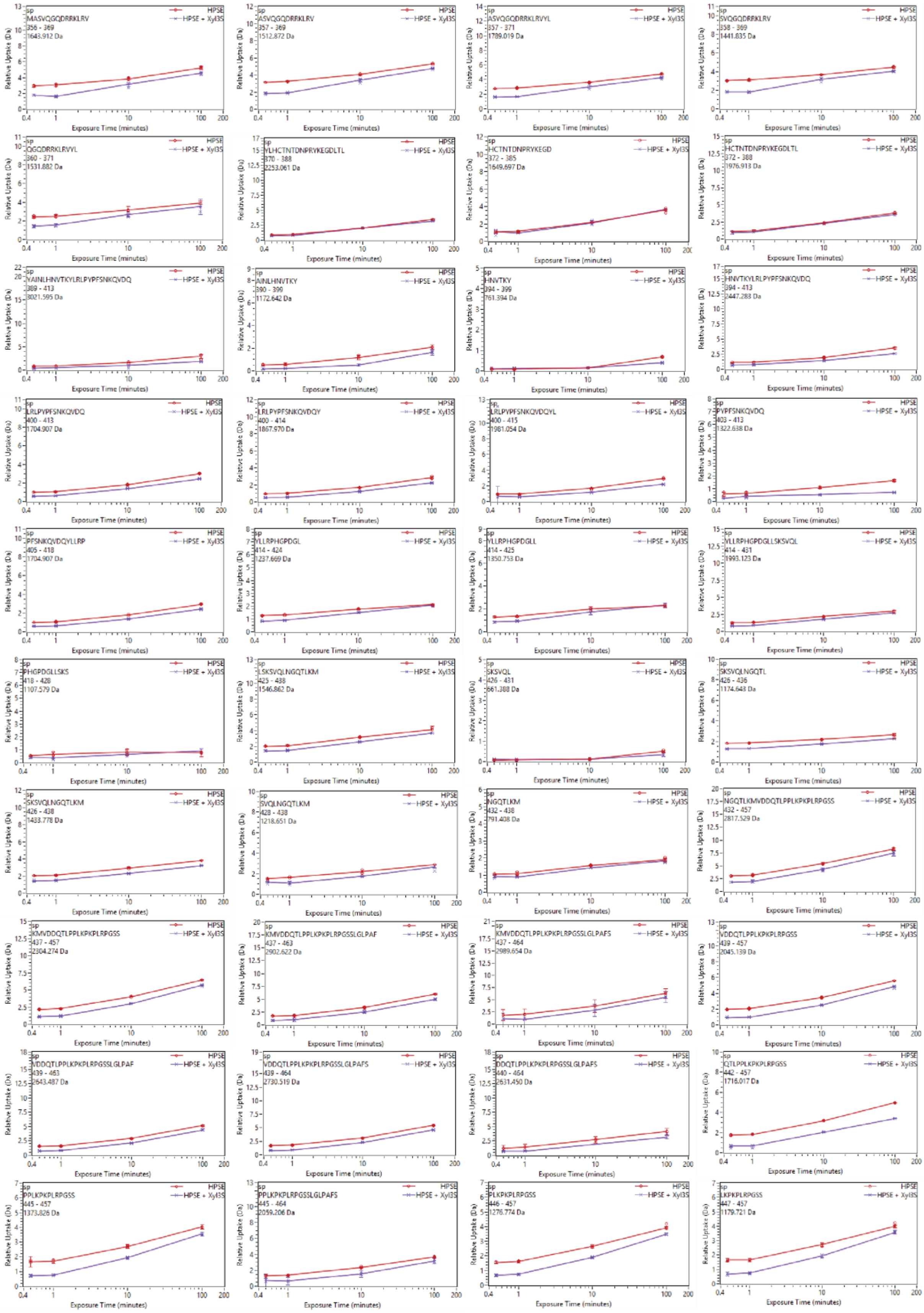

sp
MASVQGQDRRKLRV
356 - 369
1643.912 Da
ASVQGQDRRKLRV
357 - 369
1512.872 Da
ASVQGQDRRKLRVYL
357 - 371
1789.019 Da
SVQGQDRRKLRV
358 - 369
1441.835 Da
QGQDRRKLRVYL
360 - 371
1531.882 Da
YLHCTNTDNPRYKEGDLTL
370 - 388
2253.061 Da
HCTNTDNPRYKEGD
372 - 385
1649.697 Da
HCTNTDNPRYKEGDLTL
372 - 388
1976.913 Da
YAINLHNVTKYLRLPYPFSNKQVDQ
389 - 413
3021.595 Da
AINLHNVTKY
390 - 399
1172.642 Da
HNVTKY
394 - 399
761.394 Da
HNVTKYLRLPYPFSNKQVDQ
394 - 413
2447.283 Da
LRLPYPFSNKQVDQ
400 - 413
1704.907 Da
LRLPYPFSNKQVDQY
400 - 414
1867.970 Da
LRLPYPFSNKQVDQYL
400 - 415
1981.054 Da
YPFSNKQVDQ
403 - 413
1322.638 Da
PFSNKQVDQYLLRP
405 - 418
1704.907 Da
YLLRPHGPDGL
414 - 424
1237.669 Da
YLLRPHGPDGLL
414 - 425
1350.753 Da
YLLRPHGPDGLLSKSVQL
414 - 431
1993.123 Da
PHGPDGLLSKS
418 - 428
1107.579 Da
LSKSVQLNGQTLKM
425 - 438
1546.862 Da
SKSVQL
426 - 431
661.388 Da
SKSVQLNGQTL
426 - 436
1174.648 Da
SKSVQLNGQTLKM
426 - 438
1433.778 Da
SVQLNGQTLKM
428 - 438
1218.651 Da
NGQTLKM
432 - 438
791.408 Da
NGQTLKMVDDQTLPPLKPKPLRPGSS
432 - 457
2817.529 Da
KMVDDQTLPPLKPKPLRPGSS
437 - 457
2304.274 Da
KMVDDQTLPPLKPKPLRPGSSLGLPAF
437 - 463
2902.622 Da
KMVDDQTLPPLKPKPLRPGSSLGLPAFS
437 - 464
2989.654 Da
VDDQTLPPLKPKPLRPGSS
439 - 457
2045.139 Da
VDDQTLPPLKPKPLRPGSSLGLPAF
439 - 463
2643.487 Da
VDDQTLPPLKPKPLRPGSSLGLPAFS
439 - 464
2730.519 Da
DDQTLPPLKPKPLRPGSSLGLPAFS
440 - 464
2631.450 Da
QTLPPLKPKPLRPGSS
442 - 457
1716.017 Da
PPLKPKPLRPGSS
445 - 457
1373.826 Da
PPLKPKPLRPGSSLGLPAFS
445 - 464
2059.206 Da
PLKPKPLRPGSS
446 - 457
1276.774 Da
LKPKPLRPGSS
447 - 457
1179.721 Da
HPSE
HPSE + Xyl3S
Relative Uptake (Da)
Exposure Time (minutes)

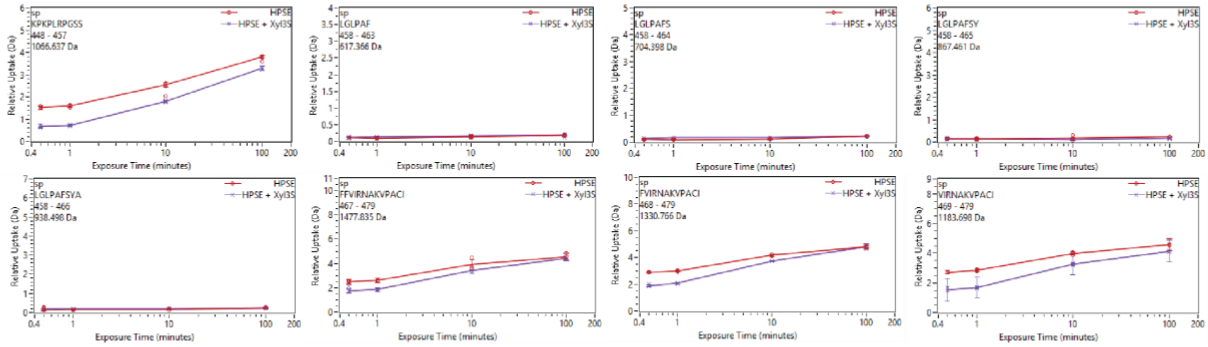


**Figure S7.** Hydrogen/deuterium exchange data of apo and $Xyl_3S$ bound HPSE. Deuterium uptake plots of the indicated peptides. Apo and $Xyl_3S$ bound HPSE are indicated by red and blue lines, respectively. Error bars represent the standard deviation from triplicate experiments.

### General Synthesis Methods

Optical rotations were measured in the solvent specified using a Rudolph Research Analytical Autopol I Automatic Polarimeter (sodium D line, 589 nm). Unless otherwise noted, proton ($^{1}H$) and carbon ($^{13}C$) NMR spectra were recorded at 18 °C in $CDCl_3$ (treated with $K_2CO_3$ and then stored over 4 Å molecular sieves) or in $D_2O$ (99.9 atom % D) on Bruker Ascend 400, 600, 700 MHz or Varian MR400 400 MHz spectrometers. Chemical shifts are recorded in parts per million (ppm) downfield from tetramethylsilane (δ TMS = 0). For $^{1}H$ NMR spectra, signals arising from the residual protio-forms of the solvent were used as the internal standards. Accordingly, residual $CHCl_3$ appearing at $\delta_H$ 7.26, and the HDO resonance at $\delta_H$ 4.79, were used to reference $^{1}H$ NMR spectra. For $^{13}C\{^{1}H\}$ NMR spectra, the central resonance of the $CDCl_3$ "triplet" appearing at $\delta_C$ 77.16 and for samples recorded in $D_2O$, ca. 1 µL of $CH_3CN$ was added and the "$CH_3$" signal arising at $\delta_C$ 1.47 was used as a reference. When required for the purposes of differentiation, signals appearing in $^{13}C$ NMR spectra are reported to two decimal places. $^{1}H$ NMR data are recorded as follows: chemical shift δ (ppm) [multiplicity, coupling constant(s) *J* (Hz), relative integral] where multiplicity is defined as: s = singlet; d = doublet; t = triplet; q = quartet; m = multiplet; b = broad, app. = apparent, obsc. = obscured, or combinations of the above. Infrared spectra were recorded on a Perkin-Elmer UATR Two spectrometer as a thin film or solid. Low-resolution mass spectrometry (LRMS) and high-resolution mass spectrometry (HRMS) were performed using positive or negative mode electrospray ionisation ($ESI^{+/-}$) techniques on a Micromass ZMD ESI-Quadrupole, a Waters LCT Premier XE, a Thermo-Fischer Scientific Orbitrap Elite™ Hybrid Ion Trap-Orbitrap or an Orbitrap QExactive mass spectrometer. A VG Fisons Auto-Spec mass spectrometer was used to obtain low- and high-resolution electron impact (EI) mass spectra. Analytical thin layer chromatography (TLC) was performed on aluminum-backed Merck silica gel 60 $F_{254}$ plates. Eluted plates were visualized using a 254 nm UV lamp, and/or charring the dried plate with a blow-torch or by treatment with a suitable dip followed by heating. These dips included phosphomolybdic acid : ceric sulfate : sulfuric acid (conc.) : water (37.5 g : 7.5 g : 37.5 g : 720 mL) or potassium permanganate : potassium carbonate : 5% sodium hydroxide aqueous solution : water (3 g : 20 g: 5 mL : 300 mL). Flash chromatographic separations were carried out following protocols defined by Still *et al.*[1] with Merck silica gel 60 (40-63 µm). THF,

$Et_2O$, and DCM were dried using a Glass Contour solvent purification system that is based upon a technology originally described by Grubbs *et al.*[2] Solvent compositions were mixed in the *v/v* ratios indicated. All solvents and reagents were used as supplied unless otherwise specified. Petroleum spirit in the boiling range of 40 – 60 °C was used. Reactions were conducted in oven-dried glassware open to the atmosphere unless otherwise specified. Desalting was performed with a BioGel P2 column (50 mm × 450 mm) and eluted with reverse osmosis purified water. The xylo-oligomeric mixture used (product number XOS-95P, food grade) was obtained from Shandong Longlive Bio-technology Co., Ltd., Qingdao, China. All sulfated compounds are sodiated but the sodium counter-ions have been omitted for clarity. Reduced pressure short-path vacuum distillation, when utilised, was conducted using a still-head connected to a receiving flask *via* a vacuum receiver adapter. Xylotetraose was purified on a Waters Alliance 2695 separation module coupled with an ACQUITY TQD detector via Waters flow splitter 15:1. MassLynx version 4.1 Software was used for data acquisition and processing.

### Specific Transformations

For the preparation of $Xyl_3S$ and $Xyl_8S$ see Malins et al.[3]

*Preparation of $Xyl_4S$*

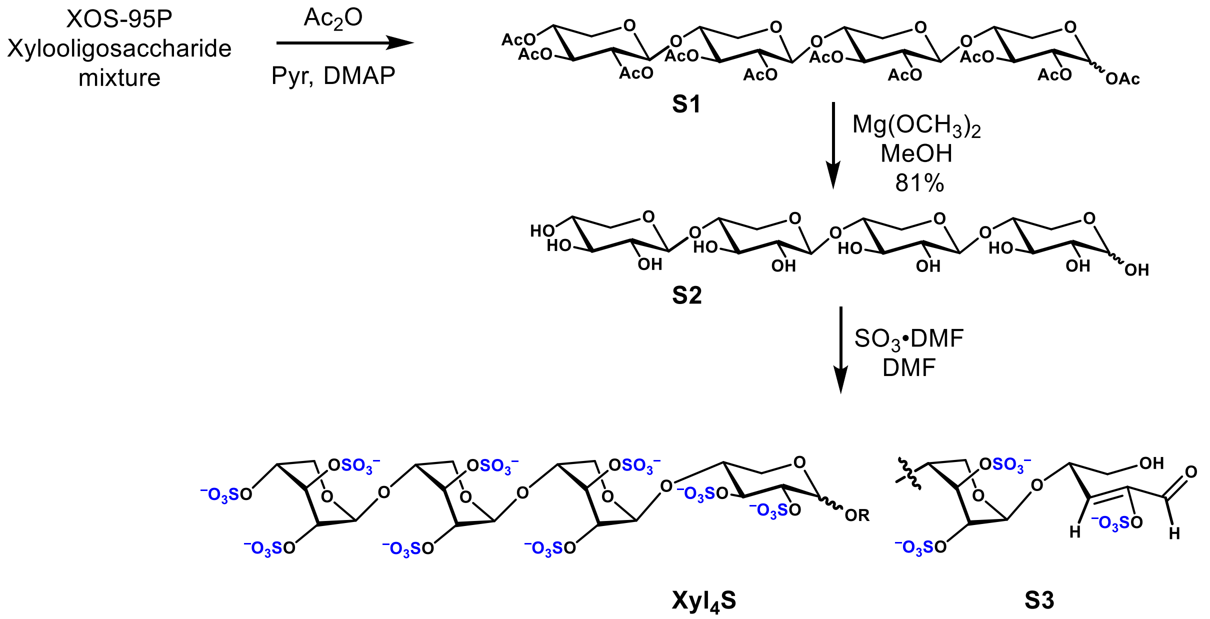

*Deca-O-acetyl-D-xylotetraose (**S1**)*

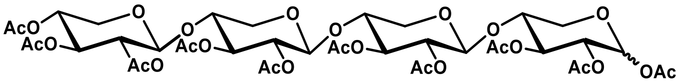


Following a procedure analogous to that reported by Malins[3] and Kosma,[4] a mixture of xylooligosaccharides (8.0 g, XOS-95P food-grade material) in anhydrous pyridine (40 mL), and maintained under an atmosphere of nitrogen, was magnetically stirred and after ca. 1 h a clear solution was obtained. Thereafter, the solution was cooled in an ice-water bath (0 – 5 °C) and treated sequentially with 4-dimethylaminopyridine (DMAP) (50 mg) and acetic anhydride (100 mL). Stirring was continued and the reaction was warmed to 18 °C over 24 h then cooled in an ice-water bath (0 – 5 °C) and treated cautiously with methanol (200 mL), portion-wise, over 0.5 h. After 1 h the mixture was co-evaporated with toluene (3 × 100 mL) and so affording a viscous pale-yellow syrup that was subjected to flash column chromatography (silica, 1:1 → 3:2 → 7:3 *v/v* EtOAc/pet. spirit gradient elution) and so affording three fractions, A, B and C.

Concentration of fraction A (**$R_f$** = 0.4 in 4:3 *v/v* EtOAc/pet. spirit) afforded hexa-*O*-acetyl-D-xylobiose (4.90 g) as a colorless foam. See Malins[3] and Kosma[4] for further details and spectral data.

Concentration of fraction B (**$R_f$** = 0.3 in 4:3 *v/v* EtOAc/pet. spirit) afforded octa-*O*-acetyl-D-xylotriose (3.80 g) as a colorless foam. See Malins[3] and Kosma[4] for further details and spectral data.

Concentration of fraction C (**$R_f$** = 0.2 in 4:3 *v/v* EtOAc/pet. spirit) afforded deca-*O*-acetyl-D-xylotetraose (**S1**) (1.20 g) as a colorless foam.

**$[\alpha]_D$** = −81.0 (*c* = 1.0, $CDCl_3$)

**$^1$H NMR** (400 MHz, $CDCl_3$) *1.0β : 0.7α mixture of C-1 anomers* δ 6.18 (d, *J* = 3.7 Hz, 1H), 5.62 (d, *J* = 7.2 Hz, 1H), 5.35 (dd, *J* = 10.3, 8.8 Hz, 1H), 5.11 (app. t, *J* = 8.4 Hz, 1H), 5.09 – 5.00 (m, 6H), 4.95 – 4.91 (m, 2H), 4.88 – 4.83 (m, 2H), 4.79 – 4.76 (m, 2H), 4.74 – 4.70 (m, 4H), 4.54 – 4.52 (m, 2H), 4.47 – 4.44 (m, 4H), 4.09 – 4.05 (m, 2H), 3.97 (dd, *J* = 11.9, 5.1 Hz, 1H), 3.94 – 3.89 (m, 4H), 3.77 – 3.72 (m, 7H), 3.63 (app. t, *J* = 10.5 Hz, 1H), 3.44 (dd, *J* = 12.0, 9.1 Hz, 1H), 3.37 (dd, *J* = 12.0, 7.7 Hz, 2H), 3.33 – 3.27 (m, 4H), 2.15 – 1.99 (20 × $CH_3$, 60H).

**$^{13}$C NMR** (101 MHz, $CDCl_3$) *1.0β : 0.7α mixture of C-1 anomers, many overlapping signals; all observed signals are reported here* δ 170.04, 169.99, 169.94, 169.92, 169.91, 169.83, 169.71, 169.66, 169.48, 169.44, 169.27, 169.25, 169.12, 100.78, 100.37, 100.34, 100.32, 99.56, 92.26, 89.31, 75.80, 74.78, 74.74, 74.72, 74.23, 72.04, 71.99, 71.93, 71.92, 70.99, 70.87, 70.51, 70.38, 70.05, 69.85, 69.52, 68.37, 63.47, 62.64, 62.55, 61.62, 61.43, 21.00, 20.89, 20.87, 20.86, 20.84, 20.76, 20.73, 20.70, 20.61.
**LRMS** (ESI+): *m/z* (rel. int.) 989 (100) $[M+ Na]^+$.
**HRMS** (ESI+) found 989.2755, calcd for $C_{40}H_{54}NaO_{27}^+$ $[M+ Na]^+$ 989.2745.
**IR** $v_{max}$ = 1748, 1368, 1216, 1043 $cm^{-1}$.

*Xylotetraose (**S2**)*

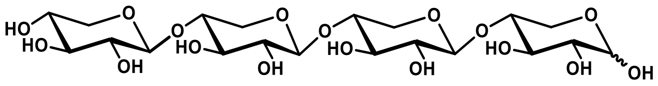


A magnetically stirred solution of deca-*O*-acetyl-D-xylotetraose (**S1**) (300 mg, 0.31 mmol) in anhydrous methanol (10 mL) was treated, at 18 °C, with a solution of magnesium methoxide in methanol (8.20 mL, 4.65 mmol, 6-10 wt. %). The resulting mixture was stirred for 48 h then quenched with Dowex 50WX4-100 ($H^+$-form) to pH 2. The resin was filtered off and washed with methanol and the combined filtrates then concentrated under reduced pressure to afford xylotetraose (**S2**) (137 mg, 81%) as an amorphous white powder that was used without further purification. A portion of the ensuing product (ca. 20 mg) was subjected to purification by HPLC (see details below) for characterisation purposes which provided, after concentration of the appropriate fractions, xylotetraose, as an amorphous white powder.
**$[\alpha]_D$** = −53.9 ($c$ = 0.5, $H_2O$), Lit[5] −59.1 ($c$ = 1.0, $H_2O$)
**$^1$H NMR** (400 MHz, $D_2O$) *1.0β : 0.55α mixture of C-1 anomers* δ 5.20 (d, $J$ = 3.7 Hz, 1H), 4.60 (d, $J$ = 7.9 Hz, 1H), 4.51 – 4.46 (m, 6H), 4.12 (dd, $J$ = 11.8, 5.3 Hz, 4H), 4.07 (dd, $J$ = 11.8, 5.4 Hz, 1H), 3.98 (dd, $J$ = 11.6, 5.5 Hz, 2H), 3.85 – 3.74 (m, 8H), 3.67 – 3.60 (m, 3H), 3.60 – 3.53 (m, 6H), 3.47 – 3.34 (m, 8H), 3.34 – 3.23 (m, 8H).
**$^{13}$C NMR** (101 MHz, $D_2O$) *1.0 β : 0.55 α mixture of C-1 anomers, many overlapping signals; all observed signals are reported here* δ 102.38, 102.20, 97.04, 92.56, 77.09, 76.93, 76.90, 76.87, 76.14, 74.53, 74.44, 74.20, 74.17, 73.31, 73.23, 71.92, 71.48, 69.72, 65.75, 63.51, 63.49, 59.36.

**LRMS** (ESI+): *m/z* (rel. int.) 569 (100) $[M+ Na]^+$.

**HRMS** (ESI+) found 569.1686 calcd for $C_{20}H_{34}NaO_{17}^+$ $[M+ Na]^+$ 569.1688.

HPLC purification of Xylotetraose: TR = 8.38 minutes (Column: Waters Carbohydrate Analysis, 3.9 x 300 mm (Part# WAT084038). Solvent A – Water + 0.1% (v/v) formic acid; Solvent B – Acetonitrile + 0.1% (v/v) formic acid. Solvent gradient: Solvent B 80% (0 min), 50% (from 0 to 7 min), 50% (from 7 to 10 min). Flow: 1 mL/min. Column temperature: 30 °C. Mass scanned from 200 to 800 (*m/z*); Ionisation mode ES-; Capillary voltage 3.60 kV; Cone voltage 45 V; Source temperature 150 °C; Desolvation Temperature 450 °C: SIR channel was also set at 545.4658 (*m/z*).

*Sulfation of xylotetraose with $SO_3$•DMF, Preparation of* ***$Xyl_4S$***

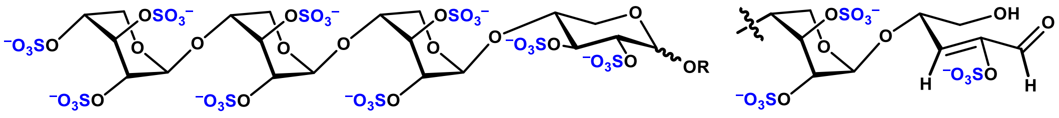


R = H or $SO_3^-$ $Xyl_4S$ **S3**

A magnetically stirred solution of $SO_3$•DMF (448 mg, 2.93 mmol) in DMF (1.5 mL) maintained under an atmosphere of nitrogen was cooled to –20 °C (using a $CO_2$(s)/EtOH/$H_2O$-based cryogenic system) then treated dropwise with a solution of xylotetraose (**S2**) (80 mg, 146 μmol) in DMF (2 mL). After 0.7 h the mixture was warmed to 0 °C (ice-bath) and stirring continued for 0.5 h before the reaction was cautiously quenched with aqueous sodium carbonate (10% w/v) until effervescence had ceased (pH ~7 – 8). The mixture was concentrated under reduced pressure by short-path distillation (1 mmHg, water bath at 40 °C) to remove volatiles and afford, as the residue, a white powder. The residue was desalted by passing it through a BioGel P2 column and fractions were analyzed by spotting on a TLC plate and charring. The appropriate fractions were collected and water removed by freeze-drying. The powder thus obtained was dissolved in water (ca. 2 mL) and subjected to ion-exchange with Dowex 50WX4-100 ($Na^+$-form) followed by freeze-drying to afford the desired product(s) (**$Xyl_4S$**) (77 mg) as a white amorphous powder. At the time of $^{13}C$ NMR data acquisition, **$Xyl_4S$** contained a mixture of R = H : R = $SO_3$ : $\Delta Xyl_{red}$(CHO) (**S3**) of 11.0 : 7.7 : 2.3. The spectral data and decomposition/hydrolysis profiles are in accord with those reported for sulfated xylooligosaccharides incorporating an unprotected reducing terminus.[3,6,7]

**$^1$H NMR** (400 MHz, $D_2O$) *mixture of C-1 anomers* δ 9.38 (s, 1H, $\Delta Xyl_{red}$(CHO)), 6.76 (d, *J* = 8.8 Hz, 1H, $\Delta Xyl_{red}$(CHO)), 5.93 (d, *J* = 3.4 Hz, 1H, α-$Xyl_{red}$-$OSO_3$), 5.77 (app. s, 1H, β-$Xyl_{red}$-$OSO_3$), 5.36 (d, *J* = 2.9 Hz, 1H, α-$Xyl_{red}$), 5.24 (app. s, 1H, β-$Xyl_{red}$), 5.18 – 5.13 (m), 5.10 – 5.08 (m), 4.84 – 4.82 (m), 4.79 – 4.78 (m), 4.77 – 4.70 (m), 4.57 – 4.54 (m), 4.54 – 4.42 (m), 4.42 – 4.38 (m), 4.35 (dd, *J* = 7.6, 2.9 Hz), 4.00 – 3.77 (m).

**$^{13}$C NMR** (101 MHz, $D_2O$) *many overlapping signals; all observed signals are reported here* δ 190.91, 147.52, 138.71, 99.74, 98.93, 98.31, 98.12, 96.07, 92.98, 91.26, 88.70, 76.23, 75.96, 75.52, 75.22, 74.94, 74.81, 74.73, 74.66, 74.43, 73.94, 73.31, 73.13, 73.07, 72.98, 72.84, 72.77, 72.39, 71.87, 71.63, 61.54, 61.36, 59.83, 59.73, 59.45, 58.92.

**LRMS** (ESI−): *m/z* 465 $[M - 3Na]^{3-}$ (R=H), $[M - 3Na]^{3-}$ (R=$SO_3$)

**HRMS** (ESI+) found 1486.6209, calcd for $C_{20}H_{25}Na_{10}O_{44}S_9^+$ (R=H) $[M+ Na]^+$ 1486.6177.

**HRMS** (ESI+) found 1588.5597, calcd for $C_{20}H_{24}Na_{11}O_{47}S_{10}^+$ (R=$SO_3$) $[M+ Na]^+$ 1588.5564.

### References


(1) Still, W. C.; Kahn, M.; Mitra, A. Rapid Chromatographic Technique for Preparative Separations with Moderate Resolution. *J. Org. Chem.* **1978**, *43* (14), 2923–2925. https://doi.org/10.1021/jo00408a041.

(2) Pangborn, A. B.; Giardello, M. A.; Grubbs, R. H.; Rosen, R. K.; Timmers, F. J. Safe and Convenient Procedure for Solvent Purification. *Organometallics* **1996**, *15* (5), 1518–1520. https://doi.org/10.1021/om9503712.

(3) Vo, Y.; Schwartz, B. D.; Onagi, H.; Ward, J. S.; Gardiner, M. G.; Banwell, M. G.; Nelms, K.; Malins, L. R. A Rapid and Mild Sulfation Strategy Reveals Conformational Preferences in Therapeutically Relevant Sulfated Xylooligosaccharides. *Chem. Eur. J.* **2021**, *27* (38), 9830–9838. https://doi.org/10.1002/chem.202100527.

(4) Abad-Romero, B.; Mereiter, K.; Sixta, H.; Hofinger, A.; Kosma, P. Synthesis of Regioselectively Sulfated Xylodextrins and Crystal Structure of Sodium Methyl β-d-Xylopyranoside 4-O-Sulfate Hemihydrate. *Carbohydr. Res.* **2009**, *344* (1), 21–28. https://doi.org/10.1016/j.carres.2008.09.018.

(5) Takeo, K.; Ohguchi, Y.; Hasegawa, R.; Kitamura, S. Synthesis of (1 → 4)-β-d-Xylo-Oligosaccharides of Dp 4–10 by a Blockwise Approach. *Carbohydr. Res.* **1995**, *278* (2), 301–313. https://doi.org/10.1016/0008-6215(95)00259-6.

(6) Itzstein, M. V.; Chang, C.-W. Sulfation Method. WO2019113646A1, June 20, 2019.

(7) Alekseeva, A.; Raman, R.; Eisele, G.; Clark, T.; Fisher, A.; Lee, S. (Larry); Jiang, X.; Torri, G.; Sasisekharan, R.; Bertini, S. In-Depth Structural Characterization of Pentosan Polysulfate Sodium Complex Drug Using Orthogonal Analytical Tools. *Carbohydr. Polym.* **2020**, *234*, 115913. https://doi.org/10.1016/j.carbpol.2020.115913.

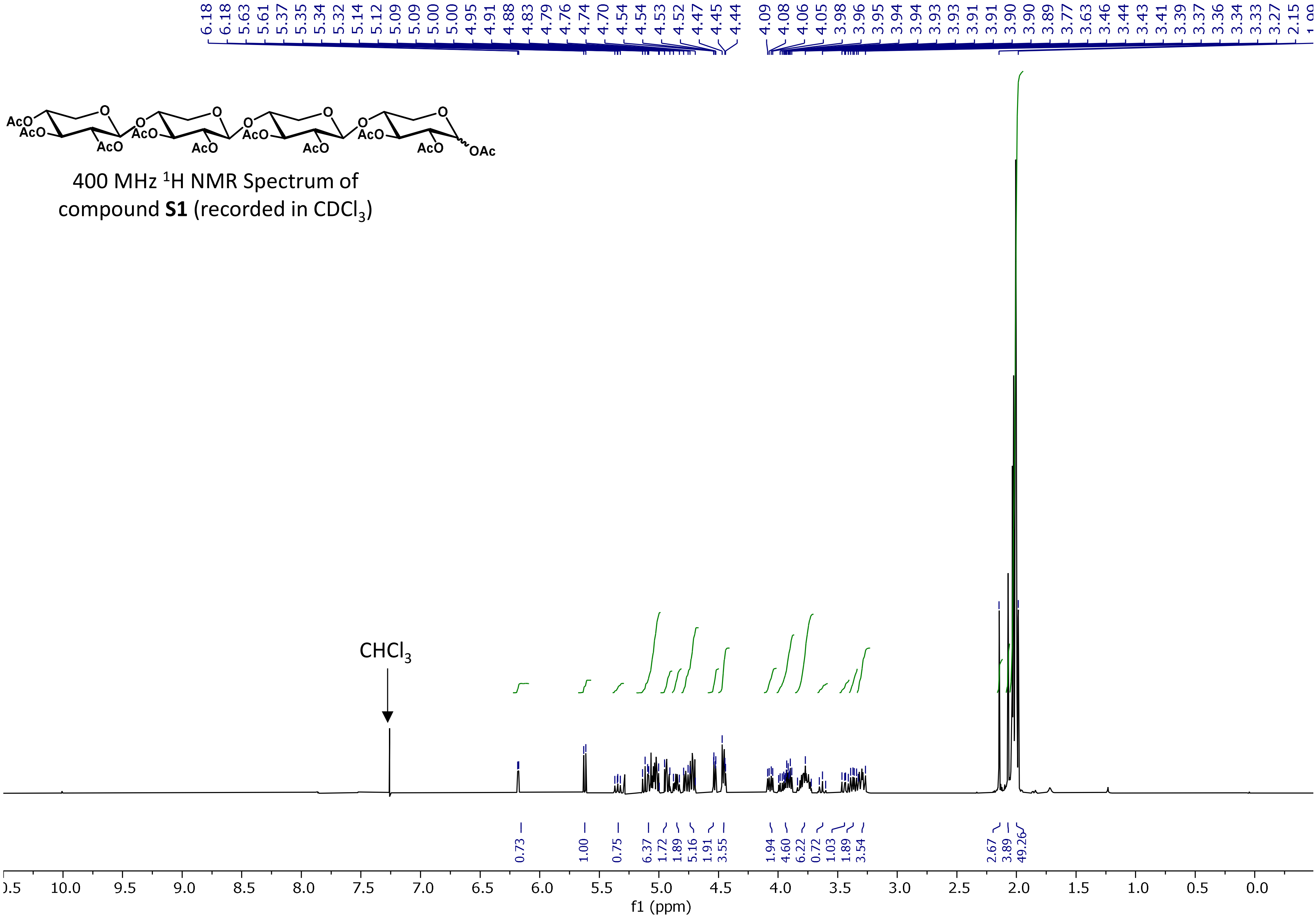
AcO
AcO
AcO
O
AcO
AcO
O
AcO
AcO
O
AcO
AcO
O
OAc
400 MHz ¹H NMR Spectrum of
compound S1 (recorded in CDCl3)
CHCl3
6.18 6.18 5.63 5.61 5.37 5.35 5.34 5.32 5.14 5.12 5.09 5.09 5.00 5.00 4.95 4.91 4.88 4.83 4.79 4.76 4.74 4.70 4.54 4.54 4.53 4.52 4.47 4.45 4.44
4.09 4.08 4.06 4.05 3.98 3.96 3.95 3.94 3.94 3.93 3.93 3.91 3.91 3.90 3.90 3.89 3.77 3.63 3.46 3.44 3.43 3.41 3.39 3.37 3.36 3.34 3.33 3.27 2.15
0.73 1.00 0.75 6.37 1.72 1.89 5.16 1.91 3.55 1.94 4.60 6.22 0.72 1.03 1.89 3.54 2.67 3.89 49.26
10.0 9.5 9.0 8.5 8.0 7.5 7.0 6.5 6.0 5.5 5.0 4.5 4.0 3.5 3.0 2.5 2.0 1.5 1.0 0.5 0.0
f1 (ppm)

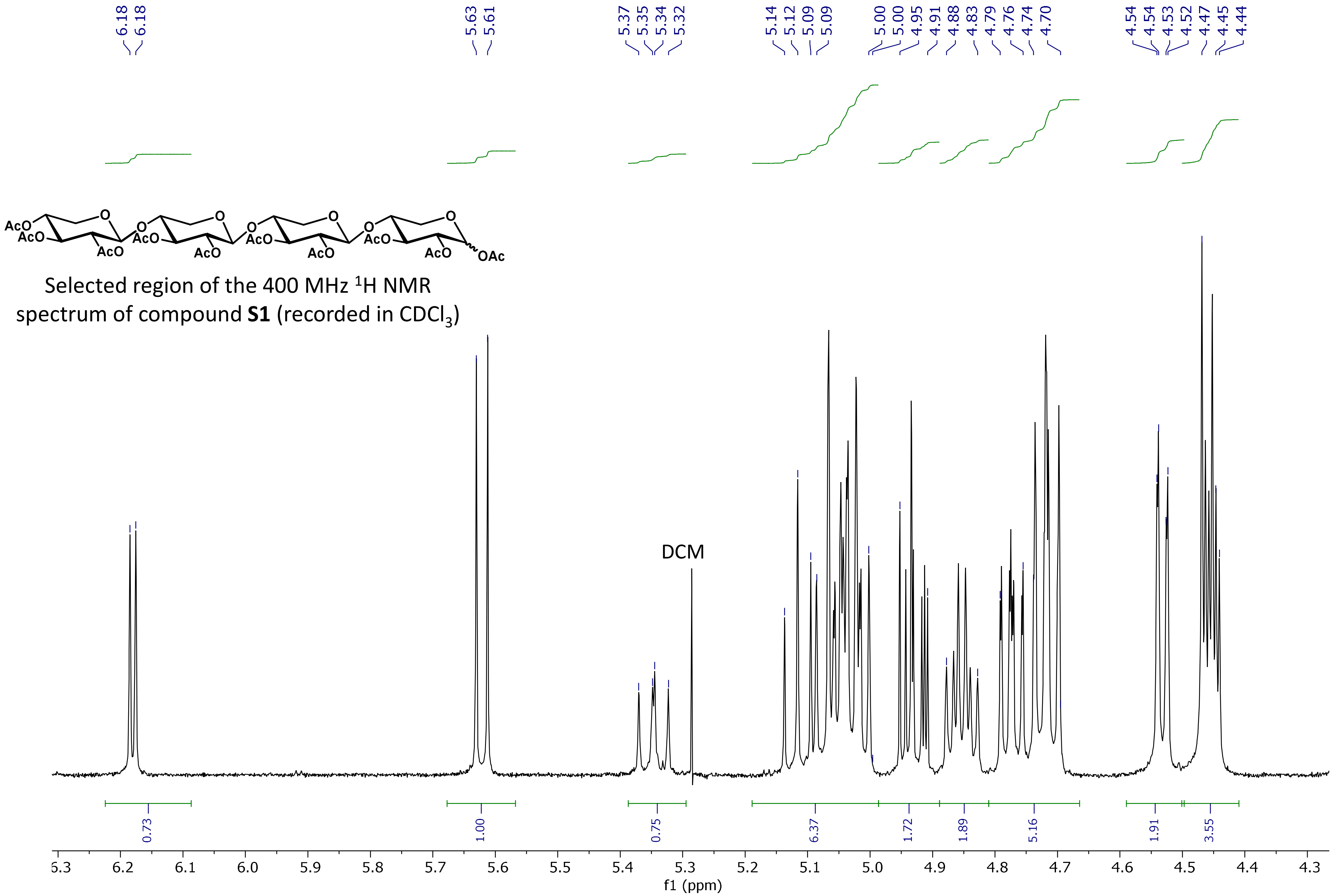

6.18
6.18
5.63
5.61
5.37
5.35
5.34
5.32
5.14
5.12
5.09
5.09
5.00
5.00
4.95
4.91
4.88
4.83
4.79
4.76
4.74
4.70
4.54
4.54
4.53
4.52
4.47
4.45
4.44
AcO
AcO
AcO
O
O
AcO
AcO
O
O
AcO
AcO
O
O
AcO
AcO
O
OAc
Selected region of the 400 MHz ¹H NMR spectrum of compound **S1** (recorded in $CDCl_3$)
DCM
0.73
1.00
0.75
6.37
1.72
1.89
5.16
1.91
3.55
6.3
6.2
6.1
6.0
5.9
5.8
5.7
5.6
5.5
5.4
5.3
5.2
5.1
5.0
4.9
4.8
4.7
4.6
4.5
4.4
4.3
f1 (ppm)

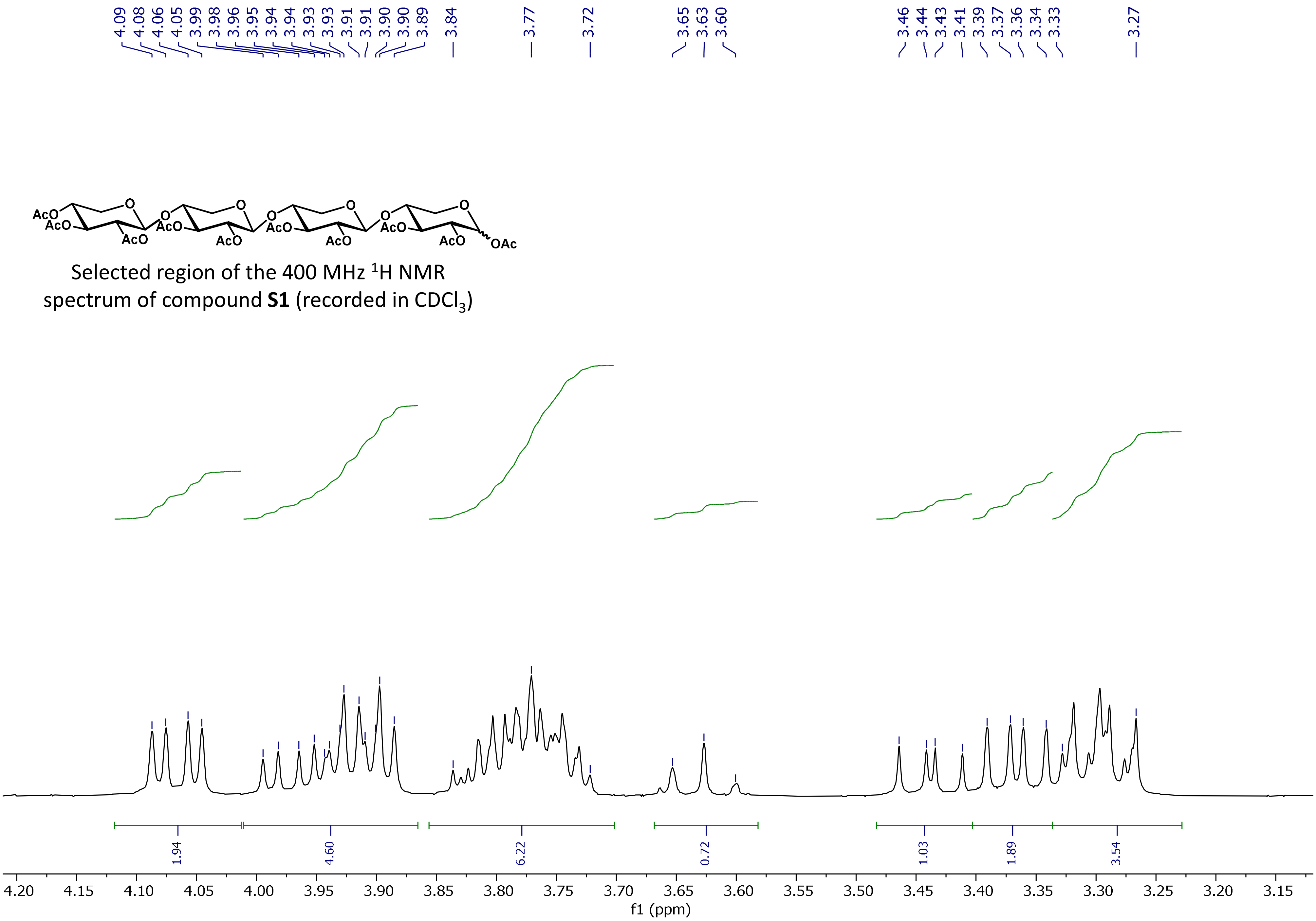


Selected region of the 400 MHz $^{1}$H NMR spectrum of compound **S1** (recorded in $CDCl_3$)

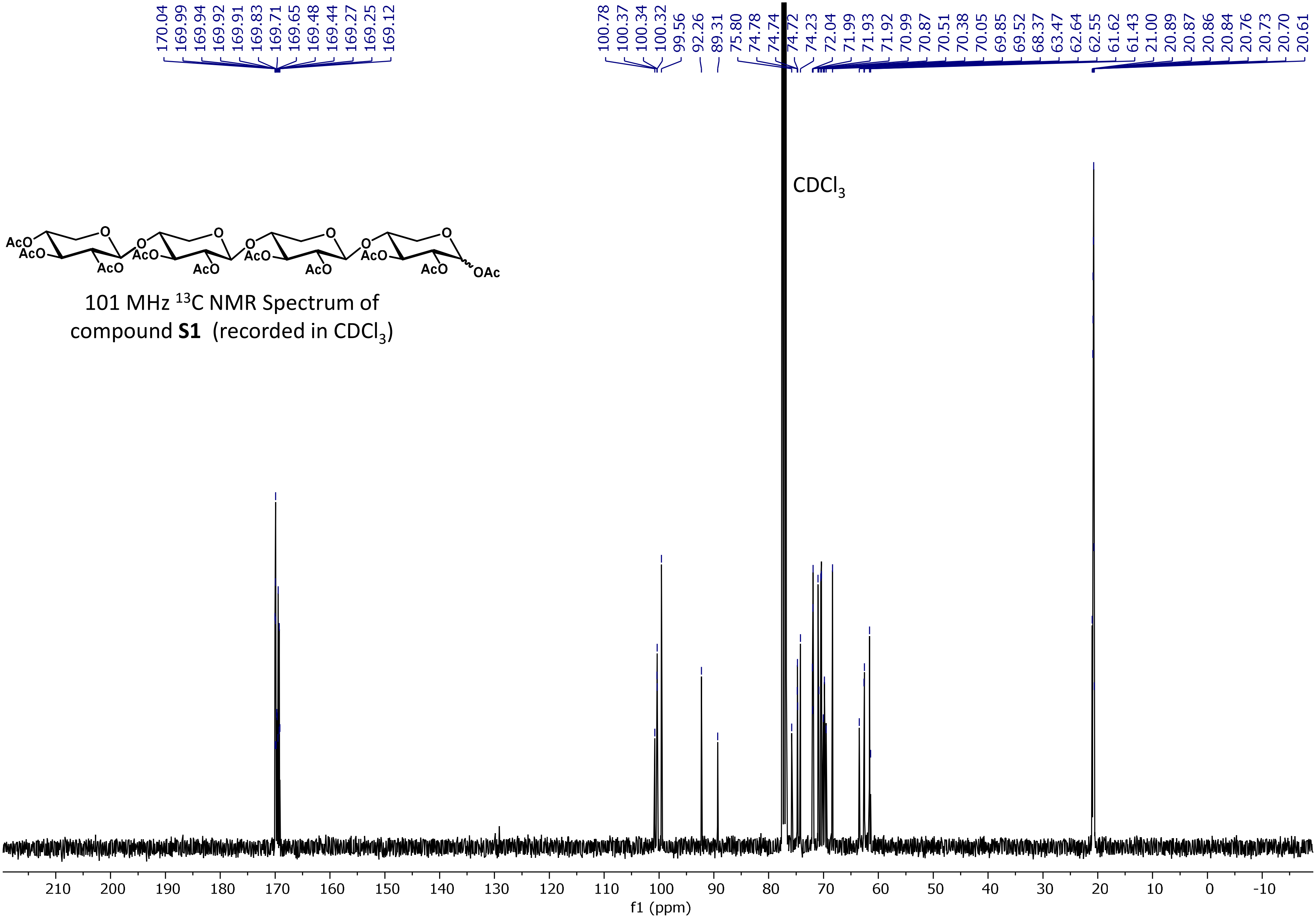


101 MHz $^{13}C$ NMR Spectrum of compound **S1** (recorded in $CDCl_3$)

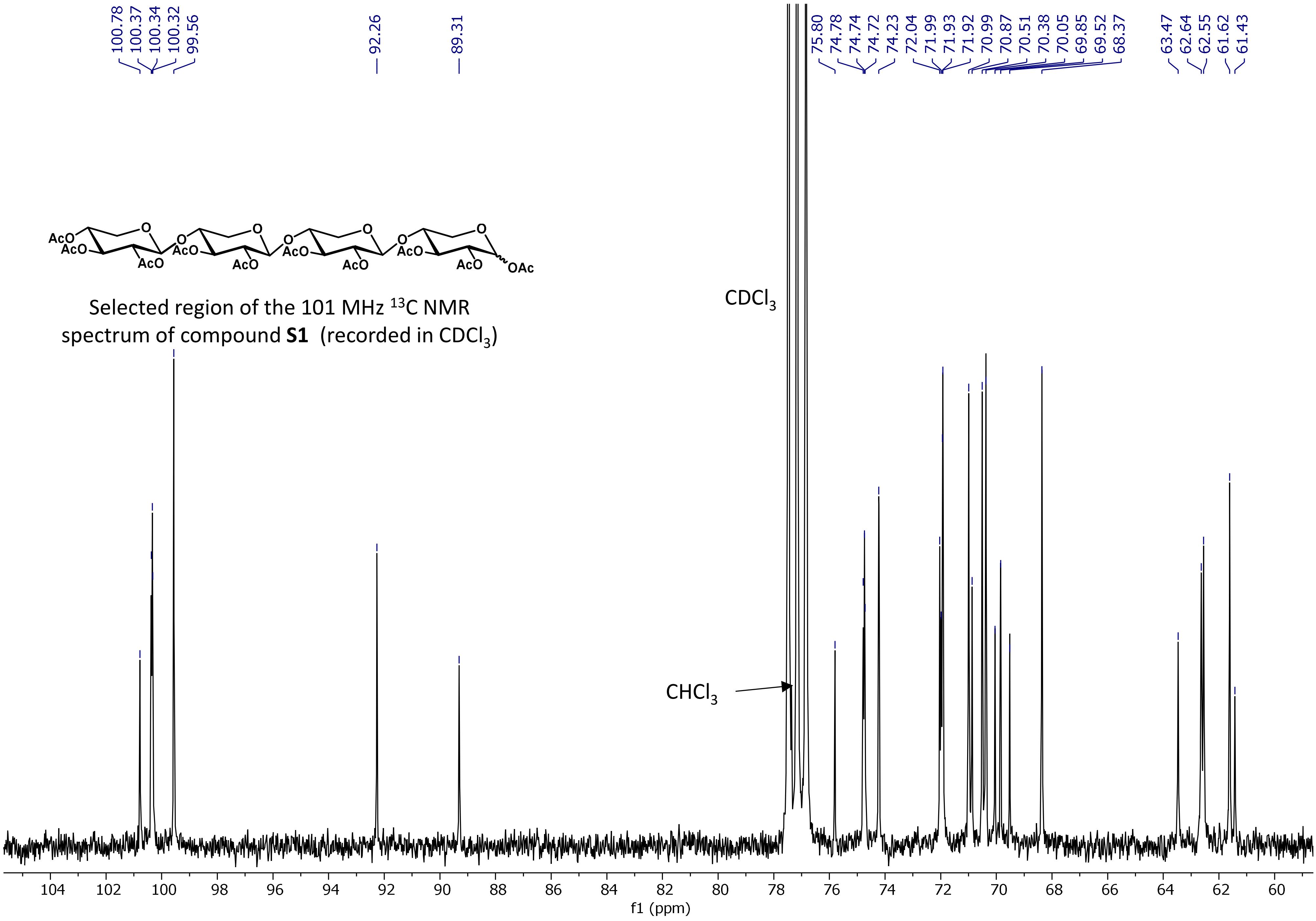


Selected region of the 101 MHz $^{13}C$ NMR spectrum of compound **S1** (recorded in $CDCl_3$)

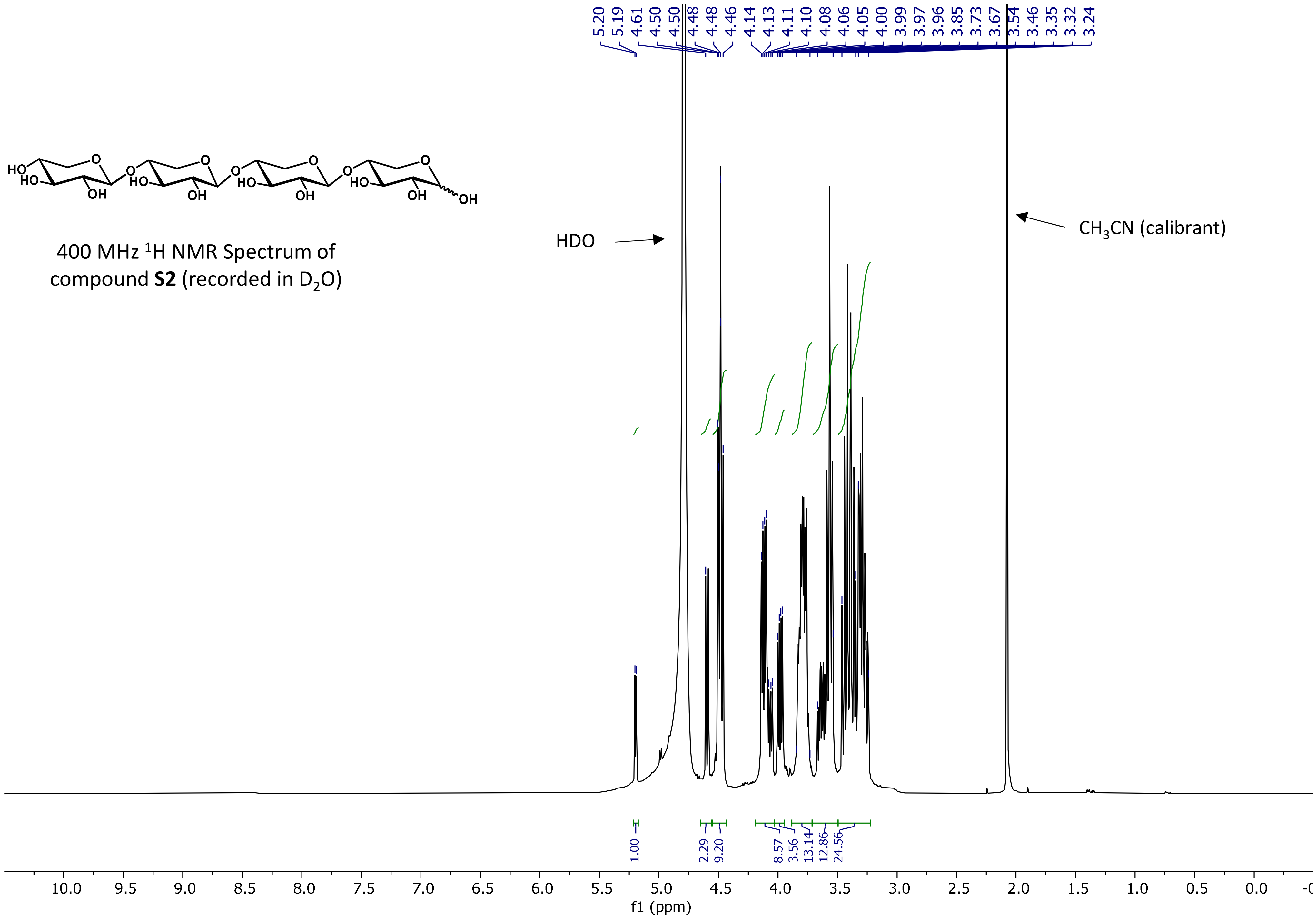
400 MHz ¹H NMR Spectrum of compound S2 (recorded in D₂O)
HDO
CH₃CN (calibrant)
5.20 5.19 4.61 4.50 4.50 4.48 4.48 4.46 4.14 4.13 4.11 4.10 4.08 4.06 4.05 4.00 3.99 3.97 3.96 3.85 3.73 3.67 3.54 3.46 3.35 3.32 3.24
1.00 2.29 9.20 8.57 3.56 13.14 12.86 24.56
f1 (ppm)

Selected region of the 400 MHz $^{1}H$ NMR spectrum of compound **S2** (recorded in $D_2O$)

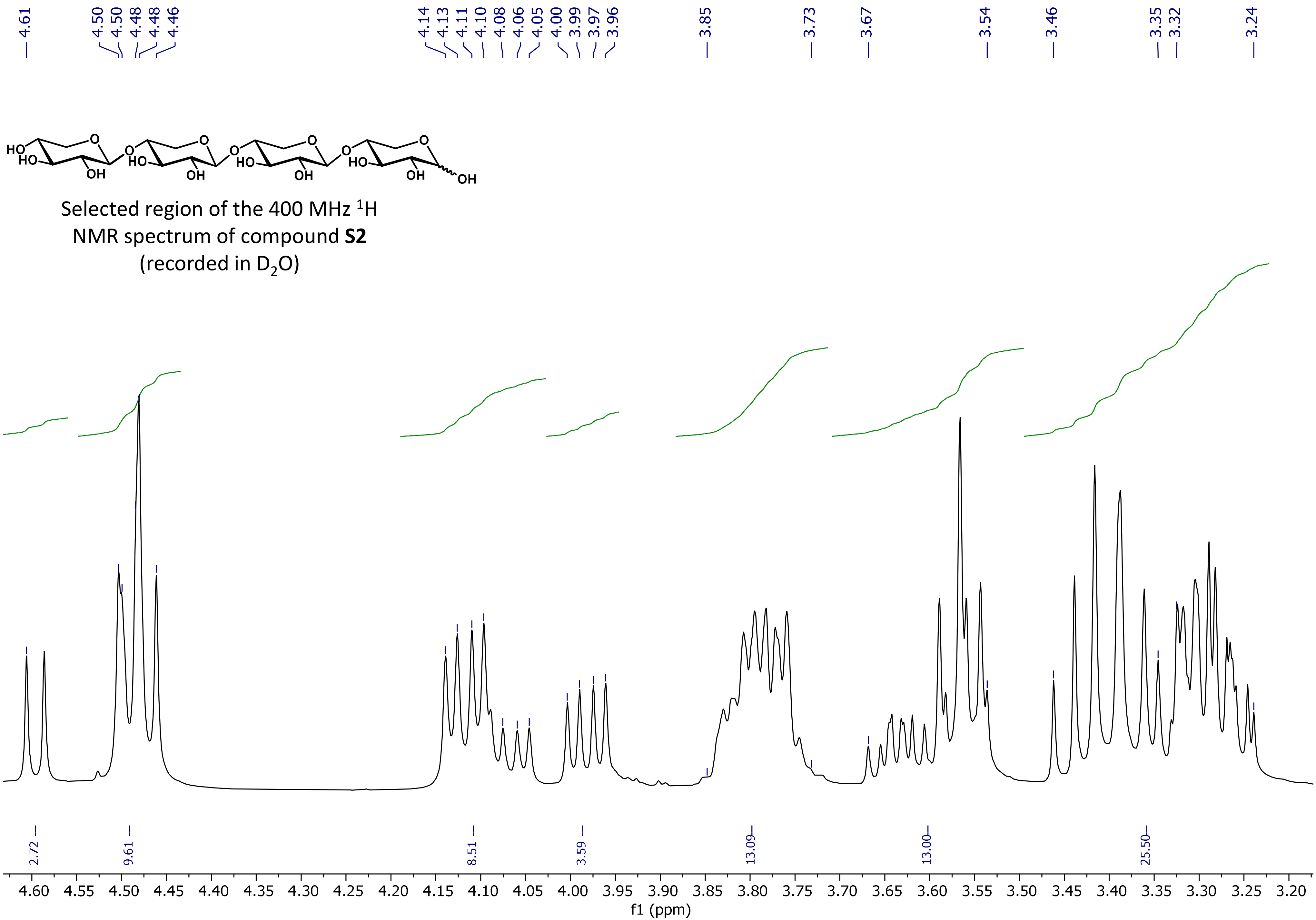

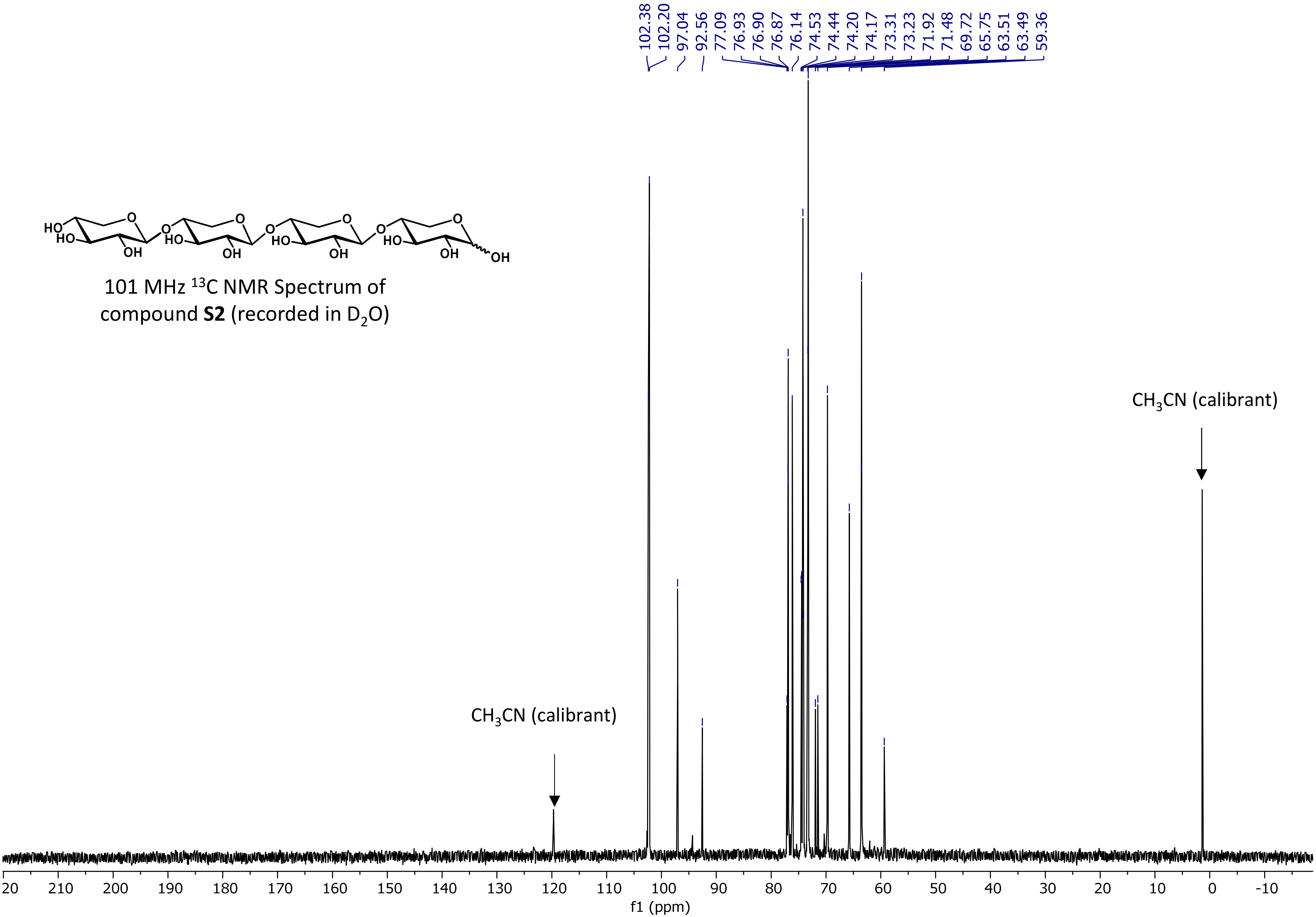


101 MHz $^{13}C$ NMR Spectrum of compound **S2** (recorded in $D_2O$)

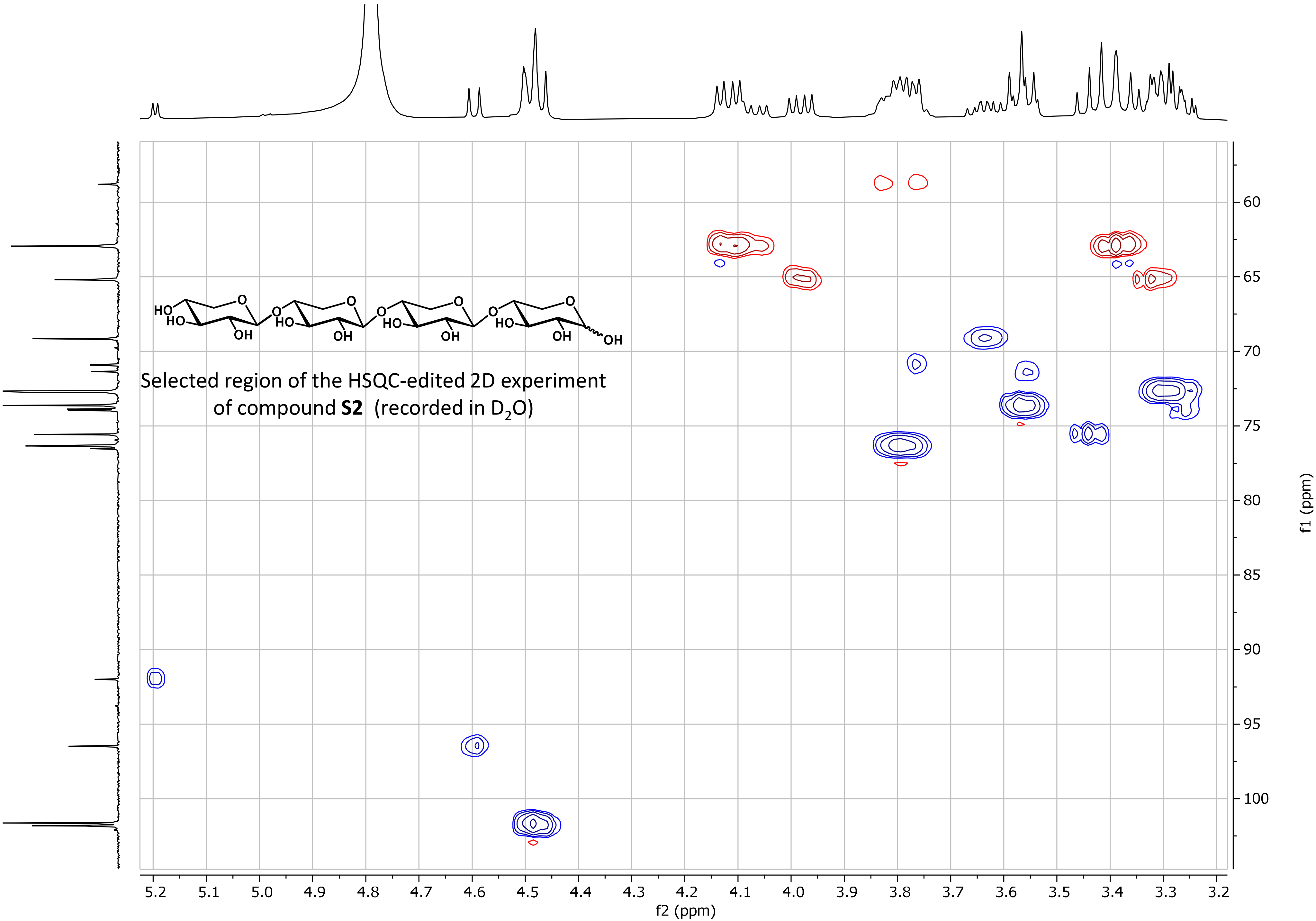


Selected region of the HSQC-edited 2D experiment of compound **S2** (recorded in $D_2O$)

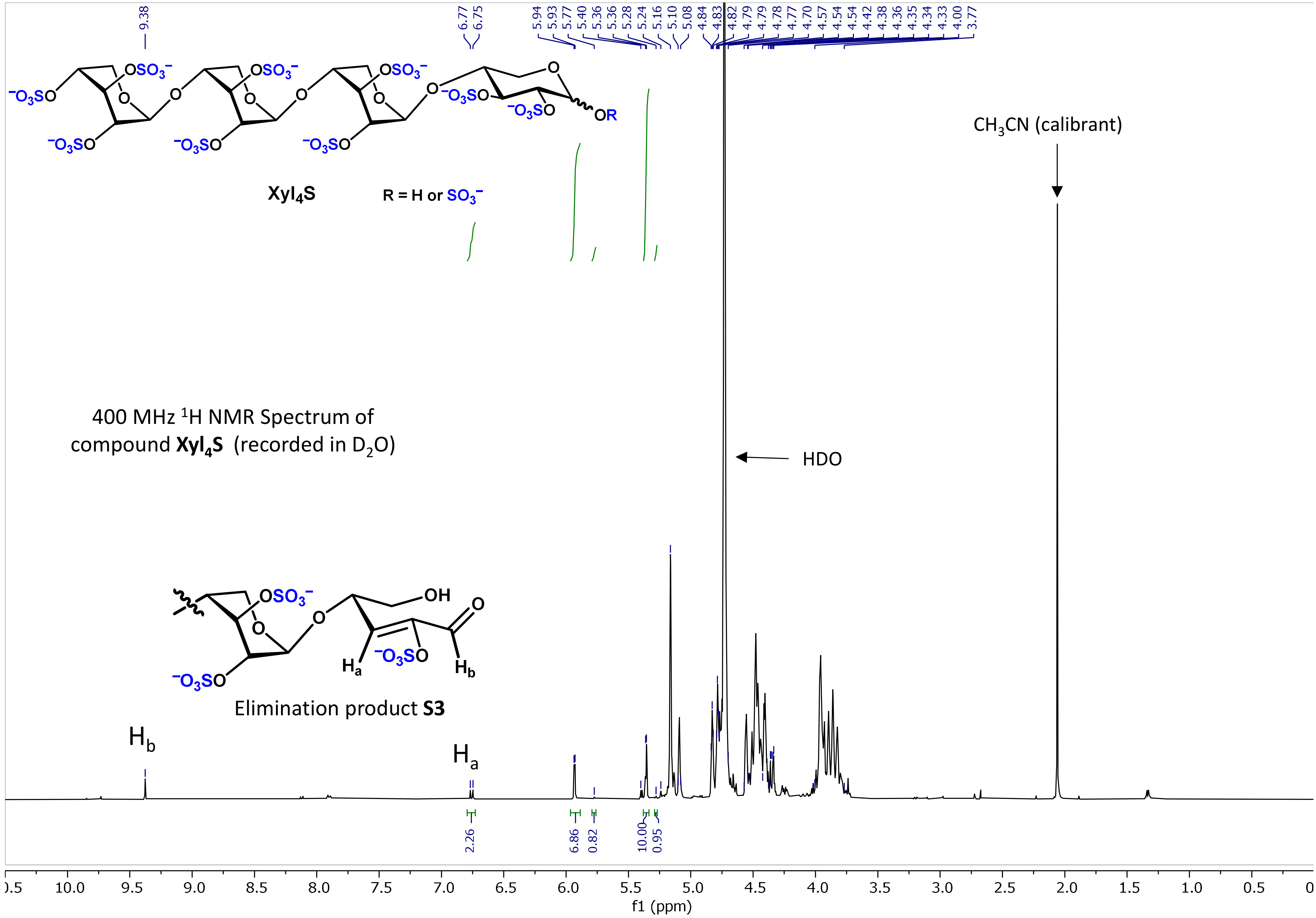
9.38
6.77
6.75
5.94
5.93
5.77
5.40
5.36
5.36
5.28
5.24
5.16
5.10
5.08
4.84
4.83
4.82
4.79
4.79
4.78
4.77
4.70
4.57
4.54
4.54
4.42
4.38
4.36
4.35
4.34
4.33
4.00
3.77
Xyl4S
R = H or SO3−
CH3CN (calibrant)
400 MHz 1H NMR Spectrum of
compound Xyl4S (recorded in D2O)
HDO
Elimination product S3
Hb
Ha
2.26
6.86
0.82
10.00
0.95
f1 (ppm)

Selected region of the 400 MHz $^1$H NMR spectrum of compound(s) **Xyl$_4$S** (recorded in $D_2O$)

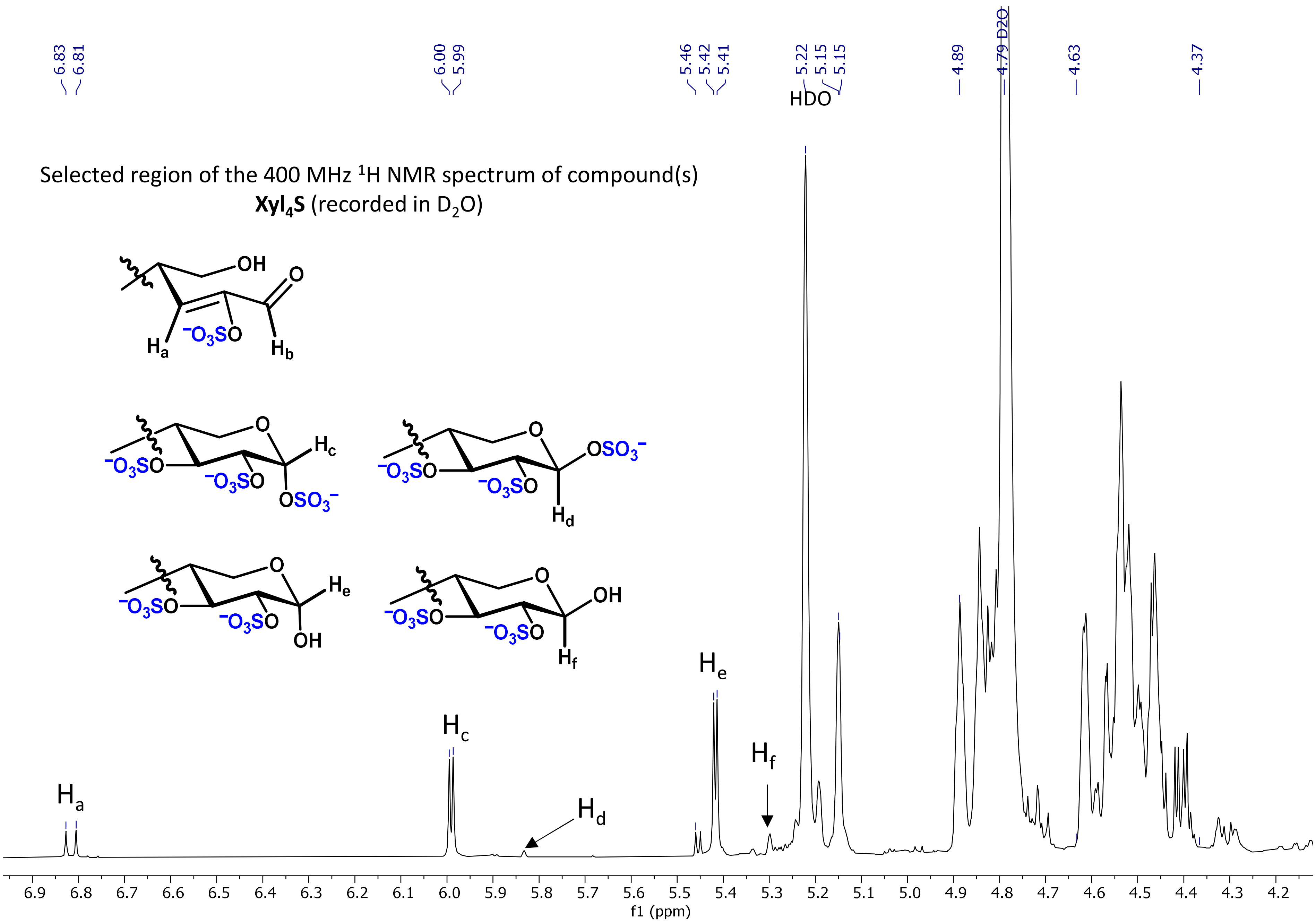

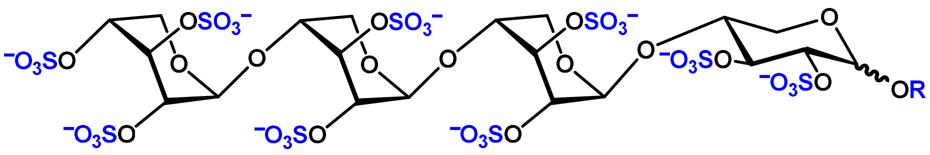

**R = H or $SO_3^-$ Xyl$_4$S**

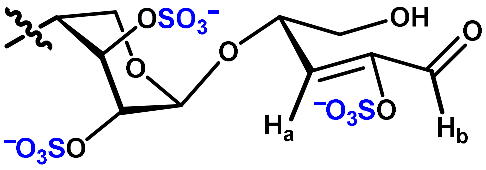

101 MHz $^{13}$C NMR Spectrum of compound **Xyl$_4$S** (recorded in $D_2O$)

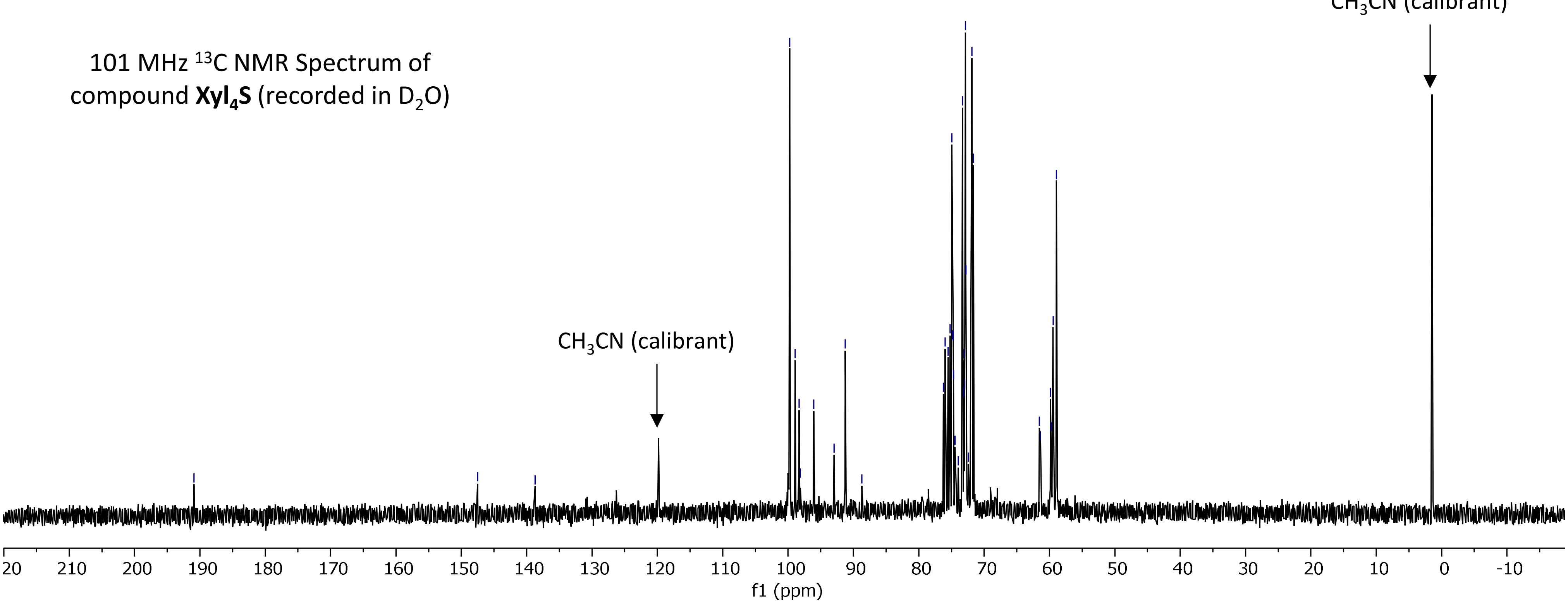

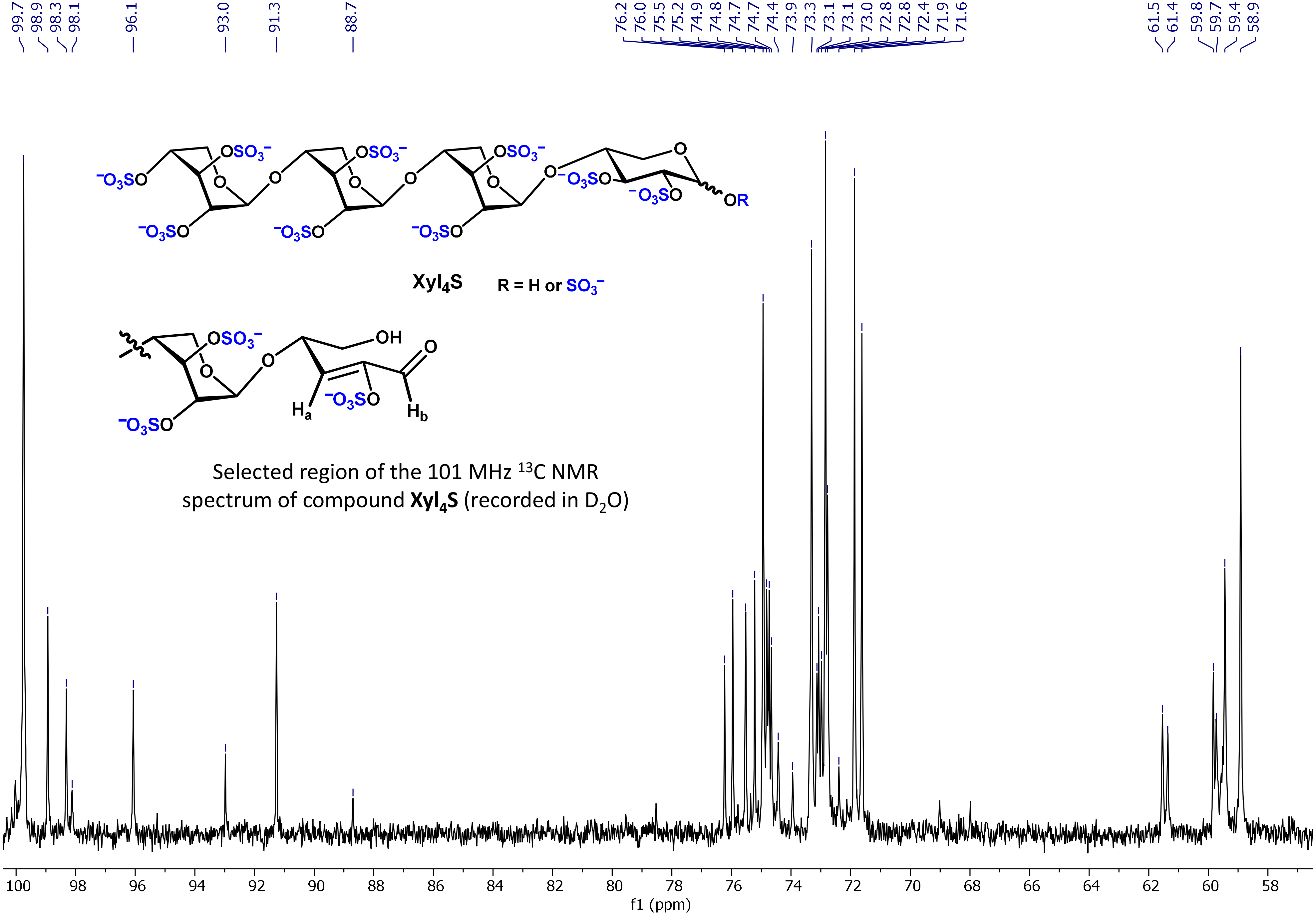


Selected region of the 101 MHz $^{13}C$ NMR spectrum of compound **Xyl$_4$S** (recorded in $D_2O$)
